# Asymmetric leading vs. trailing edge shifts since the Last Glacial Maximum underpin the modern bimodal latitudinal diversity gradient in planktonic foraminifera

**DOI:** 10.64898/2025.12.16.694627

**Authors:** Tasnuva Ming Khan, Marina C. Rillo, Lukas Jonkers, Isaiah E. Smith, Ádám T. Kocsis, Wolfgang Kiessling

## Abstract

**Aim:** The modern latitudinal diversity gradient of planktonic foraminifera is bimodal with a distinct depression near the equator, surrounded by mid-latitude diversity peaks. This pattern emerged after the Last Glacial Maximum, but it is unclear how species’ spatial dynamics contributed to its formation. Here, we investigate how species range dynamics, i.e., trailing-edge contractions (extirpations) and leading-edge expansions (colonisations), shaped the modern bimodal pattern, and how global biodiversity patterns arise from local patterns.

**Location:** Global open ocean, with basin-specific analyses in the Atlantic and Pacific Oceans.

**Time period:** Last Glacial Maximum (19 - 23 ka) (LGM) and the Pre-Industrial (modern)

**Major taxa studied:** Planktonic foraminifera (unicellular eukaryotes)

**Methods:** We analysed taxonomically standardized LGM and modern foraminiferal assemblage datasets to characterize changes in species richness at multiple spatial scales: global ocean, basin-wide, and within basin. We quantified species’ range shifts by comparing their trailing- and leading-edge movements. We estimated temporal turnover locally, and the net imbalance between colonisations and extirpations (NICE) within sites, and tested whether species’ thermal preferences correlate with their extirpation risk.

**Results:** We found no evidence of systematic trailing edge contractions, indicating that equatorial extirpations did not drive the bimodal LDG pattern. In the Atlantic, leading-edge expansions generated a coherent increase in species richness in the mid-latitudes, whereas the Pacific exhibited highly spatially heterogeneous responses, including extirpation hotspots in the western tropical Pacific and colonisation zones in the eastern and southern Pacific. Species’ thermal optima weakly predicted extirpations, with species adapted to lower temperatures more at risk of extirpation, consistent with the general warming trend since the last ice age.

**Main conclusions:** The modern bimodal LDG of planktonic foraminifera arises primarily from mid-latitude colonisations rather than equatorial extirpations. Colonisations were particularly frequent in the North Atlantic. Localized extirpations in the western Pacific highlight small-scale patches of vulnerability, that spatially aggregated richness metrics mask. Our results underscore the need to distinguish between trailing and leading processes in climate-induced range shifts, and to consider spatial variability in monitoring and protection of marine biodiversity.

## 1 Introduction

The latitudinal diversity gradient (LDG) of marine species is often attributed to the sea surface temperature (SST) gradient, as higher thermal energy can enhance productivity and thereby support greater diversity (Roy et al., 1998; Tittensor et al., 2010). Yet, despite SST generally peaking at the equator and declining poleward, the LDG of open-ocean taxa is typically bimodal, with a tropical diversity depression flanked by mid-latitude (at ~20°-50°) peaks (Rutherford et al., 1999; Tittensor et al., 2010; Chaudhary et al., 2016; Saeedi et al., 2017; Yasuhara et al., 2020). This mismatch between SST and biodiversity indicates that assuming a simple linear relationship between them is an oversimplification. More importantly, the processes that generated this bimodal pattern, remain unresolved. Past periods of warming, such as the last deglaciation, provide an opportunity to examine whether bimodality rises through increased colonisation of species in the mid-latitudes (i.e. by increasing suitable habitat area), or by species losses in the tropics as thermal limits are exceeded, or a combination of both.

Planktonic foraminifera, a widespread group of calcifying zooplankton, offer an exceptional fossil record for examining how marine biodiversity responded to past warming. Like many other open-ocean taxa, they exhibit a bimodal LDG in the modern ocean (Rutherford et al., 1999; Tittensor et al., 2010; Yasuhara et al., 2020). However, their LDG was unimodal during the Last Glacial Maximum (LGM) (19,000 - 22,000 years ago) and shifted towards bimodality around 15 ka, following rapid deglacial warming (Yasuhara et al., 2020; Strack et al., 2022). A detailed North Atlantic study showed that planktonic foraminiferal species moved polewards during this deglaciation, yet no corresponding decline in equatorial richness was observed (Strack et al., 2022), implying that species’ range expansions may have driven the emergence of bimodality. What remains unclear is whether such range shifts occurred across ocean basins and symmetrically between the hemispheres, how local imbalances between colonisations and extirpations scale up to form regional and global biodiversity patterns, and how these dynamics relate to species’ thermal affinities. Addressing these questions is essential for understanding species biogeographical responses to climate change.

Here, we hypothesize that deglacial poleward range shifts drove the emergence of the modern bimodal LDG in planktonic foraminifera - a mechanism consistent with the poleward movements observed in many marine taxa under current climate warming (Sunday et al., 2012; Poloczanska et al., 2013). Using global, taxonomically standardized datasets of modern and LGM planktonic foraminiferal assemblages (Siccha and Kucera, 2017; Jonkers et al., 2024), we investigate the balance between trailing-edge range contractions (equator-facing) and leading-edge range expansions (pole-facing). We further dissect these dynamics by ocean basins to assess spatial variation in species turnover. Our analyses reveal that species’ responses to deglacial warming differed between the Pacific and Atlantic Oceans, and that despite this regional heterogeneity in species gains and losses, extensive mid-latitude colonisations were pivotal in generating the modern bimodal LDG.

## 2 Methods

### 2.1 Assemblage composition data

Ocean-floor sediment assemblages preserve a record of planktonic foraminifera community composition through time (Kucera, 2007). We used coretop assemblage data of planktonic foraminifera from the ForCenS database (Siccha and Kucera, 2017), which represents the pre-industrial baseline (Jonkers et al., 2019) - hereafter referred to as the “modern” dataset. The sampling methodology for all entries in the database is standardized and consists of counting around 300 specimens larger than 150 μm from core-top samples, with nearly all individuals identified to species-level based on shell morphology (Siccha and Kucera, 2017). For this present work, we used data that was previously cleaned and published (Rillo et al., 2022). The cleaned data consists of relative abundances from 3802 unique samples distributed across the global ocean.

The ForCenS-LGM database (Jonkers et al., 2024) is the Last Glacial Maximum counterpart to the ForCenS database and follows the same criteria for inclusion. Most cores from the LGM were sampled at several core depths (i.e., at different ages). Abundance data in the ForCenS-LGM is presented as they were originally reported in the primary source, i.e., whenever possible, count data is archived, while the remainder are recorded as relative abundances (Jonkers et al., 2024). We standardized taxonomy to be directly comparable to the modern dataset: we removed all sites that did not distinguish between *Globorotalia menardii* and *Globorotalia tumida*, *Globigerinoides ruber* pink and white, and *Turborotalita humilis* and *Berggrenia pumilio* (as in Rillo et al., (2022) thereby removing 19 sites from the Pacific, one from the South Atlantic, and 10 from the Mediterranean. The final dataset consists of relative abundances from 2100 samples, from 630 unique sites. We used the 35 species (Supplementary Table 1) that were present in both the LGM and in the modern datasets for our study, from a total of 50 known living species (Brummer and Kučera, 2022).

### 2.2 Latitudinal diversity gradient of species richness

Species-specific sensitivity (or resistance) to dissolution may affect the species composition recorded in sediments deposited below the calcite lysocline (Berger, 1970). To minimise the effect of dissolution on the biodiversity patterns, we conduct all of our analyses using only samples that were recovered from water depths above 3000 m as these are generally above the calcite lysocline (Yasuhara et al., 2020; Rillo et al., 2022)). Following Yasuhara et al., (2020), we included the fifty-three samples from the ForCenS-LGM that did not have any water depth information. This dissolution filter resulted in 1570 samples in the LGM dataset from 369 unique sites (Fig. 1a), and 2086 unique sites in the modern dataset (Fig. 1b).

**Figure 1:**
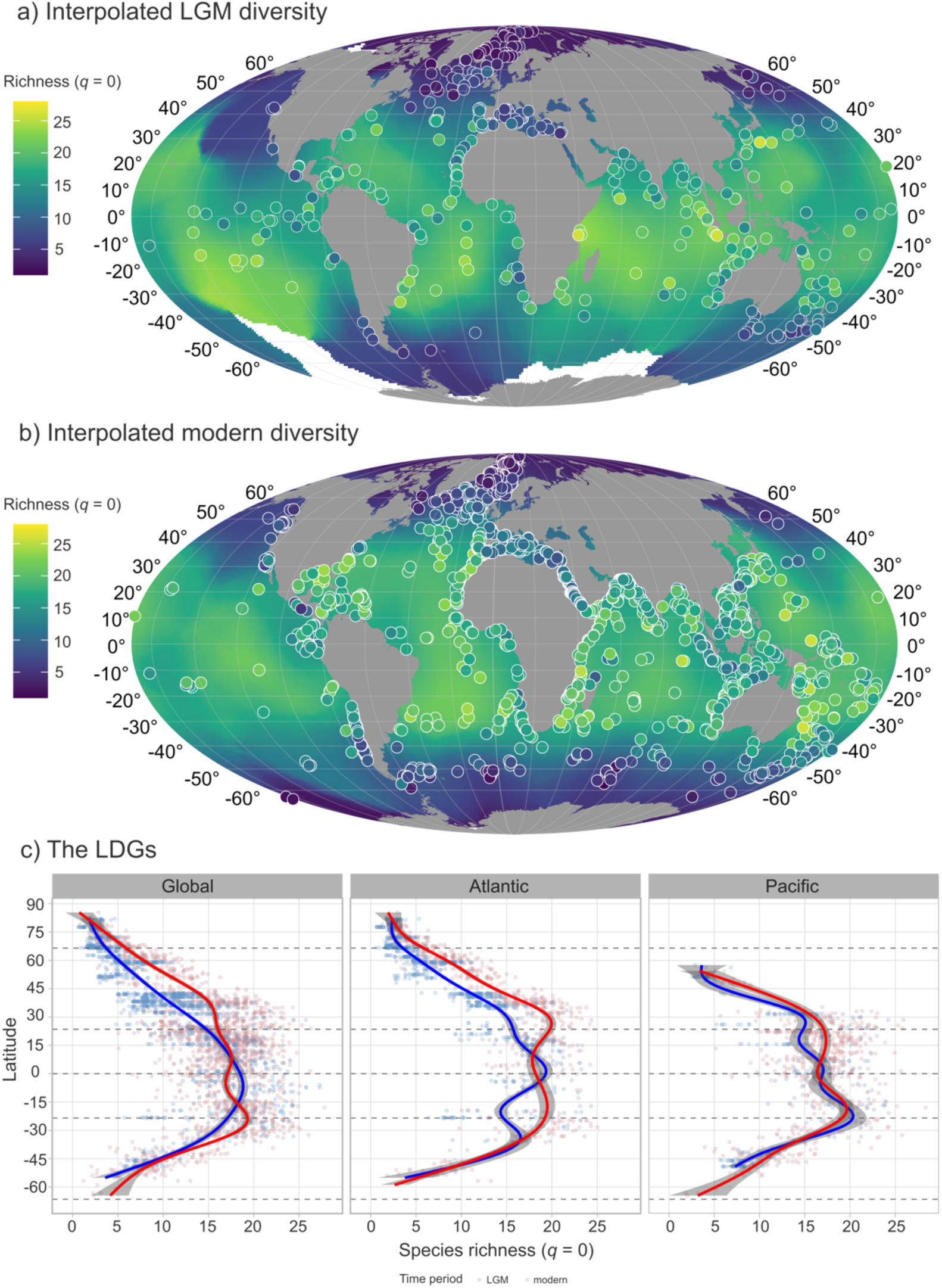
Global patterns of species richness (*q* = 0). Species richness in the **(a)** LGM samples (dots) and **(b)** modern samples (dots), with global richness interpolated over a 12-degree grid based on richness in these sites. White areas indicate the lack of samples in proximity. **(c)** The latitudinal diversity gradient (LDG) of species richness in the Global Ocean, the Atlantic Ocean, and the Pacific Ocean. Dots refer to species richness in the samples, while the solid lines show the GAM-smoothed trends in LGM (blue) and modern (red) samples. Maps are on the Mollweide projection.

We performed all statistical analyses with R 4.5.1 (R Core Team, 2022) (see deposited code). We used the “*specnumber”* and *“diversity”* functions in the *vegan* package (Oksanen et al., 2022) to calculate Hill numbers: the effective number of species at specific orders of biodiversity (Hill, 1973; Chao et al., 2014). At Hill order *q* = 0, diversity corresponds to species richness, while at *q* = 1, diversity considers equally richness and evenness of the assemblages (Chao et al., 2019). Higher orders of biodiversity give increasing weight to common species. We visualised diversity patterns in the two time bins globally by interpolating from the site-scale diversity data with iterated rotations of a penta-hexagonal grid using the *“grapply”* function from the *icosa* package (Kocsis, 2024). The interpolated rasters show expectations for core-scale diversity in a randomly-oriented, 12 degree edge-length grid (Fig. 1a-b).

To quantify the LDGs, we used a GAM-smoothing function on the data (Fig. 1c). Fig. 1 shows results at Hill number *q* = 0; see Fig. S1 for *q* = 1. We further investigated the LDG patterns in the two largest ocean basins, i.e., the Atlantic and the Pacific oceans (Fig. 1c). For the Pacific, we used the cores that were labeled as “Pacific” in the LGM compilation and “North Pacific” and “South Pacific” in the modern compilation. For the Atlantic, we merged “North Atlantic”, “South Atlantic”, “Arctic”, and some “Southern” ocean records (as in (Rillo et al., 2022)). In order to assess longitudinal variation, we investigated the LDG in the eastern and western subsets (Fig. 2, see Fig. S2 for *q* = 1). We split our samples in the Atlantic Ocean along the Mid-Atlantic Ridge (Fig. 2) diverging plate boundaries (Bird, 2003) accessing the boundaries via the *tectonicr* package (Stephan et al., 2023) and using the *spatstat* package (Baddeley and Turner, 2005) to create the subset masks. We split the Pacific split at 150°W (Fig. 2) by convention of the Oceanic Fisheries Programme (Harley et al., 2015).

**Figure 2:**
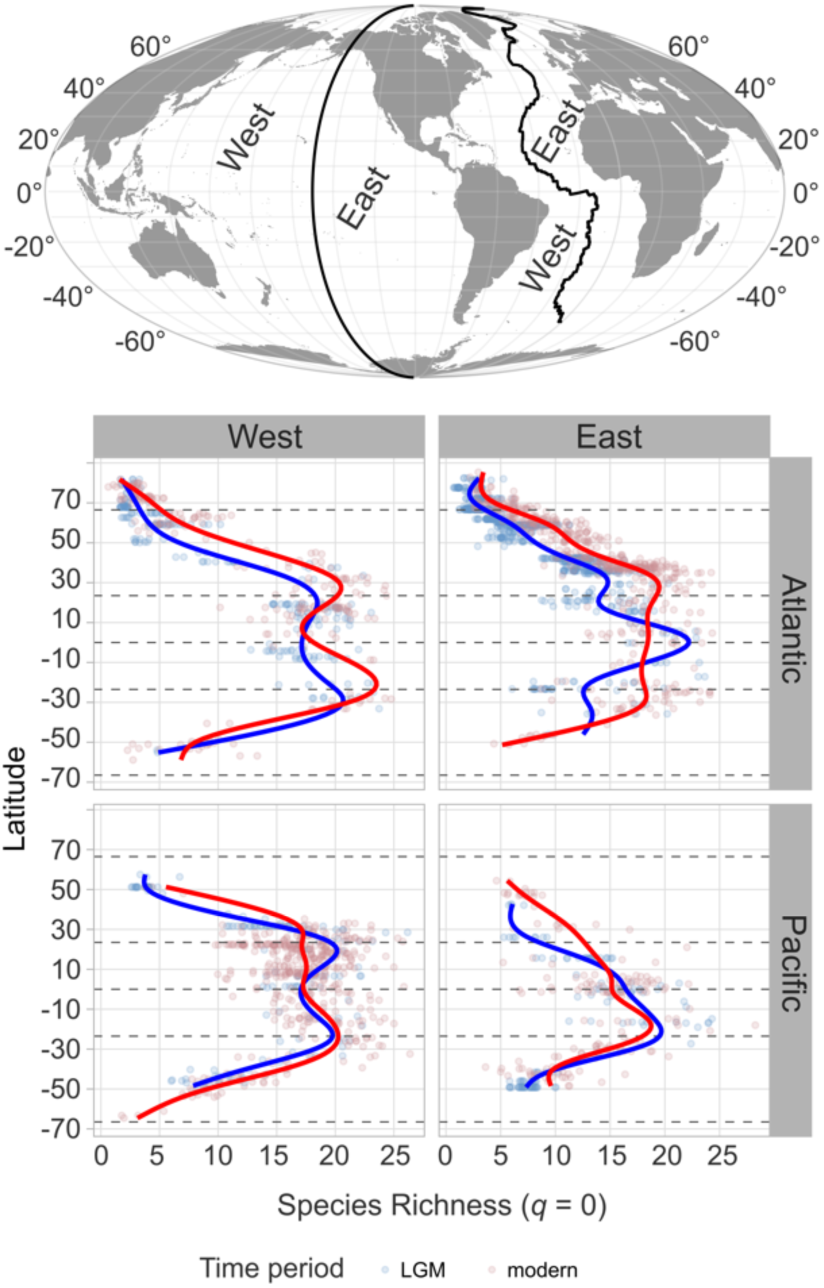
The LDG of species richness (*q* = 0) in the eastern and western Atlantic and Pacific Oceans. The Mid-Atlantic Ridge is used to demarcate between the western and eastern Atlantic, while the 150°W longitude line is used to separate the Pacific Ocean. Map is on the Mollweide projection.

### 2.3 Species latitudinal ranges

We calculated the latitudinal range of each species “globally”, and in the Pacific and Atlantic (Fig. 3) separately. Here, “globally” refers to the combined dataset from the Pacific and Atlantic oceans, rather than all oceans, as only these two basins provide continuous habitat allowing shifts towards both poles. We applied by-occurrence subsampling (rarefaction) in our range size calculations to account for the large differences between the number of sampled cores in the modern and during the LGM (see Fig. 1). Following the approach of Kiessling et al., (2012) we performed subsampling by drawing the same number of sample cores from each 5° latitudinal band and time interval, and allowed the quota to vary among bins. The quota was determined by the time period which had the lower number of samples in a particular 5° latitudinal bin (usually the LGM). To allow for variability of selected cores among the subsampling trials, the quota was chosen to be 10% lower than the minimum sample size.

**Figure 3:**
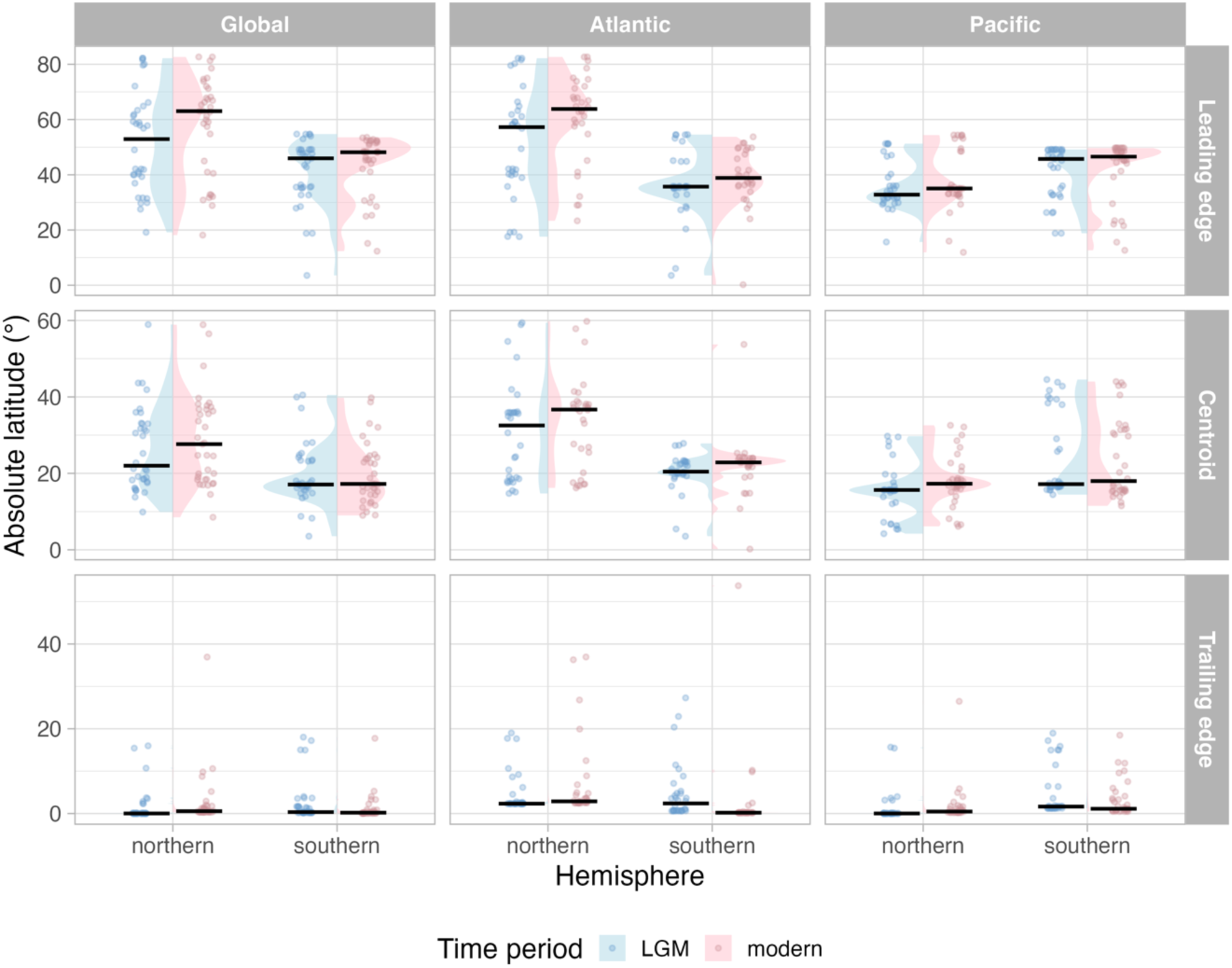
The changing latitudinal distribution of species ranges between the LGM and modern. Violin plots show the density of the edges at a particular latitude. Blue violins show the densities in the LGM; pink violins show the densities in the modern. Jittered points show the underlying data (one dot per species), with triangles referring to LGM samples and dots referring to modern samples. The black horizontal lines highlight the median latitude for each category. Note that because the trailing-edge points cluster at the equator, the trailing-edge violin plots are barely visible or not visible at all.

We report all subsampling results as the mean over 1000 iterations. For each species, the “leading edge” is taken to be the latitude of its most poleward occurrence; the “centroid” is the median latitude of its occupancy per hemisphere, and the “trailing edge” is taken to be the latitude of its most equatorward occurrence. Effect sizes for each metric (Supporting Information S2) were calculated as the difference between modern and LGM values (modern − LGM), and 95% confidence intervals for the effect sizes were estimated using bootstrap resampling with 20,000 iterations (Fig. S3). Statistical significance is determined by whether the CI overlaps with zero (Fig. S3). One degree of latitude corresponds to approximately 111 km.

### 2.4 Assemblage compositional change

In order to identify the temporal changes in diversity at the assemblage scale, we assessed differences in LGM-modern pairs of sites in the Pacific and the Atlantic oceans. We calculated the great circle distance between every LGM site and every modern site and matched the sites with the smallest distance within a radius of 120 km (as in Jonkers et al., (2023)). This radius is reasonable for capturing local temporal changes given the known spatial integration of planktonic forams in sedimentation (v. Gyldenfeldt et al., 2000). If a LGM site had more than one sample (i.e., when the same core was sampled at multiple ages), we randomly sampled one.

We identified 104 pairs of sites in the Pacific and 372 pairs of sites in the Atlantic (Supporting Information S2). For each pair, we calculated the percentage change in the species richness (*q =* 0) through time, divided by the number of species in the LGM (Fig. 4a). We assessed temporal turnover by calculating the Morisita-Horn dissimilarity (Fig. 4b), a measure of compositional turnover that takes species abundance into account (Morisita, 1959). The Morisita-Horn dissimilarity, ranging from 0 to 1, is the probability that two randomly sampled individuals from two assemblages do not belong to the same species, with 0 indicating identical communities, and 1 indicating completely different communities.

**Figure 4:**
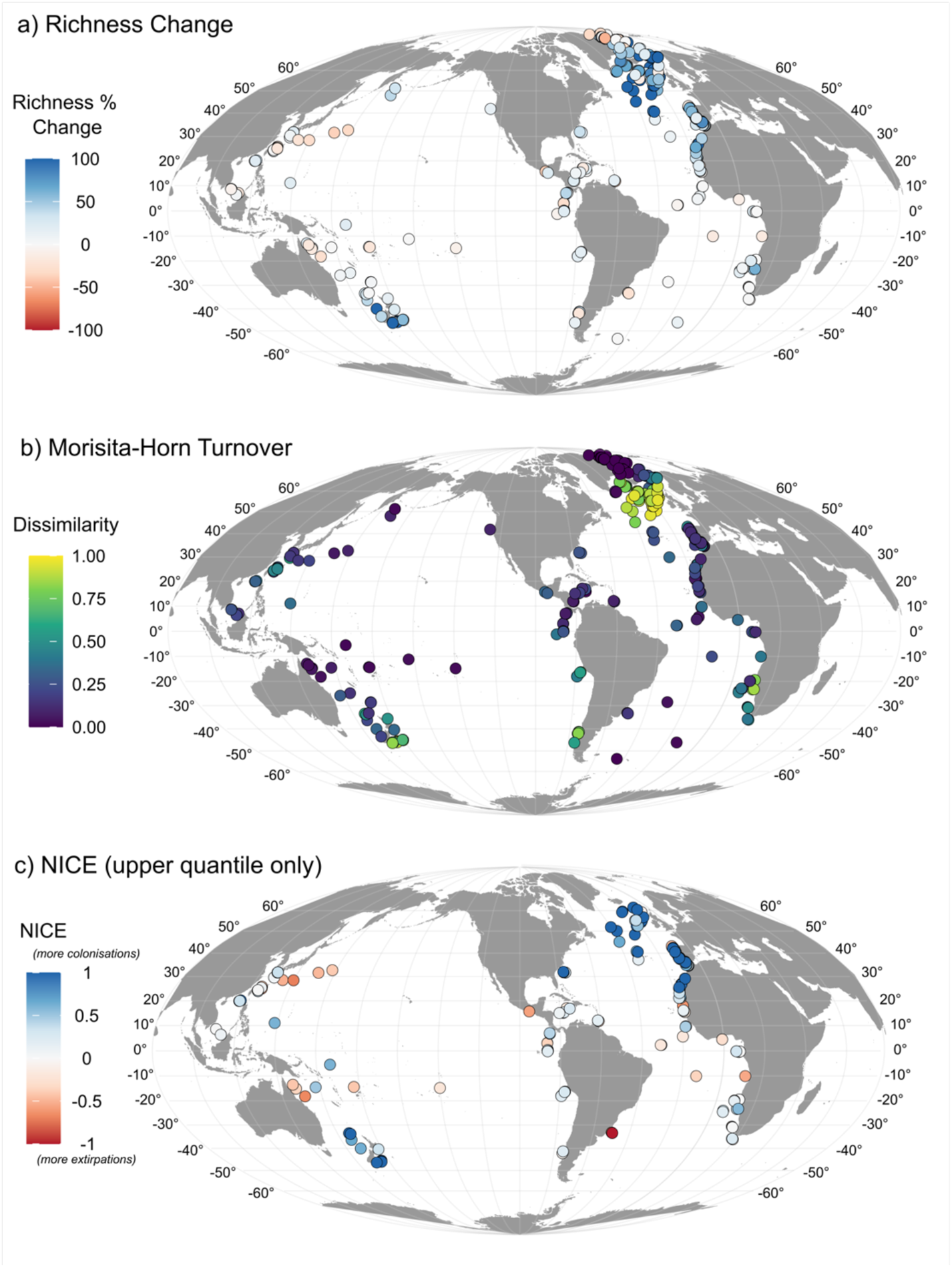
Temporal dynamics of planktonic foraminifera diversity between LGM-modern pairs of sites in the Pacific and Atlantic basins. **(a)** Percentage change in species richness from the LGM to the modern, with reds indicating losses since the LGM, and blue indicating gains since the LGM. **(b)** Temporal turnover between the LGM and the modern, with blues indicating identical communities and yellows indicating completely different communities. **(c)** The spatial distribution of species extirpations (red) and local colonisations (blue), plotted as the Net Imbalance between Colonisations and Extirpations (NICE). Only LGM-modern site pairs with gains or losses above the 75^th^ percentile are shown in (c), representing regions of exceptionally high species turnover. Maps are on the Mollweide projection. Coordinates represent the mean latitude and longitude between each LGM site and its nearest modern site within a 120km radius.

### 2.5 Species extirpations and colonisations

Using the LGM-modern sample pairs, we quantified, per site, how many and which species went extinct between the LGM and the modern - and how many and which species colonised new sites. We calculated the effect size of these changes in each pair of sites as such: for the percentage of species extirpating, we divided the number of lost species by the number of species present in the LGM sample, and for the percentage of species colonising, we divided the number of new species by the total number of species in the modern sample.

To quantify spatial patterns of species gains and losses, we calculated, for each pair of samples, the Net Imbalance between Colonisations and Extirpations (NICE; (Kuczynski et al., 2023)).

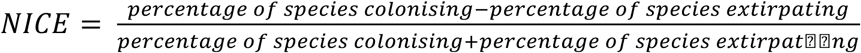

Negative NICE values indicate more extirpations than colonisations from the LGM to the modern, and positive values indicate more colonisations than extirpations. For ease of visualization, we only present the pairs of LGM-modern samples which experienced more than the 75^th^ percentile (upper quantile) of either losses or gains, as this helps identify regions with exceptionally high species gains (i.e., species colonising zones) or losses (i.e., species extirpation zones). We performed sensitivity tests on these results to assess the impact of rare species on our extirpation and colonisation zones by excluding species that had a relative abundance of <1%, and <5% within a given sample (Fig. S4).

To quantify species-specific patterns of range shifts, we calculated the number of extirpation and immigration events each species experienced between the LGM and the modern pairs, expressed as proportional changes (Fig. S5). We then assessed whether species’ modern thermal distributions explained their propensity for extirpation or colonisation. For each species, we calculated the Species Thermal Index (STI, the abundance-weighted median of the thermal distribution) and the Species Thermal Range (STR, abundance-weighted range, defined as the 90^th^ minus the 10^th^ percentile of the thermal distribution). Thermal distributions per species were calculated using the modern assemblage data (Siccha and Kucera, 2017) and SST data from the Extended Reconstructed Sea Surface Temperature from 1854 to 1899 (ERSST ver. 5; (Huang et al., 2017)), following the data retrieval and methodology described in Rillo et al., (2022) and Burrows et al., (2019). Finally, we plotted the proportion of colonisation and extirpation events against STI and STR to explore relationships between species’ thermal distribution and range dynamics; we explored these relationships using generalized linear models and a quasibinomial error structure to account for proportional response variables (Fig. 5, Fig. S6).

**Figure 5:**
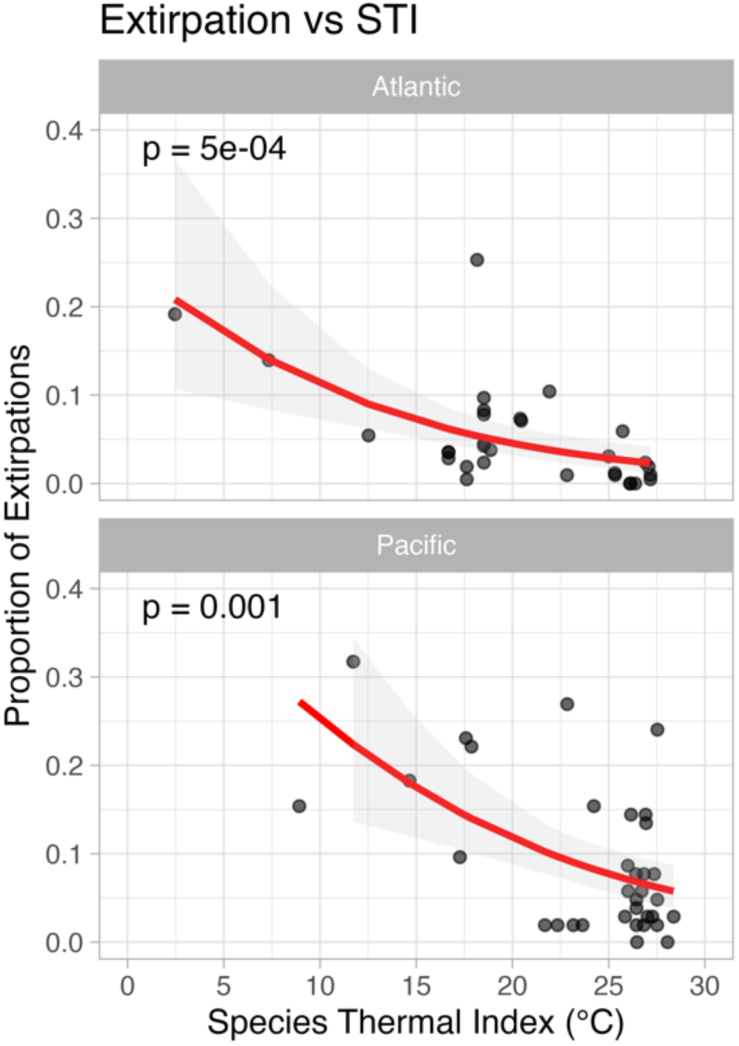
The relationship between species’ thermal indices (STI), and the proportion of extirpations each species faced in the LGM-modern pairwise assessments in a) the Pacific Ocean, b) the Atlantic Ocean.

To catch more nuanced changes, in addition to colonisation and extirpation, we also computed for each pair of sites whether a given species experienced a decline in relative abundance, experienced no change, or experienced an increase in relative abundance (Supporting Information S3). We also spatially interpolated each species’ relative abundance globally during the LGM and the modern, the same way as we interpolated diversities.

## 3 Results

### 3.1 Reassessing the LDG

The global assessment of the latitudinal diversity gradient for species richness (*q* = 0) of planktonic foraminifers confirms that the LDG in the Last Glacial Maximum (LGM) was unimodal, reaching highest richness (~19 species) just south of the equator (Fig. 1a, 1c). Modern richness is highest at approximately 23°S (~20 species), with local peaks at 10°N (~18 species) and 35°N (~15 species), and a notable depression at the equator (~16 species) (Fig. 1b, 1c).

The LDG patterns in the Atlantic and Pacific oceans show differing patterns of species richness (Fig. 1c). In the glacial Atlantic, richness was highest at approximately 5°N (~19 species), with a local maximum at around 40°S (~17 species). In the modern Atlantic, the LDG is more symmetric, with two richness peaks at the tropics (~20 species), and a depression at the equator (~19 species) (Fig. 1c). The modern LDG is highly bimodal in the western Atlantic, and flatter in the eastern Atlantic (Fig. 2). Species richness increased through time on both sides in the mid-latitudes, and declined near the equator between 10°S to 10°N; both the increase and decrease are more prominent in the eastern Atlantic (Fig. 2).

In the glacial Pacific (Fig. 1c), richness was greatest at approximately 23°S (~21 species). Local richness peaks were present in the northern hemisphere at approximately 30°N (~16 species) and 8°N (~17 species), and equatorial richness was approximately 17 species. In the modern Pacific, richness is distinctly bimodal and nearly symmetrical, with peaks at 23°N (~17 species) and at 20°S (~20 species), and a distinct depression at the equator (~16 species). Tropics in the northern hemisphere had an increase in species richness (Fig. 1c). However, there is significant longitudinal heterogeneity (Fig. 2): while richness through time increased between 10°N – 50°N in the eastern Pacific, at the western end, the number of species decreased between the equator and 40°N.

The major distribution patterns for the abundance-weighted effective number of species (diversity metric *q* = 1; Fig. S1) are consistent with richness (*q* = 0). The LDG shifted from a glacial unimodal distribution towards a bimodal pattern in the modern oceans. Equatorial diversity declined from the LGM to the modern, while richness in the subtropics and mid-latitudes increased. These trends occurred globally and within both oceans, but like species richness (Fig. 2), the changes were not uniform along longitude, showing clear east-west differences (Fig. S2).

### 3.2 Geographic ranges of species in the LGM and in the recent

Key metrics of latitudinal species distributions - leading edges, centroids, and trailing edges - are shown in Fig. 3 and Table 1. Globally, species’ leading edges shifted poleward from the LGM to the modern (Fig. 3). The greatest changes occurred in the northern hemisphere (Fig. 3), where the global median leading edge shifted significantly (Fig. S3) poleward by 10° of latitude (1124 km). In contrast, the shift in the southern hemisphere was much smaller, with the median moving from −45.6° to −48°, a distance of only 245 km. Poleward movements were generally greater in the Atlantic than in the Pacific in both hemispheres (Fig. 3 and Table 1).

**Table 1:** The median range shifts between the LGM and the modern for each edge type (leading and trailing), plus the centroid of the species geographical distribution. Effect size is calculated by modern value minus LGM value. Significance is based on whether effect size differs significantly from 0, i.e., whether the 95% CI overlap with zero.

|  | Northern hemisphere |  |  | Southern hemisphere |  |  |
| --- | --- | --- | --- | --- | --- | --- |
|  | Lead edge<br>effect size | Centroid<br>effect size | Trail edge<br>effect size | Lead edge<br>effect size | Centroid<br>effect size | Trail edge<br>effect size |
| Global | 10.1°<br>1124 km<br>polewards<br>(significant) | 5.6°<br>626 km<br>polewards<br>(ns) | 0.5°<br>59 km<br>polewards<br>(significant) | 2.2°<br>245 km<br>polewards<br>(ns) | 0.14°<br>16 km<br>polewards<br>(ns) | -0.15°<br>16 km<br>equatorwards<br>(ns) |
| 95% CI of<br>effect size | [2.8°; 23.0°] | [-1.2°; 9.7°] | [0.3°; 1.1°] | [-10.7°; 1.5°] | [-5.3°; 1.7°] | [-0.03°; 1.3°] |
| Atlantic<br>Ocean | 6.6°<br>729 km<br>polewards<br>(significant) | 4.2°<br>462 km<br>polewards<br>(significant) | 0.54°<br>59 km<br>polewards<br>(significant) | 3.1°<br>349 km<br>polewards<br>(significant) | 2.4°<br>267 km<br>polewards<br>(significant) | -2.2°<br>243 km<br>equatorwards<br>(significant) |
| 95% CI of<br>effect size | [2.9°; 22.8°] | [1.1°; 12.6°] | [0.2°; 1.3°] | [-8.1°; -0.8°] | [-2.9°; -0.9°] | [0.4°; 3.5°] |
| Pacific<br>Ocean | 2.2°<br>248 km<br>polewards<br>(ns) | 1.6°<br>181 km<br>polewards<br>(significant) | 0.45°<br>50 km<br>polewards<br>(significant) | 0.8°<br>89 km<br>polewards<br>(ns) | 0.8°<br>89 km<br>polewards<br>(ns) | -0.5°<br>57 km<br>equatorwards<br>(ns) |
| 95% CI of<br>effect size | [-0.9°; 3.4°] | [0.8°; 2.9°] | [0.3°; 0.7°] | [-10.2°; 1.1°] | [-12.7°; 1.4°] | [-1.2°; 0.8°] |

The poleward shift in the median (centroids) species distribution was greatest in the northern hemisphere, where species’ centroids moved more than 5.6°, equivalent to about 600 km polewards. In contrast, the shift in the southern hemisphere was just 0.14°, or 16 km (Fig. 3, Table 1).

The trailing edges did not notably shift polewards, and most species maintained their equatorial occupancy (Fig. 3, Table 1). In the northern hemisphere, trailing edges moved poleward by only approximately 0.5° (59 km), while in the southern hemisphere they even moved slightly towards the equator by 0.15° (16 km). Though some trailing edge shifts were statistically significant (Fig. S3), this is likely due to tight clustering of points.

The vast majority of species did not show any significant change in their trailing edge occupancy from the LGM to the modern (Supplementary Table 1). Some rare (i.e., low relative abundance) species had an effect size in their trailing edges of more than 5° latitude (555 km): *Candeina nitida*, *Dentigloborotalia anfracta, Globigerinella adamsi, Globigerinita uvula, Globorotalia hirsuta, Sphaeroidinella dehiscens, Globorotalia menardii, Globorotalia theyeri, Globorotalia tumida, Sphaeroidinella dehiscens, Tenuitella iota, Turborotalita humilis,* but the direction of these movements was not uniform.

### 3.3 Site-level changes in species richness and community composition

Localised changes in species richness and community composition were calculated from LGM-modern pairs of sites (Fig. 4, Supporting Information S2). The change in relative abundance of each species through time can be found in Supporting Information S3.

In the Atlantic Ocean, within the Arctic circle, sites both gained and lost species since the LGM (Fig. 4a). Overall, species composition remained relatively constant, with low turnover (Fig. 4b), except in the mid-northern latitudes where many species were gained (Fig. 4a), creating highly dissimilar communities (Fig. 4b). Species richness remained relatively constant in the tropical open ocean between 23.5°N to 23.5°S though there were changes in species composition (Fig. 4a-b). There is little evidence of longitudinal heterogeneity in diversity change within the northern Atlantic Ocean, but in the southern Atlantic, while richness decreased and turnover was low off South America, richness increased, and dissimilarity was higher off the coast of Africa.

In the Pacific Ocean, localised species richness declined most strongly in the tropical northwestern Pacific (median loss of 20%, 13 pairs of sites), while it increased most notably south of 20°S (~33%, 27 pairs of sites; Fig. 4a). Longitudinal heterogeneity is apparent: richness increased in the eastern Pacific, while on the western end, sites either maintained or lost species (Fig. 4a). Temporal compositional change does not always mirror the richness trends in the Pacific (Fig. 4b). In general, dissimilarity is low at sites where species richness declined, suggesting nestedness, and high where species richness increased, suggesting turnover. However, dissimilarity is also high off the coast of South America, where richness decreased or was maintained (Fig. 4b).

### 3.4 Species extirpations, colonisations and thermal preferences

In the Atlantic, the mean number of species extirpating through time (per pair of LGM-modern sites) is 1.5±0.1; in the Pacific it is 3.1±0.2. The mean number of species colonising in the Atlantic is 4.1±0.1; in the Pacific it is 3.9±0.2. The upper quantiles of the net imbalance between colonisations and extirpations (NICE, Fig. 4c) show latitudinal patterns in the Atlantic Ocean, with little longitudinal differences. In the northern Atlantic mid-latitudes, the vast majority of sites gained more species than lost (Fig. 4c, blue dots), showing a large number of colonisations between approximately 30-65°N. The equatorial latitudes have some notable species declines (Fig. 4c, red dots). Gains were more muted in the southern hemisphere, with many sites also experiencing losses (20-40°S). One site off the coast of South America had disproportionately high losses (Fig. 4c, red dot south coast of Brazil).

In contrast, in the Pacific Ocean, there is strong longitudinal heterogeneity of local extirpations and colonisations. There is a clear concentration of species losses in the western Pacific between 20-35°N, while there is no data to support the magnitude of such changes in the eastern Pacific at these same latitudes. A balance of winners and losers is apparent in the equatorial eastern Pacific off the coast of Central America, with some sites gaining more species, and others losing, while sites in the equatorial western Pacific gained more species. Further south, a few sites off the northern coast of Australia (10-20°S) also lost more species than gained (Fig. 4c, red dots). In contrast, at these same latitudes in the eastern Pacific, more species are gained than lost between 10-20°S (Fig. 4c, blue dots). Waters off New Zealand are clear species colonisation zones (Fig. 4c, blue dots).

These spatial patterns are robust to rare species: excluding taxa with <1% or <5% relative abundance per core did not alter the main trends (Fig. S4). The northwestern Pacific remained a hotspot of species losses, expanding to the southwestern Pacific in the more conservative test, while the northern mid-Atlantic consistently gained species; equatorial Atlantic sites experienced notable losses only under the stricter threshold.

The variability by ocean was also species-specific (Fig. S5). In the Atlantic Ocean, only *G. scitula* was lost in 20% of the pairs, with *N. pachyderma* being lost at 17% of the site-pairs. In the Pacific, the species *T. quinqueloba, G. crassaformis, G. rubescens, G. inflata*, and *G. truncatulinoides* were extirpated in more than 20% of the LGM-modern site pairs assessed. Similarly, species that colonised most sites from the LGM to the modern also differed between the two oceans. In the Pacific, *G. scitula, B. digitata, G. tumida, O. universa, G. menardii,* and *N. dutertrei* colonised more than 20% of the pairs. In contrast, in the Atlantic, the species *G. hirsuta*, *T. sacculifer*, *G. calida, G. ruber* (pink), *G. rubescens,* and *N. dutertrei* colonised more than 20% of the sites. There is a weak but significant negative relationship between species’ thermal indices, STI (i.e., thermal optima), and the proportion of local extirpation events they faced (*p* = 0.001 Pacific; *p* = 0.0009 Atlantic) (Fig. 5). Only in the Pacific Ocean, there is a significant relationship between species’ STI and colonisations (*p* = 0.024), but there is no relationship between species STR and either colonisations or extirpations in either ocean (Fig S6).

## 4 Discussion

The emergence of a bimodal latitudinal diversity gradient (LDG), with low species richness near the equator and peaks at mid-latitudes, has often been linked to climate warming (Rutherford et al., 1999; Tittensor et al., 2010; Chaudhary et al., 2016; Saeedi et al., 2017; Yasuhara et al., 2020). Previous work suggested that climate warming could drive cause the trailing edges of species distributions to shift polewards, as equatorial sea surface temperatures exceed species’ thermal tolerances, reducing tropical diversity (Sunday et al., 2012; Cheung et al., 2013; Burrows et al., 2014; Antão et al., 2020). We tested the hypothesis that the underlying mechanism behind the bimodal planktonic foraminifera LDG was trailing edge contractions in the modern ocean, as a result of equatorial extirpations. However, our results indicate that trailing edge contractions were not the dominant mechanism shaping the modern bimodal LDG in planktonic foraminifera since the Last Glacial Maximum (Fig. 3). Instead, species’ poleward range expansions and associated colonisations played a more substantial role, while equatorial occupancy was maintained (Fig. 4).

Planktonic foraminifera species exhibit high dispersal capabilities (Rillo et al., 2022) and have maintained relatively stable thermal niches over the past 800,000 years (Antell et al., 2021). These properties suggest that during rapid deglacial warming (~15,000 years ago), species primarily tracked shifting isotherms rather than adapting *in situ*, creating novel assemblages as migrating species encountered persisting ones (Strack et al., 2022). While a modelling study suggests that some functional ecogroups of planktonic foraminifera may have exhibited limited acclimatization, the majority of the functional species retained their thermal preferences and migrated polewards, indicating that tracking thermal fronts, rather than adaptive shifts, was the main driver of the distributional changes over the LGM-deglacial transition (Ying et al., 2024). These mechanisms are consistent with the global patterns we observe: the median trailing edge positions changed minimally, while the median leading edges shifted polewards (Fig. 3).

Across the two largest oceans - the Atlantic and the Pacific - spatial heterogeneity in species dynamics is apparent. Net changes in ocean-scale richness provide a useful overview of diversity shifts but conceal distinct underlying processes (Hillebrand et al., 2018): richness increase may reflect colonisations and community restructuring, but no net change can still involve complete species turnover, and richness losses can arise from extirpations (nested loss) or full compositional reshuffling. Our pairwise analyses allow us to disentangle these underlying processes (Fig. 4). The Atlantic exhibits relatively high meridional coherence, with consistent mid-latitude richness increase, particularly in the North Atlantic, due to colonisations associated with deglacial warming (Fig. 4), supporting previous work (Strack et al., 2022; Jonkers et al., 2023). By contrast, the Pacific shows strong longitudinal differences, with net colonisations in some eastern mid-latitudes sites, and net extirpations in the western tropical sites (Fig. 4c). Turnover analyses reveal that the zones of concentrated species losses in the western tropical Pacific are consistent with nested species loss (extirpations) rather than total community restructuring, whereas colonisation hotspots in the south Pacific and north Atlantic reflect substantial compositional change as new species became established and dominant. These site-level patterns produce a patchwork that is not evident in aggregated, ocean basin-scale metrics (Fig. 1), emphasizing the importance of spatial grain.

Extirpation events are weakly but significantly associated with species’ thermal optima: cooler-adapted species are more likely to experience extirpations (Fig. 5, Fig S6), presumably because climate warming pushed some species beyond their thermal limits. At the same time, equatorial regions showed relatively few extirpations and minimal trailing edge shifts, suggesting that either tropical temperatures changed little since the LGM, or that equatorial foraminifera species may have broader thermal ranges than realised during the LGM. Recent work has suggested that some foraminifera species are able to acclimatize to changing temperatures (Ying et al., 2024). Unexpectedly, our results suggest that species with broader thermal ranges experienced more extirpations, though these are not significant (Fig S5); this result is more likely a limitation of model fits than an ecological response. Nevertheless, these thermal traits indicate that while thermal optima may influence vulnerability to extirpation, environmental variability and starting conditions may strongly modulate species responses, explaining why colder-adapted species were more affected, and yet equatorial assemblages remained relatively stable. At face value, these results may therefore indicate unexpected resilience of tropical zooplankton assemblages to global warming, but meaningful predictions of the response of tropical biodiversity requires knowledge of the fundamental thermal tolerance of the species.

Taken together, our results highlight that the processes shaping global biodiversity patterns can be masked when only considering aggregated metrics like global or ocean-scale richness change. Local losses in the Pacific demonstrate that significant species extirpations occurred (Fig. 2, Fig. 4c), but these extirpations are not apparent in the global summaries (Fig. 1c). This pattern variability at smaller spatial scales cautions that aggregated metrics can overlook hotspots of vulnerability, emphasizing the need to also integrate spatial scale into monitoring frameworks. Previous modelling using the RCP 8.5 scenario (business-as-usual anthropogenically driven warming scenario) predicted that the broader Indo-Pacific region may face high extirpation rates of marine species by 2100 (García Molinos et al., 2016). Our pairwise analyses provide fossil evidence of extirpation hotspots in the western Pacific (Fig. 4c), demonstrating that species losses are already spatially structured and that smaller-scale areas can experience disproportionate climate-driven biodiversity decline. More generally, our findings align with predictions that future climate change will threaten not only trailing edge populations but also the core habitats of many species, projected to decline by over 50% under high-emission RCP 8.5 scenarios (Hodapp et al., 2023). Present day observations indicate that even over the past century, when planktonic foraminifera tracked polewards, migration alone was insufficient to buffer populations against the very rapid ongoing and predicted future environmental change (Chaabane et al., 2024). Together, our results underscore the importance of considering multiple spatial scales and dimensions of biodiversity change to understand how global patterns such as the LDG were shaped.

## 5 Code and Data Availability

Supporting Information S1 contains Supplementary Figures S1 – S6 and Supplementary Table S1.

Supporting Information S2 contains all of the results in an Excel Spreadsheet.

Supporting Information S3 is a zipped folder containing figures of interpolated and site level abundance change.

All data and code required to replicate all figures in this paper can be found on GitHub: https://github.com/Mingmingkhan/ldg-forams

## 8 Supporting Information S1

**Supplementary Table 1:**
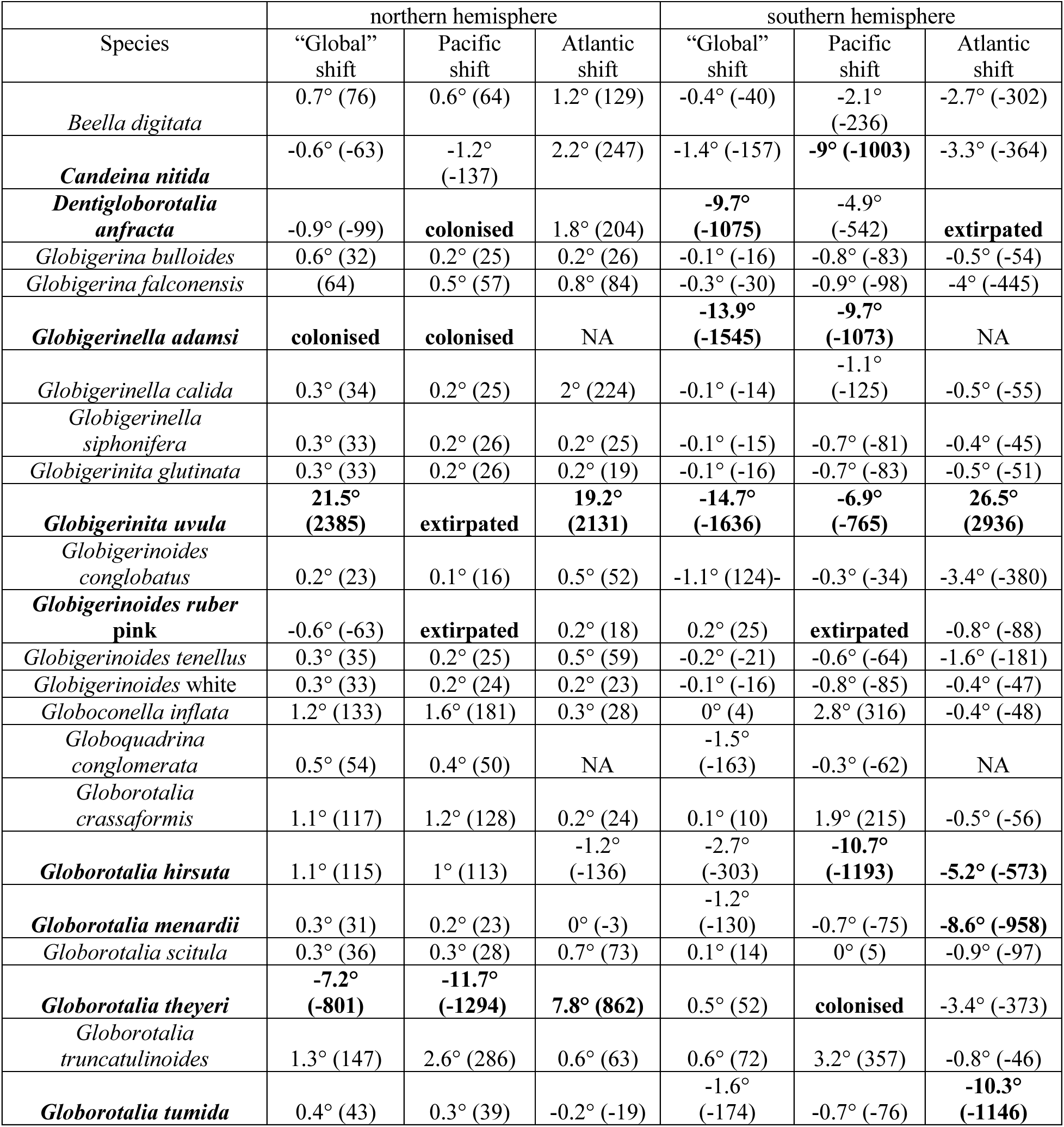
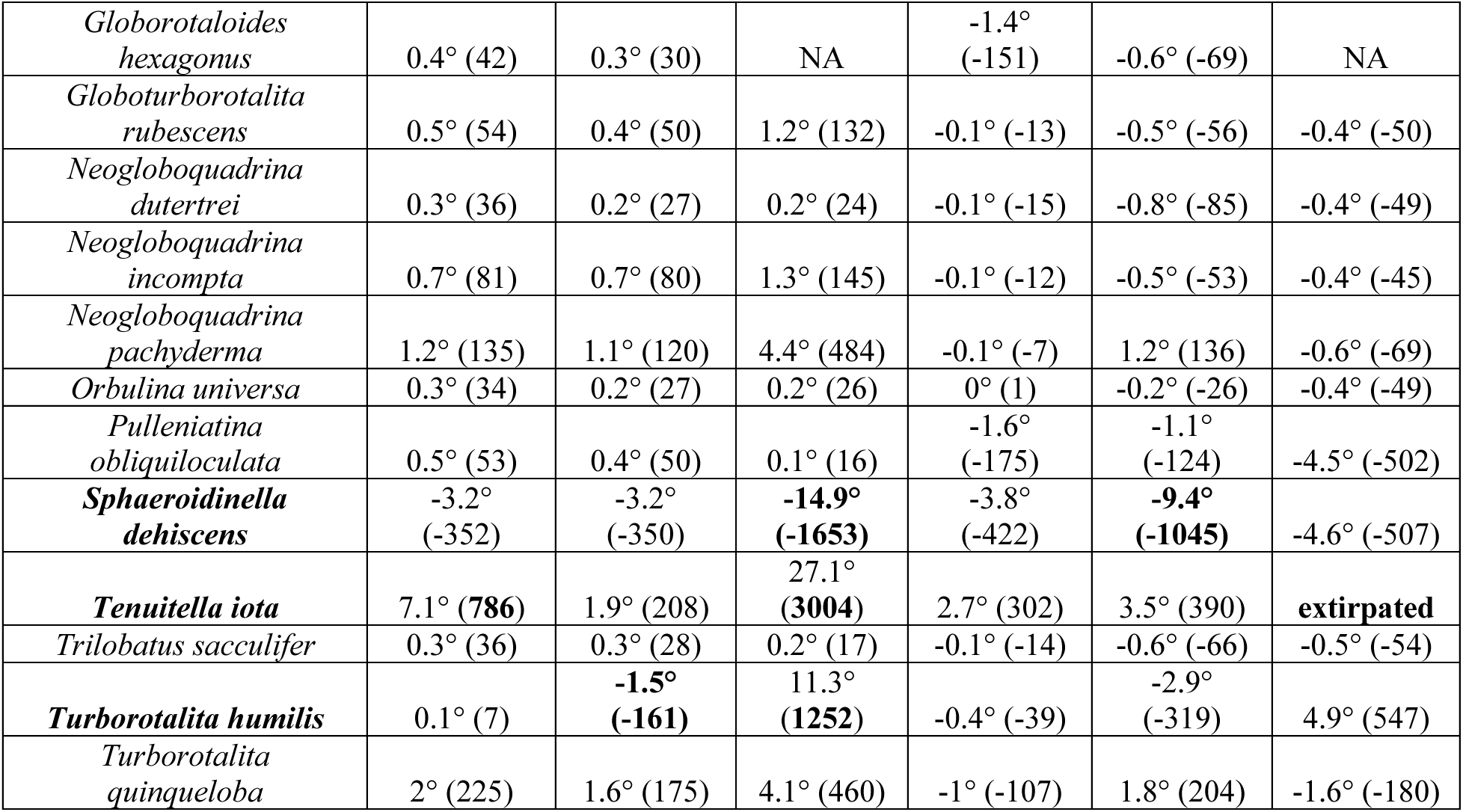
Species’ shifts in the trailing edges, from the LGM to the modern, shown in degrees of latitude (°). The shift in kilometres is in parentheses. Negative values indicate shifts towards the equator, while positive values indicate shifts away from the equator. Values in bold show species that had greater than 5° (555 km) of latitude change in occupancy. Species-specific changes in the leading edges and centroids are available in Supporting Information S2. Note that “global” refers to the combined Pacific + Atlantic dataset.

**Figure S1:**
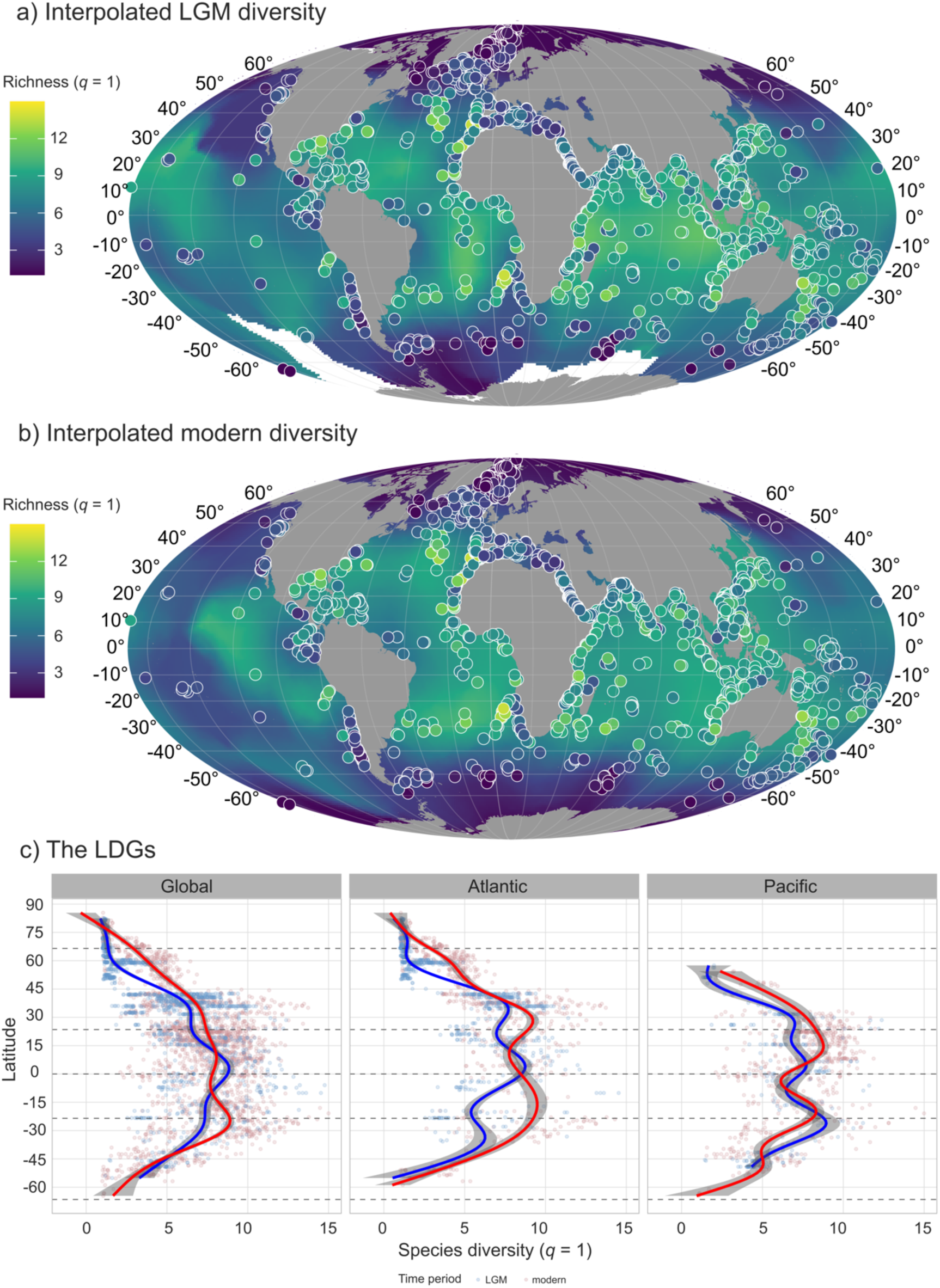
Global patterns of species diversity at *q* = 1 during (a) the LGM, (b) the modern, and (c) the latitudinal diversity gradient at *q =* 1.

**Figure S2:**
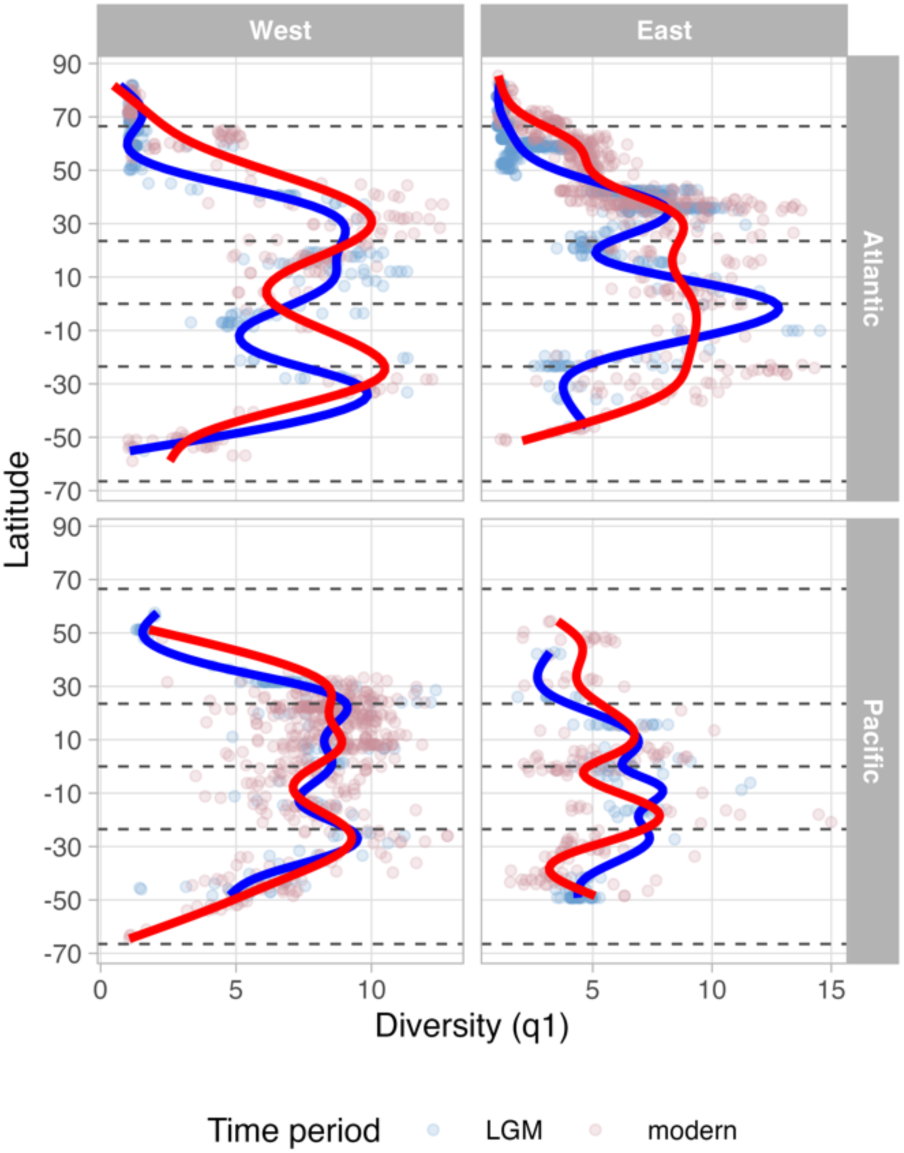
The LDG of species diversity (*q* = 1) in the eastern and western Atlantic and Pacific Oceans. The Mid-Atlantic Ridge is used to demarcate between the western and eastern Atlantic, while the 150°W longitude line is used to separate the Pacific Ocean.

**Figure S3:**
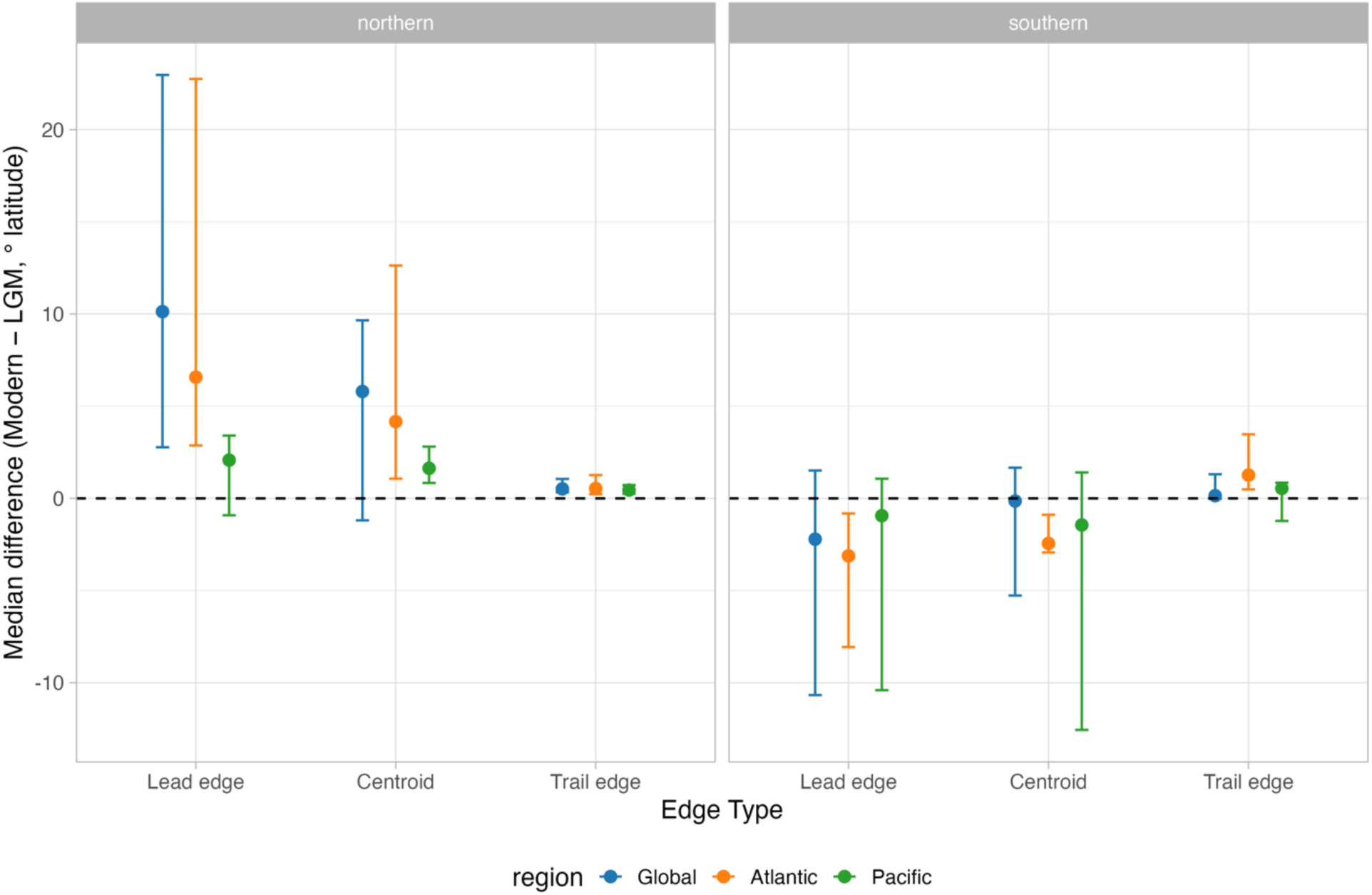
95% confidence intervals of the effect size of range shifts between the LGM and the modern, for each edge type, in the “Global”, Atlantic and Pacific Oceans. The effect size is taken to be significant if the error bars do not overlap with zero.

**Figure S4:**
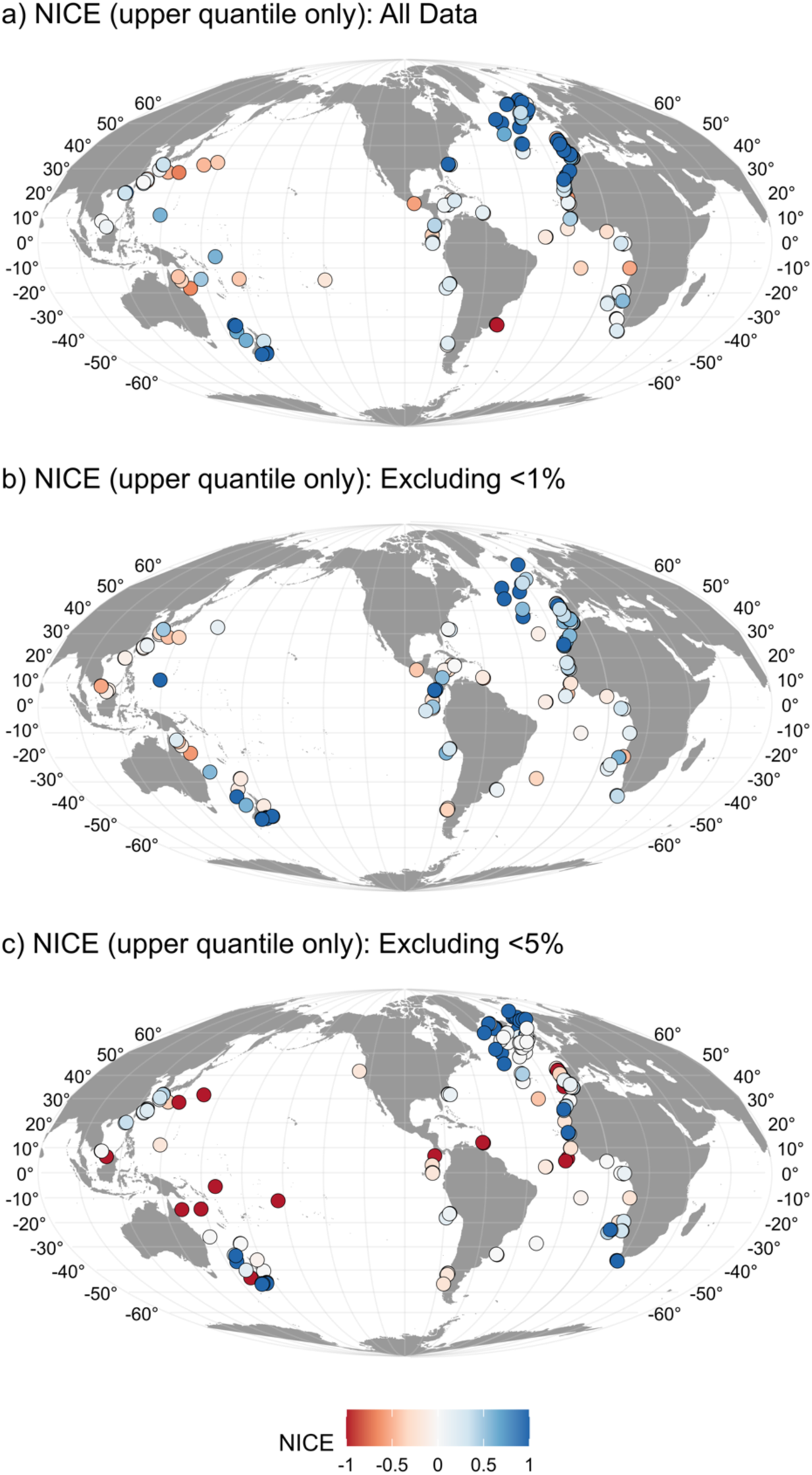
The impact of rare species in determining the spatial distribution of the upper quantile of species extirpations and species hotspots.

**Figure S5:**
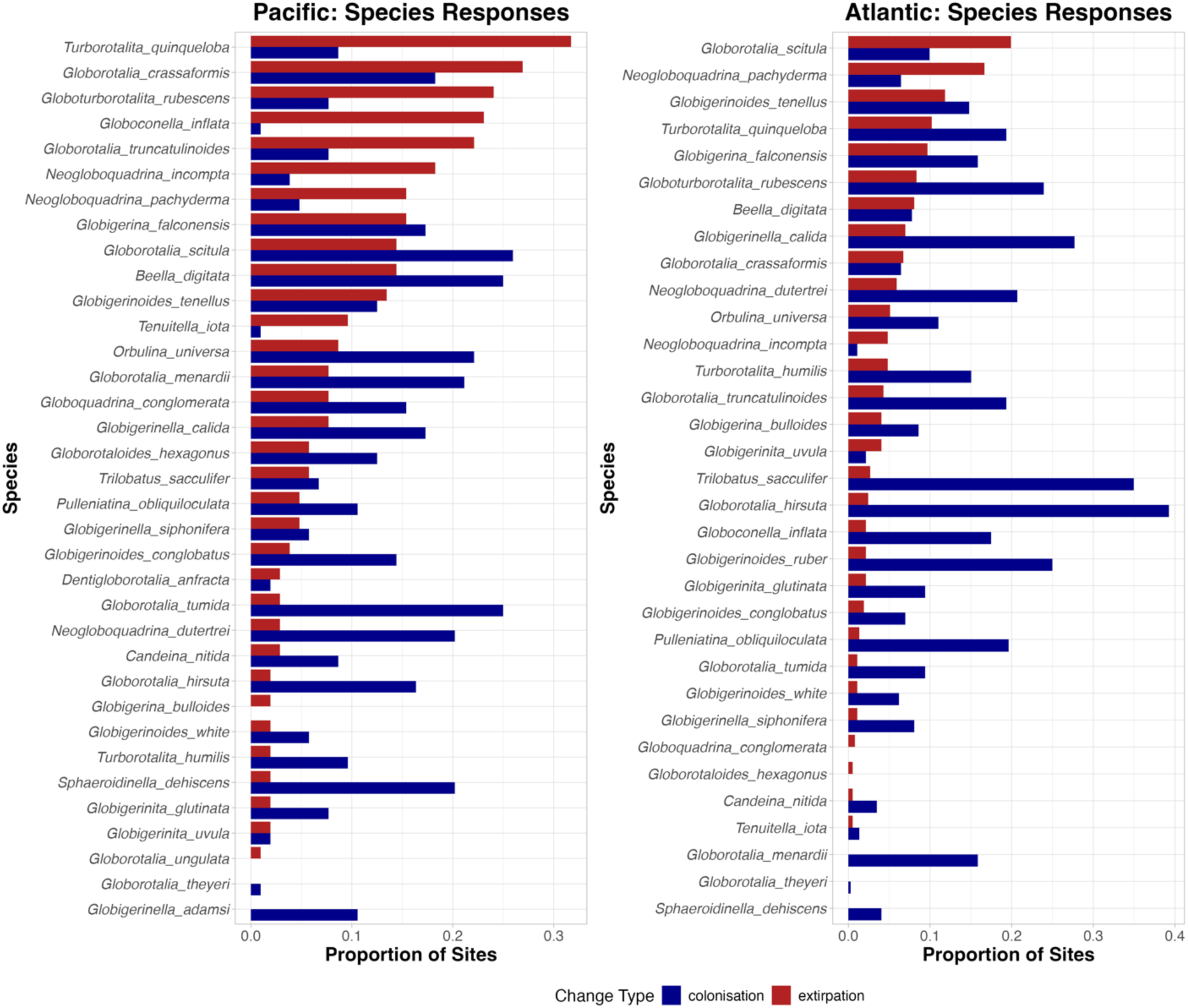
Species-specific responses in the Pacific and Atlantic oceans. Colonisations (blue) and extirpations (red) events within LGM-modern pairs of sites. Species are ordered by the greatest proportion of extirpation events.

**Figure S6:**
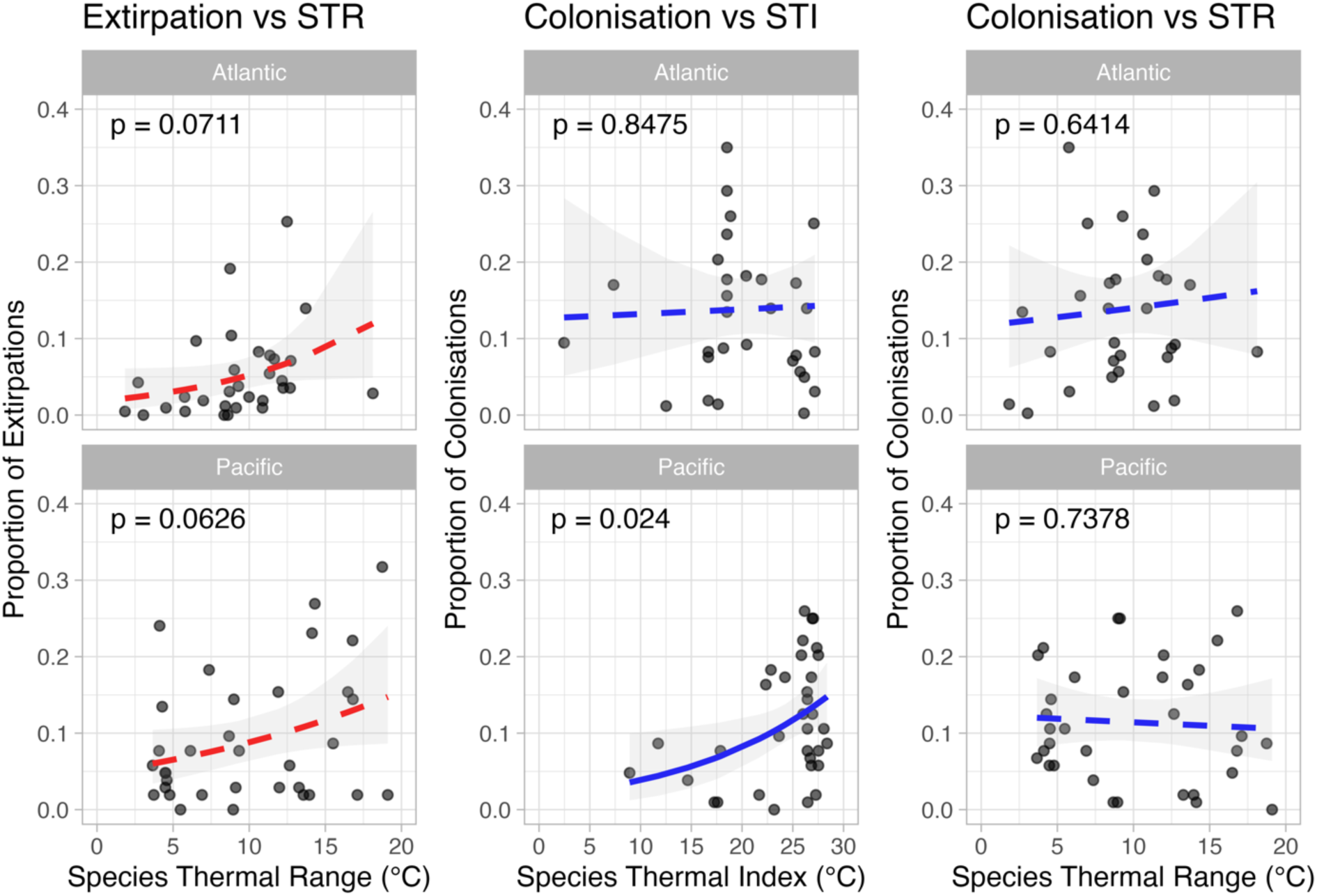
Species’ thermal relationships, and the proportion of colonisation/extirpation events they experienced in the pairwise analyses.

