## Supplementary figures and images for "Asymmetric leading vs. trailing edge shifts since the Last Glacial Maximum underpin the modern bimodal latitudinal diversity gradient in planktonic foraminifera"

### abundance_change_Beella_digitata.png

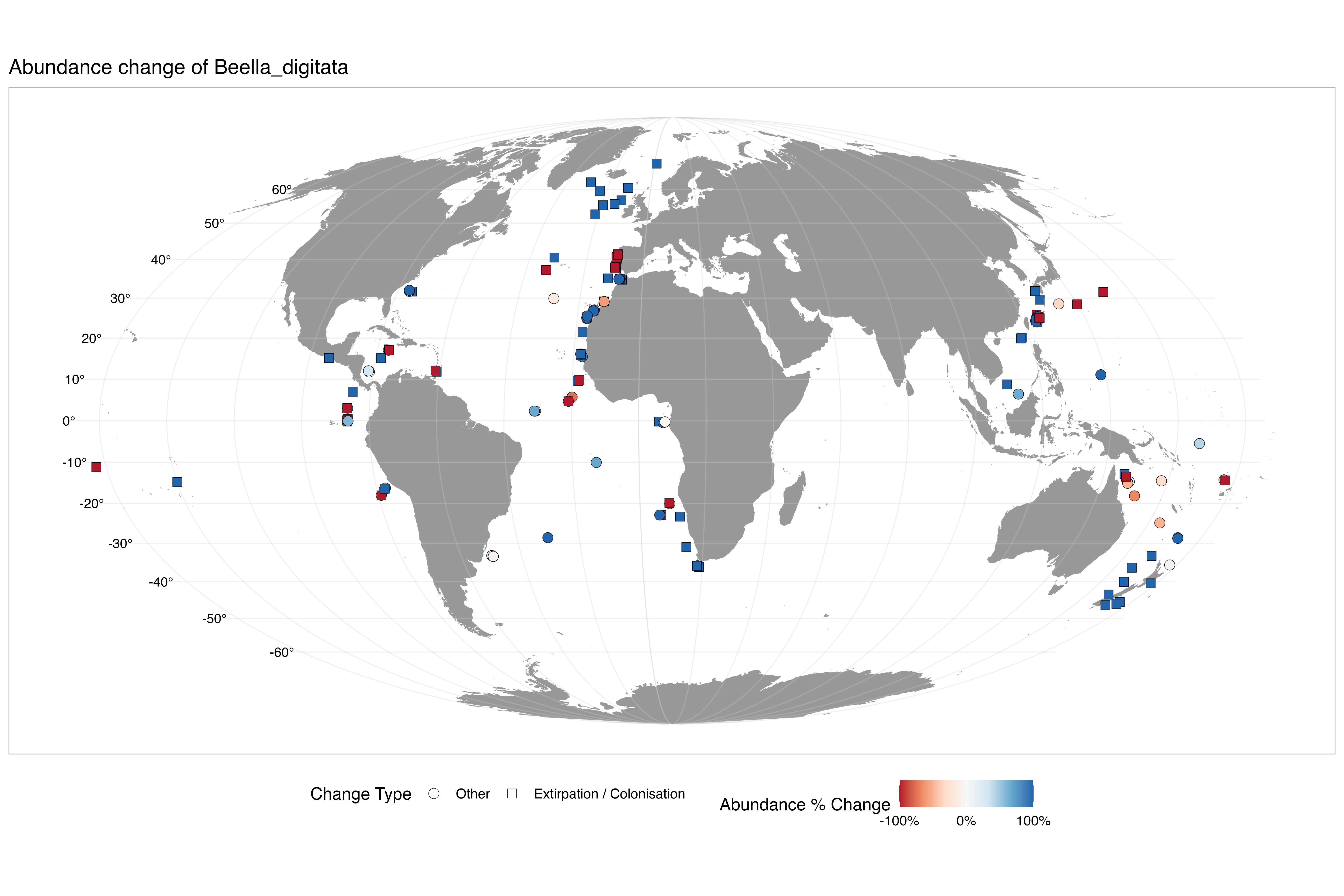

### abundance_change_Candeina_nitida.png

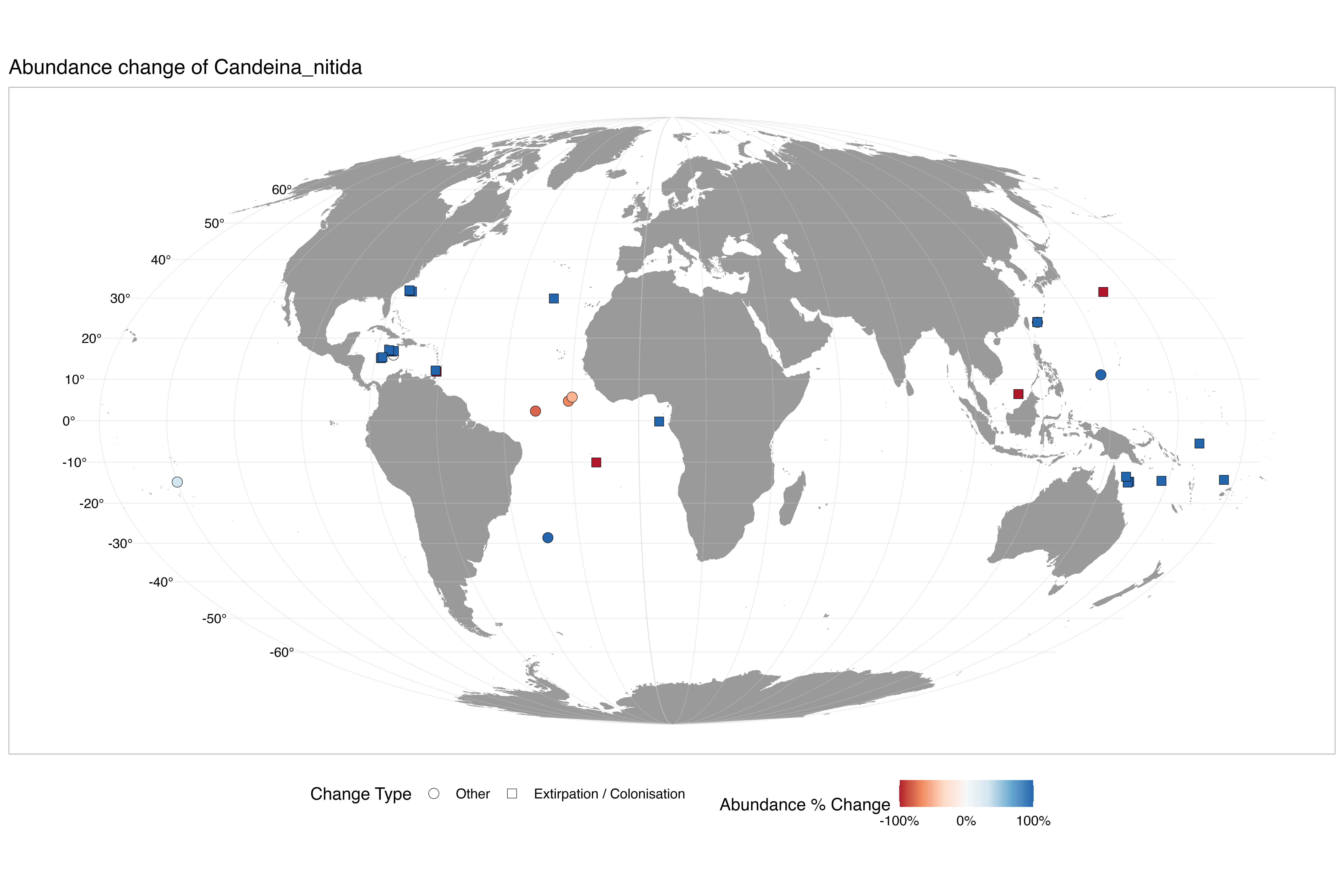

### abundance_change_Dentigloborotalia_anfracta.png

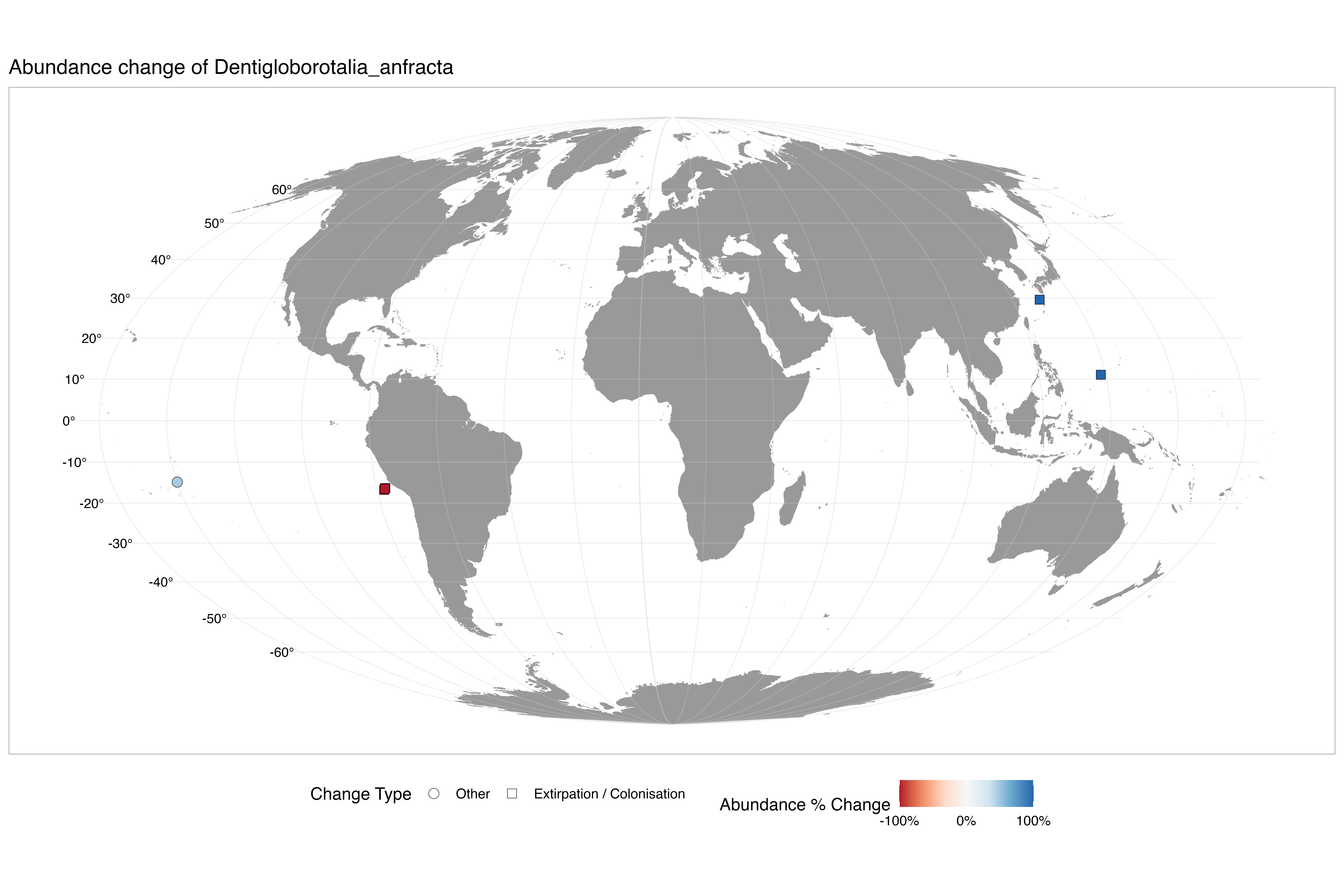

### abundance_change_Globigerina_bulloides.png

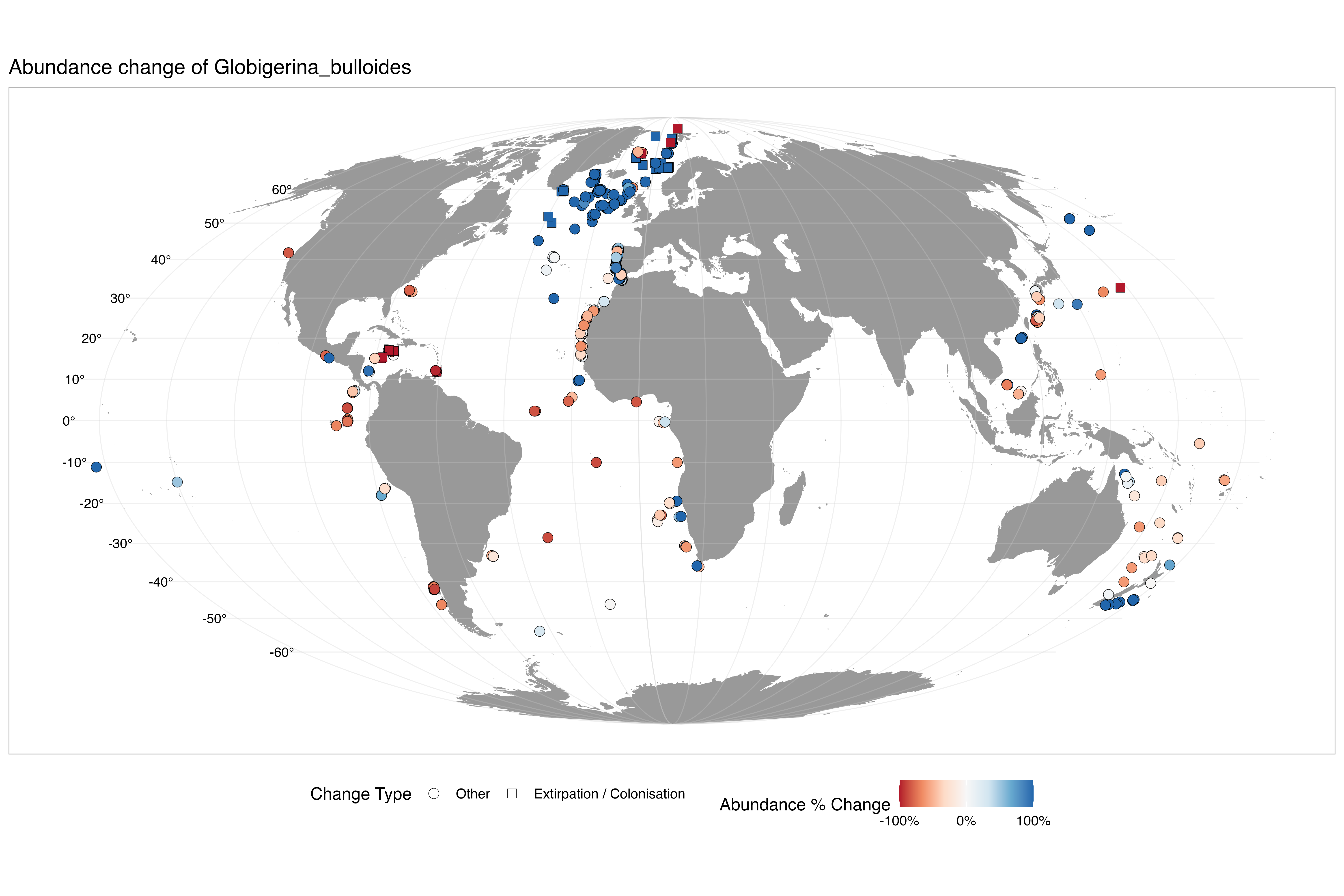

### abundance_change_Globigerina_falconensis.png

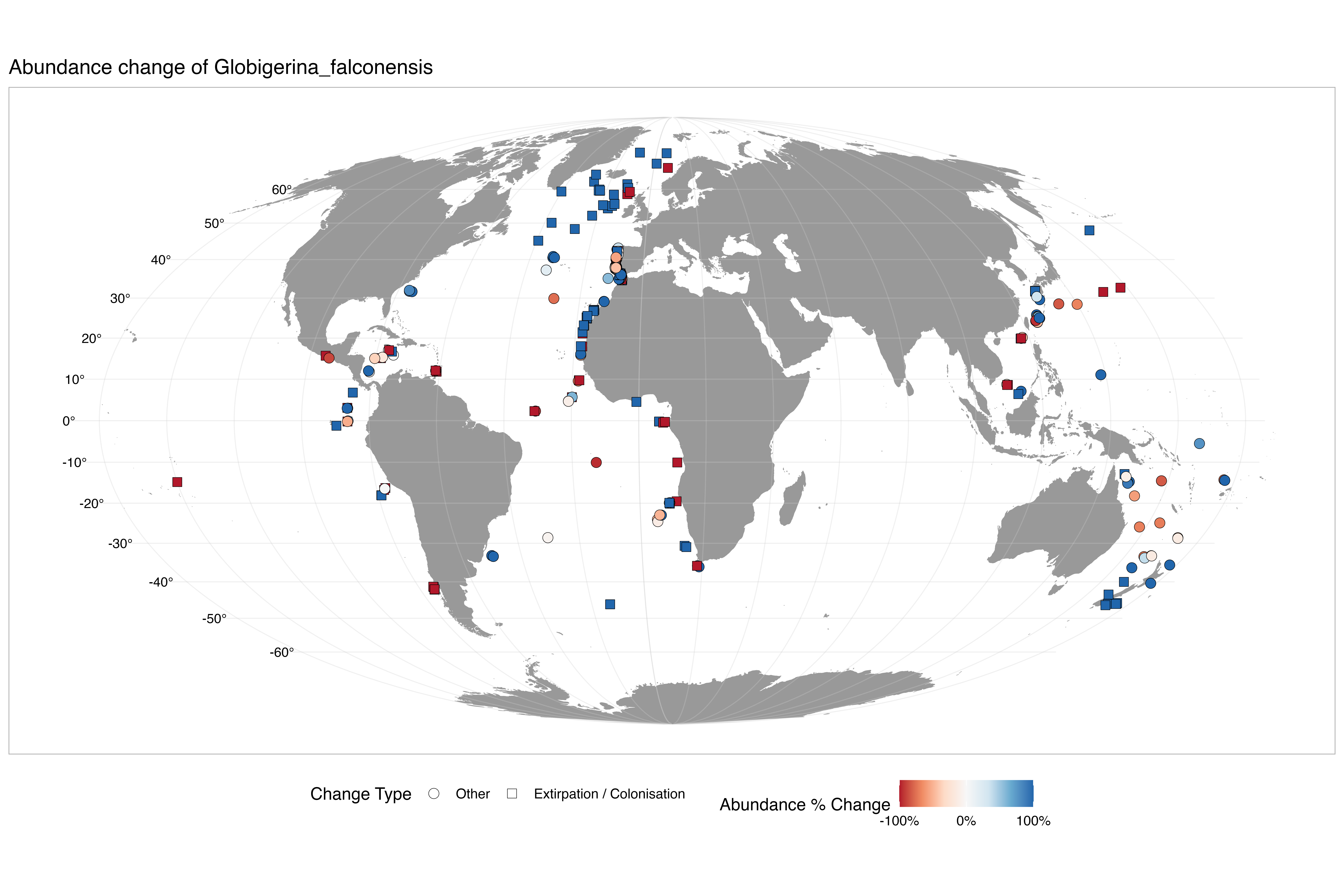

### abundance_change_Globigerinella_adamsi.png

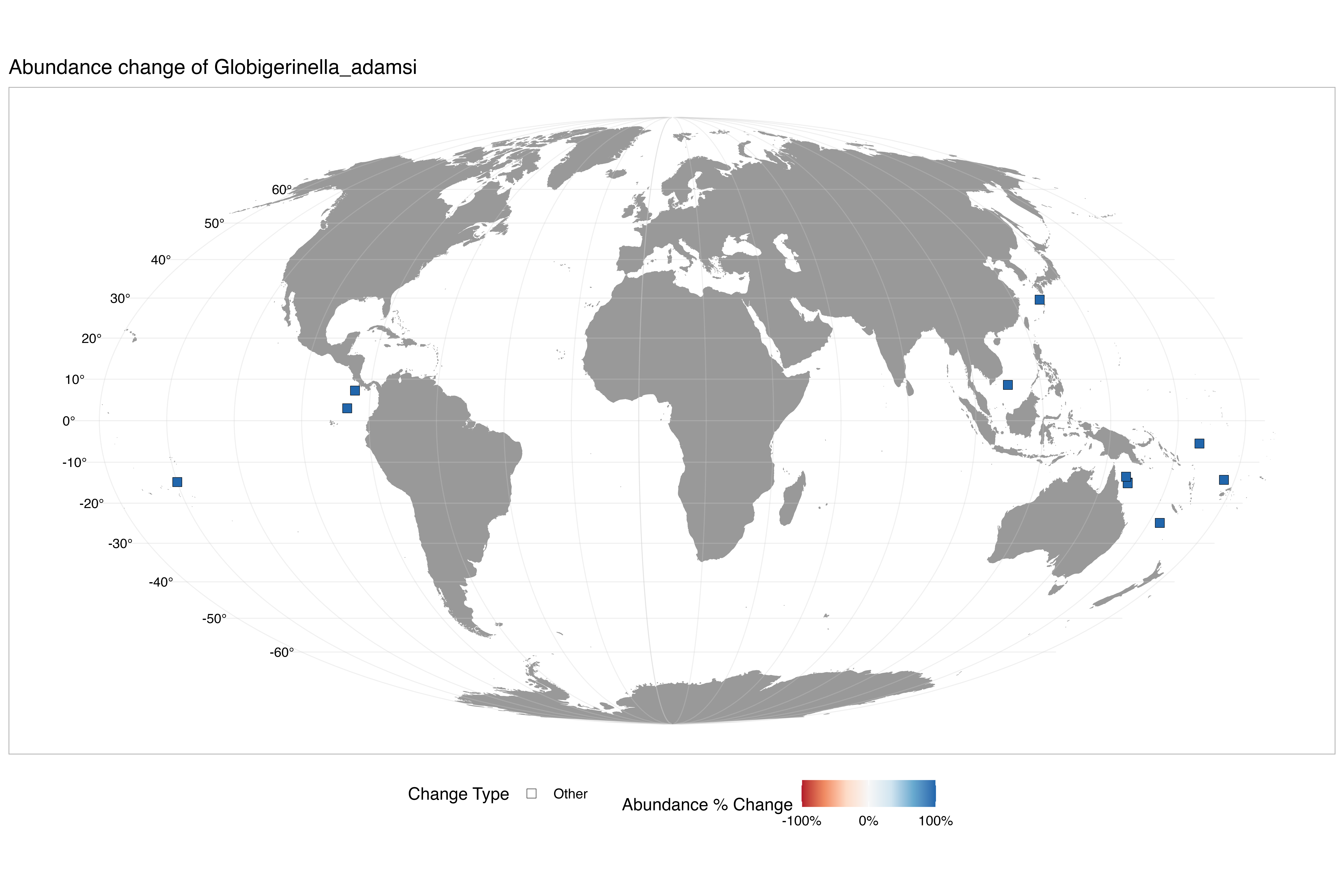

### abundance_change_Globigerinella_calida.png

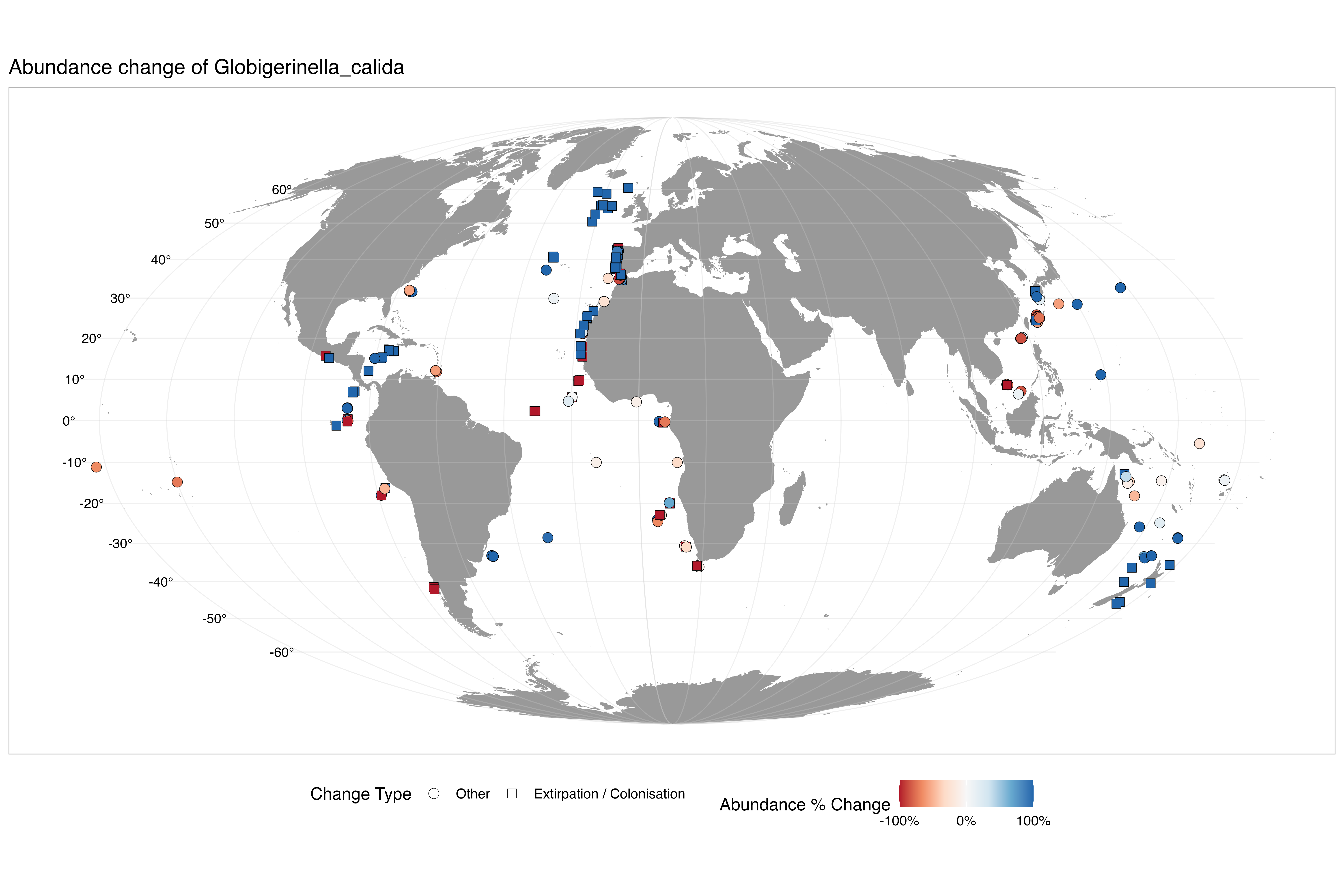

### abundance_change_Globigerinella_siphonifera.png

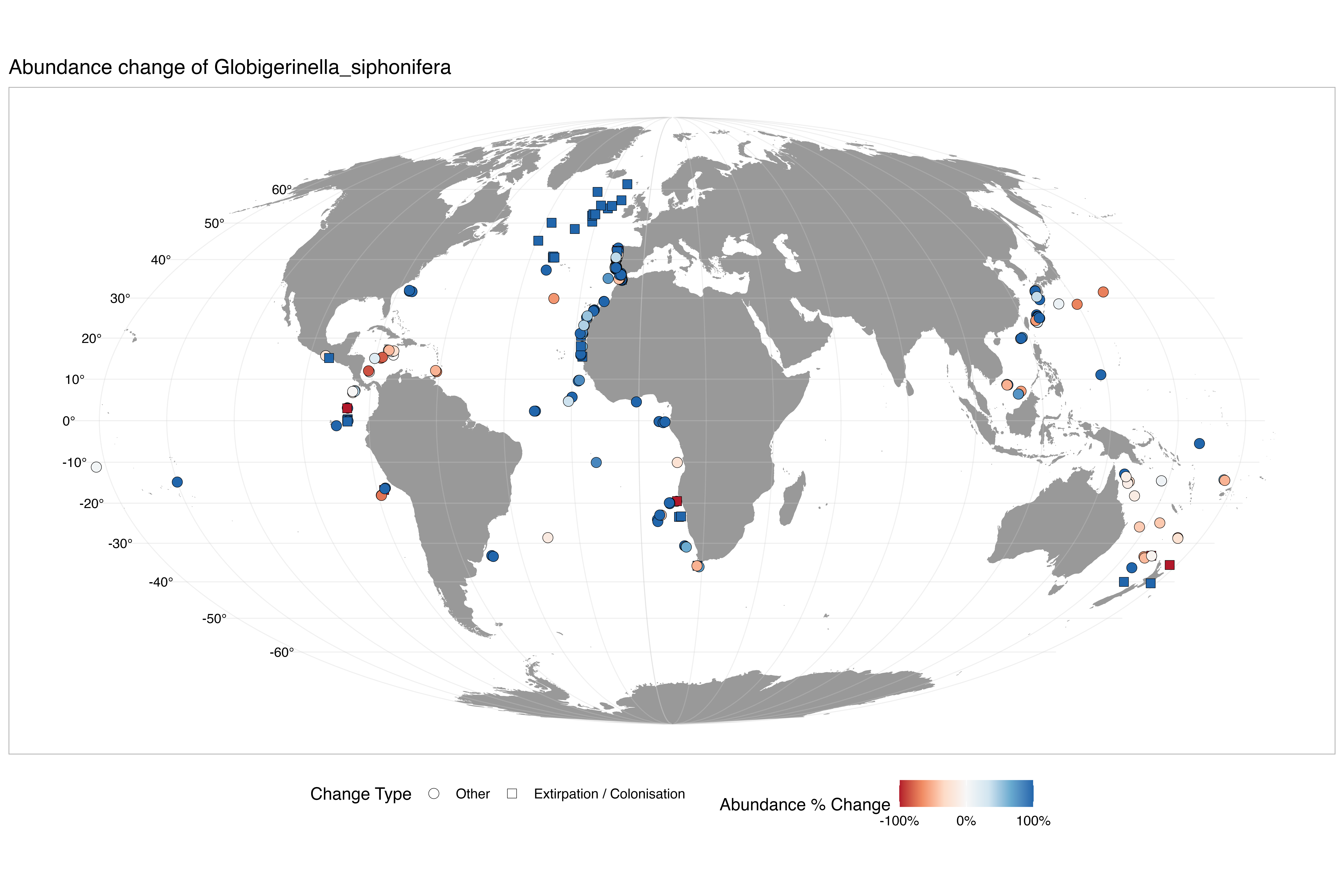

### abundance_change_Globigerinita_glutinata.png

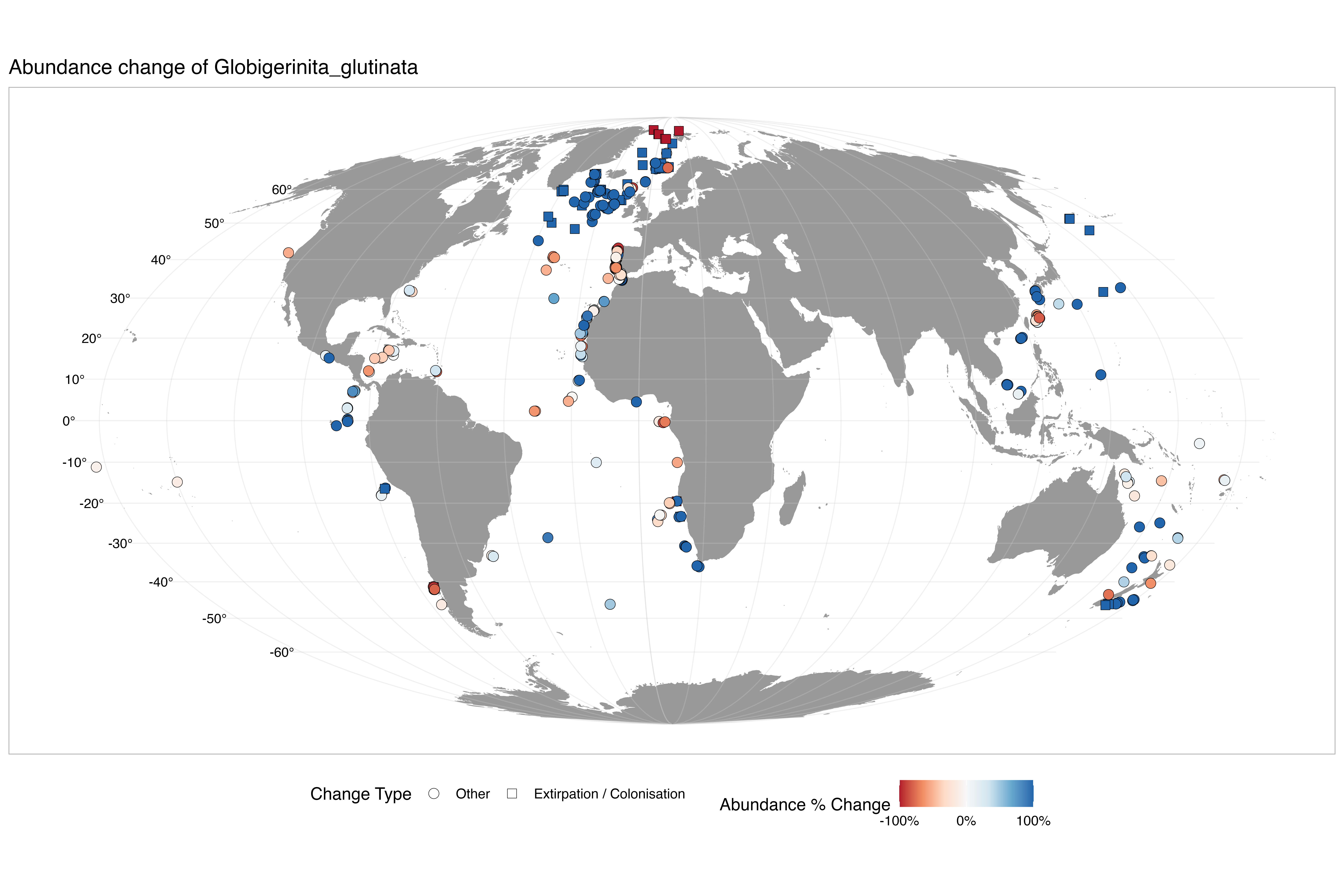

### abundance_change_Globigerinita_uvula.png

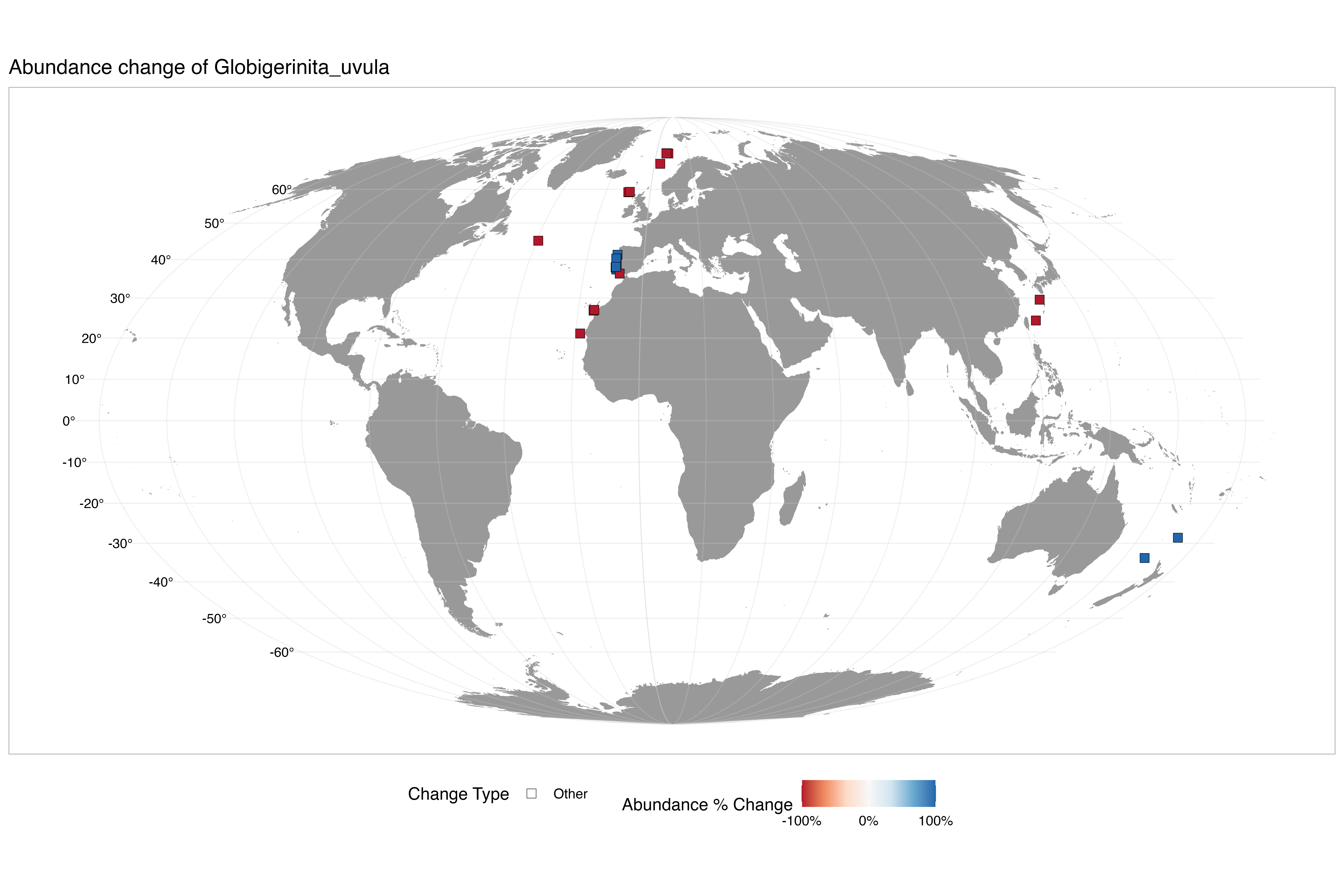

### abundance_change_Globigerinoides_conglobatus.png

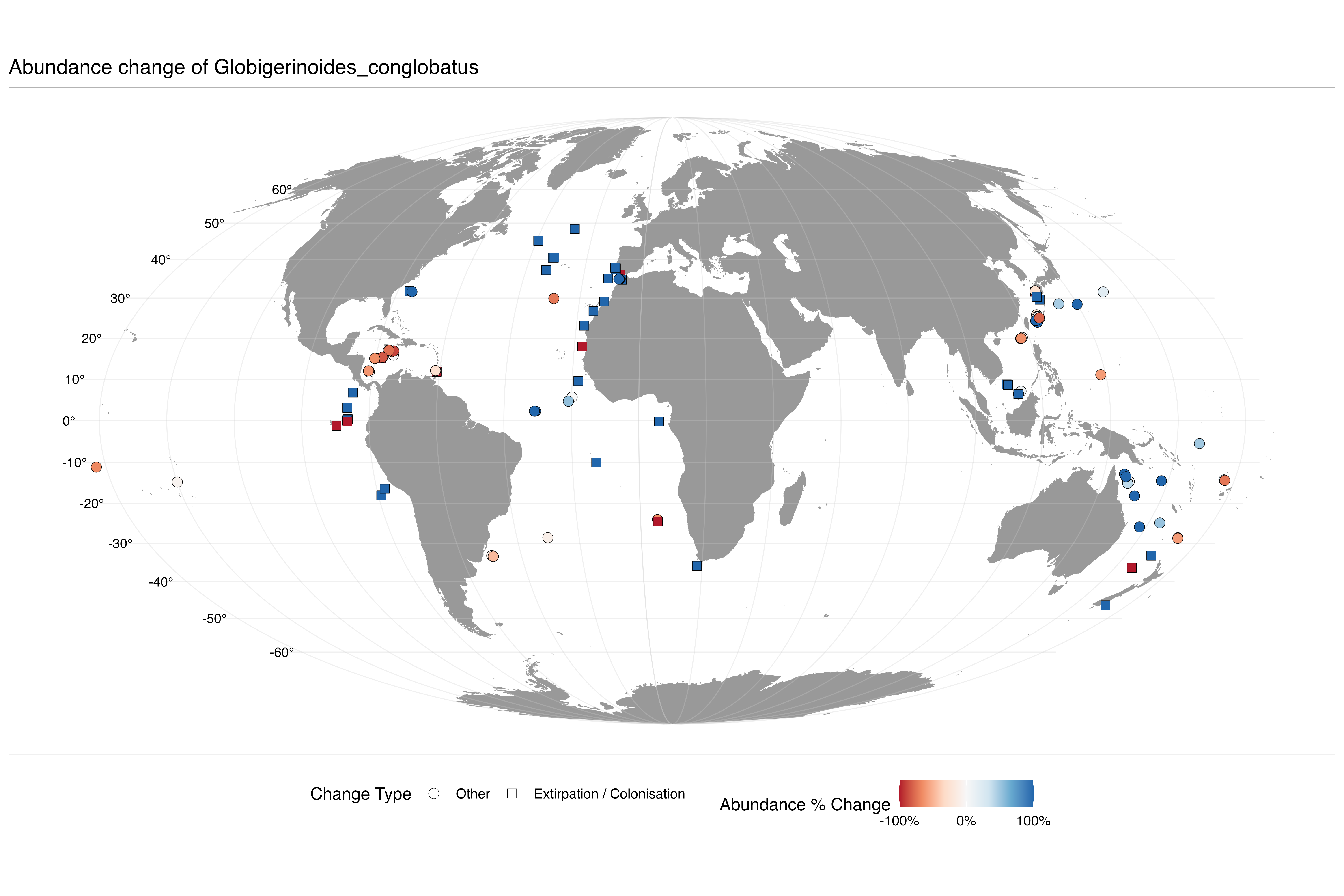

### abundance_change_Globigerinoides_ruber.png

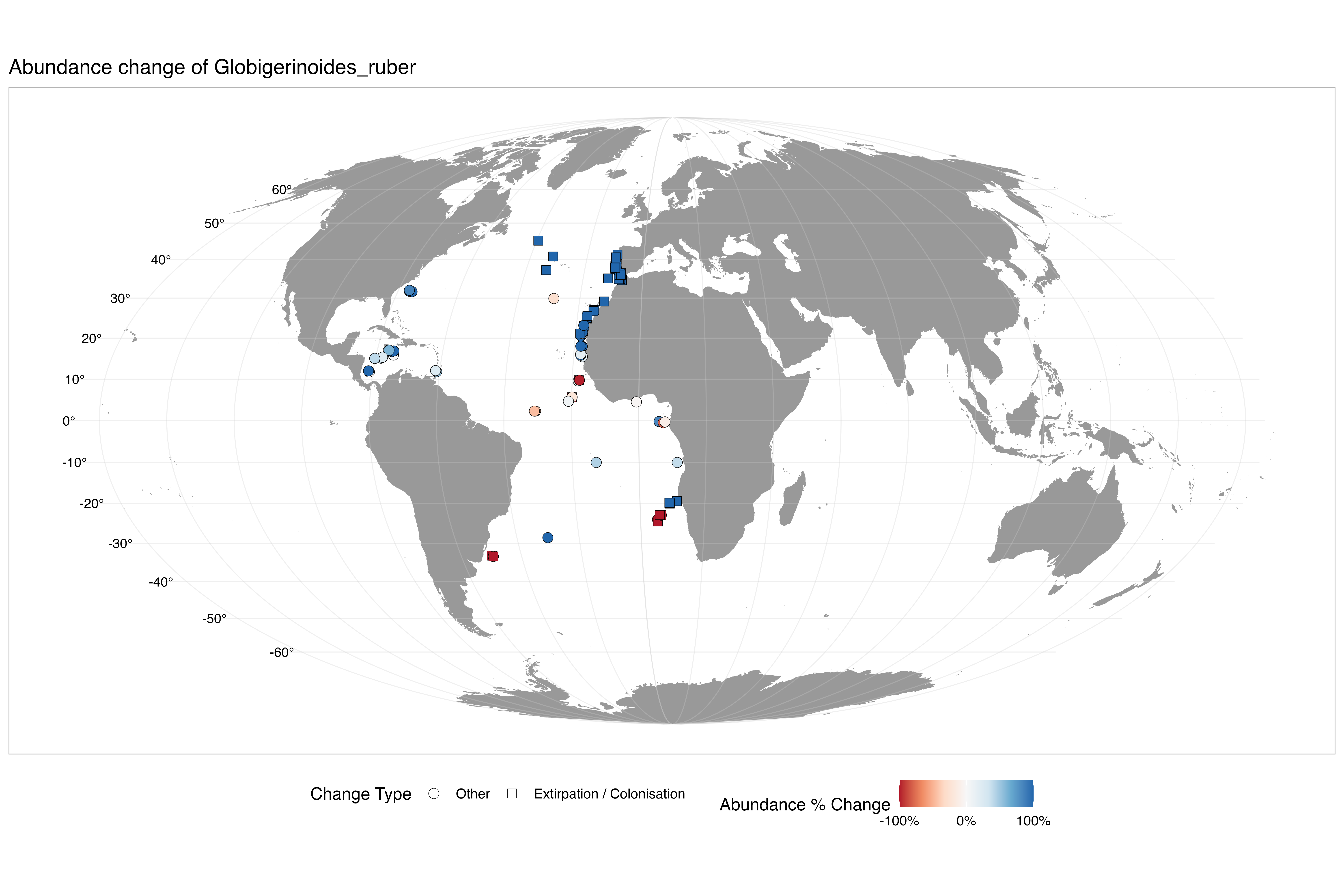

### abundance_change_Globigerinoides_tenellus.png

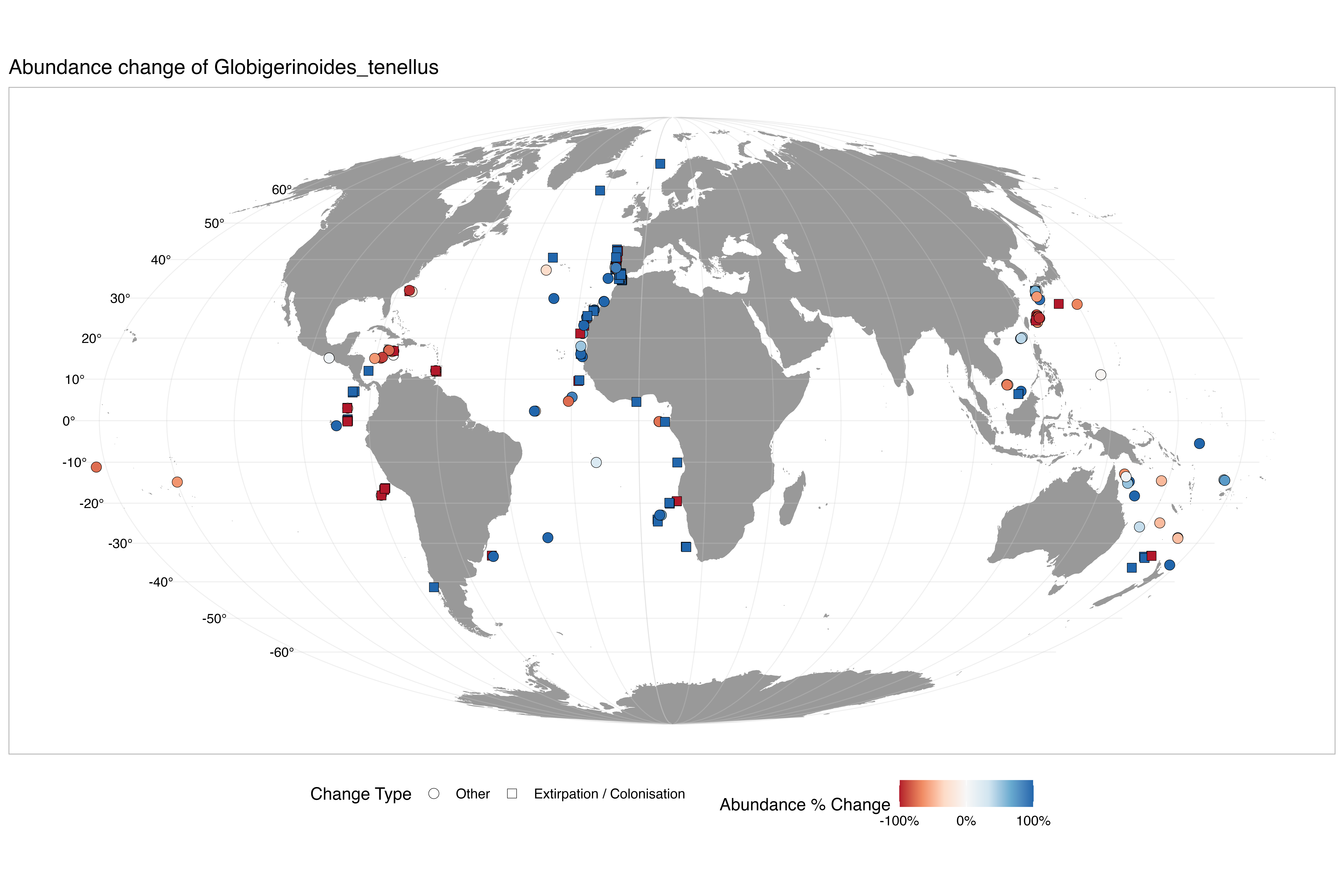

### abundance_change_Globigerinoides_white.png

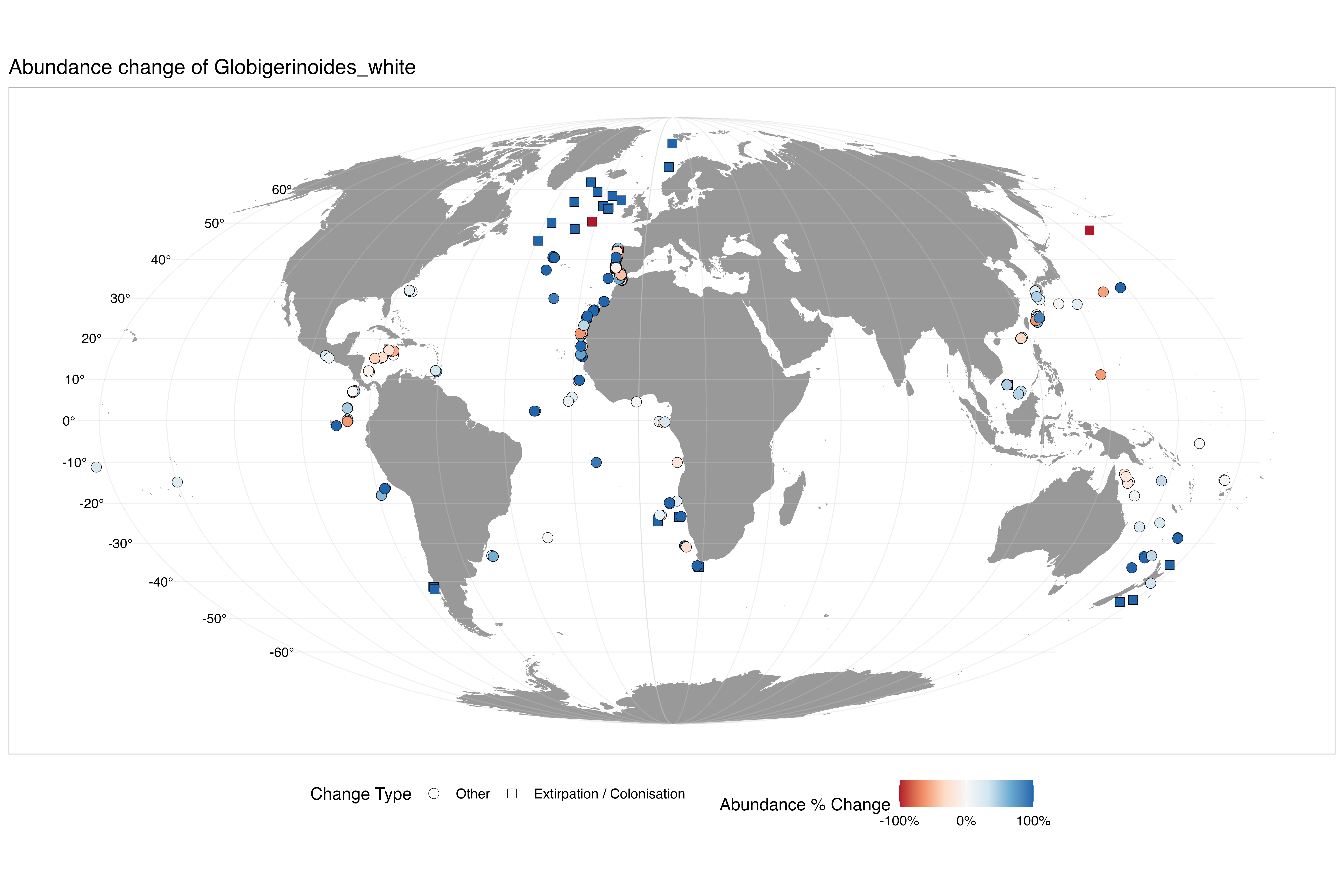

### abundance_change_Globoconella_inflata.png

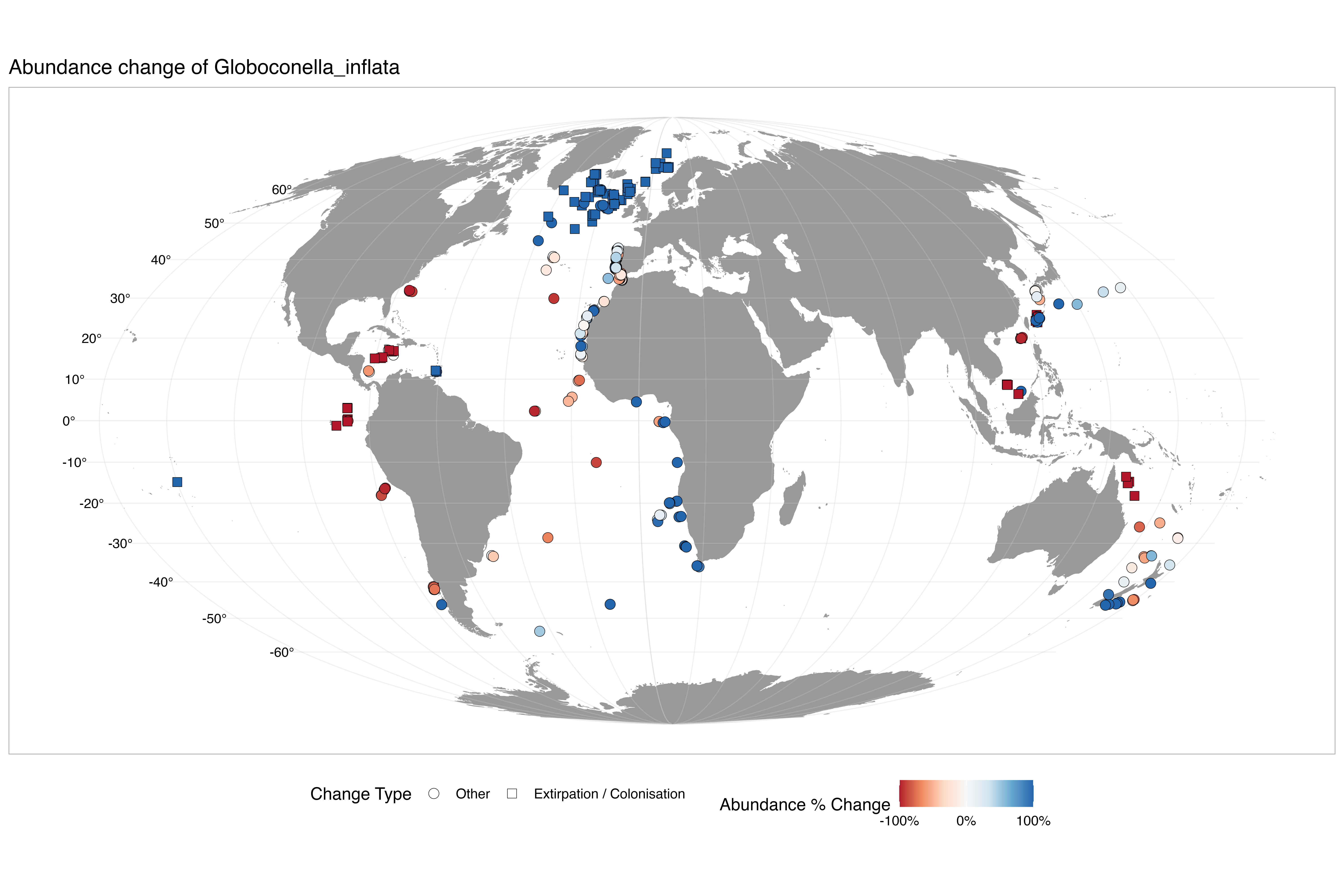

### abundance_change_Globoquadrina_conglomerata.png

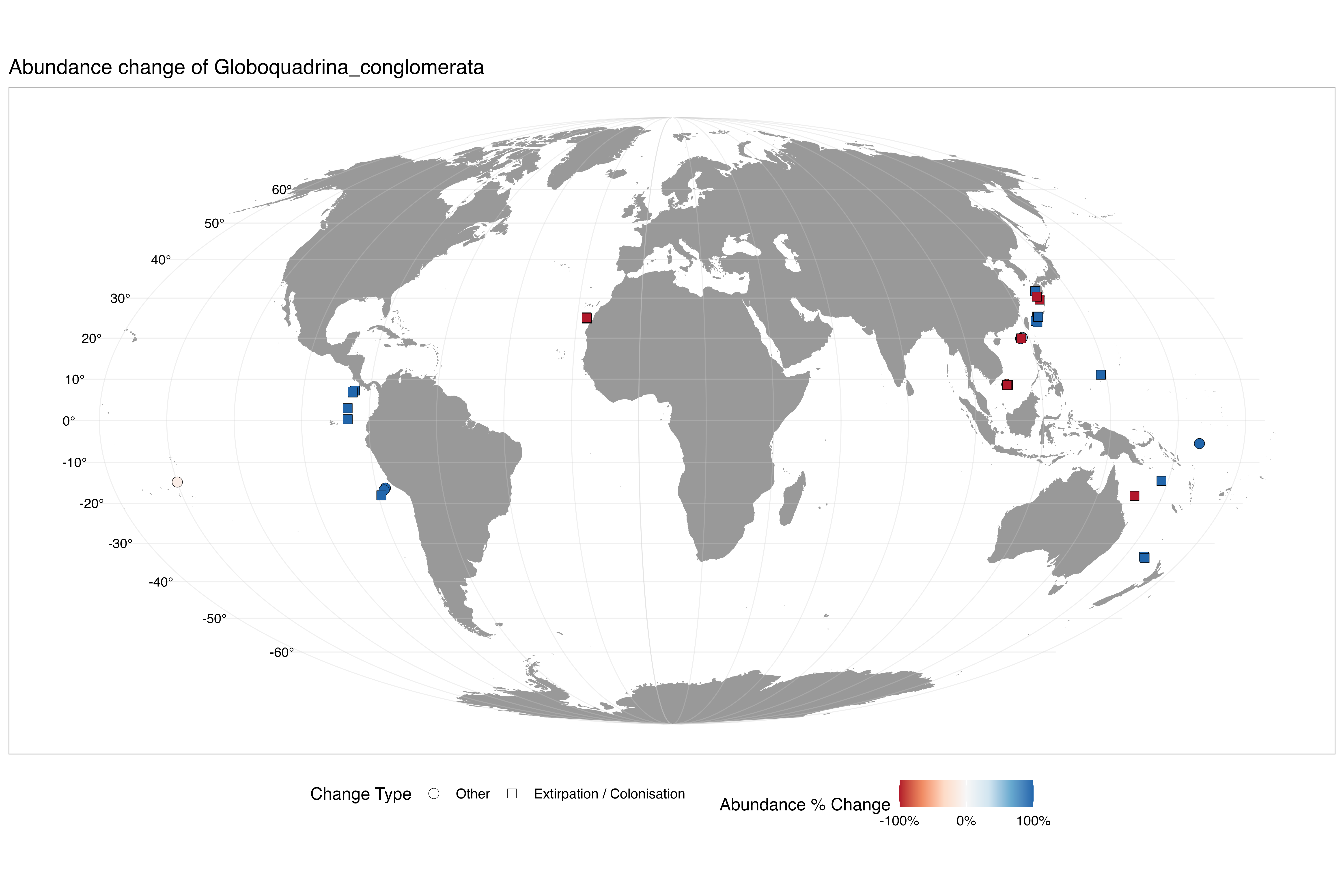

### abundance_change_Globorotalia_crassaformis.png

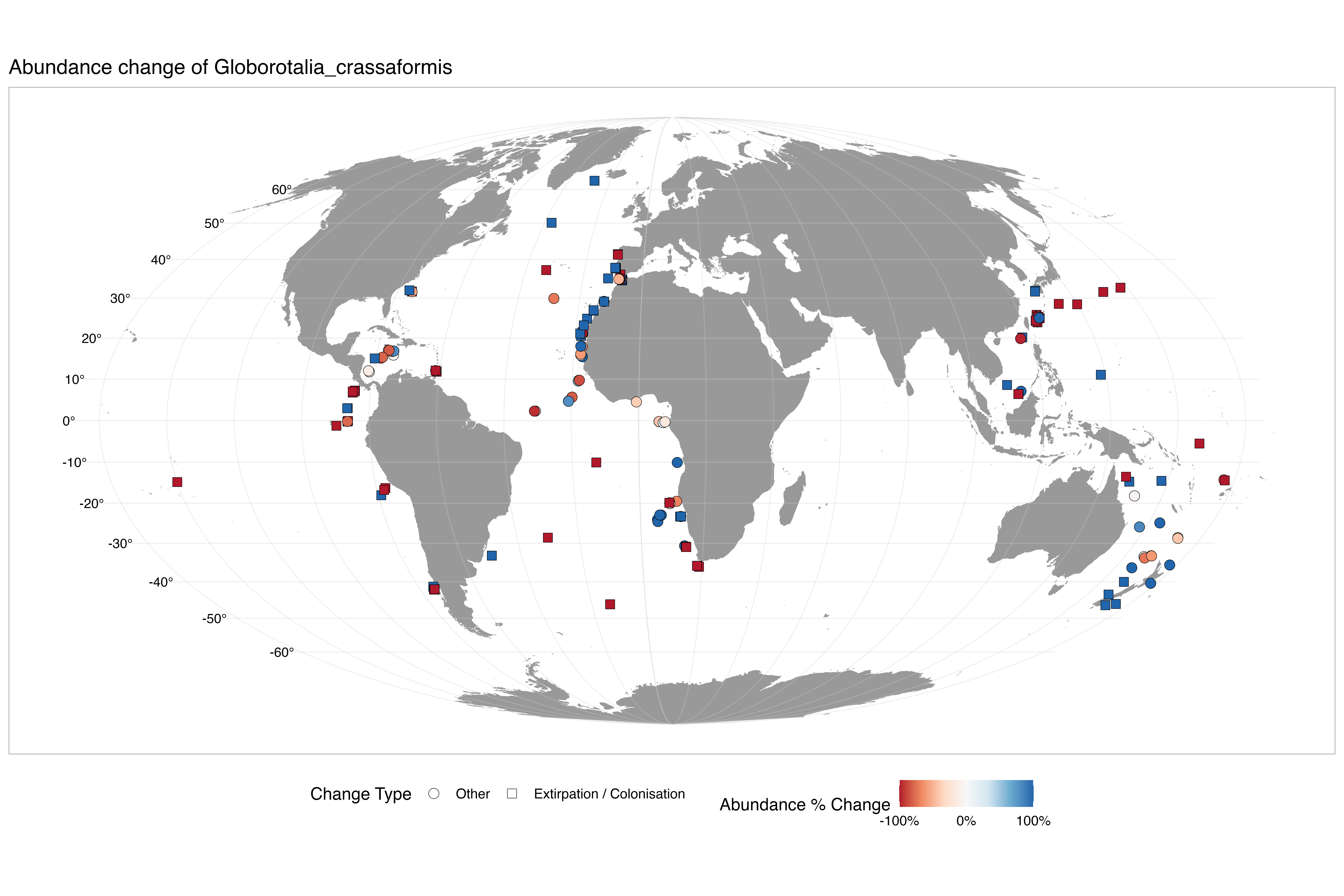

### abundance_change_Globorotalia_hirsuta.png

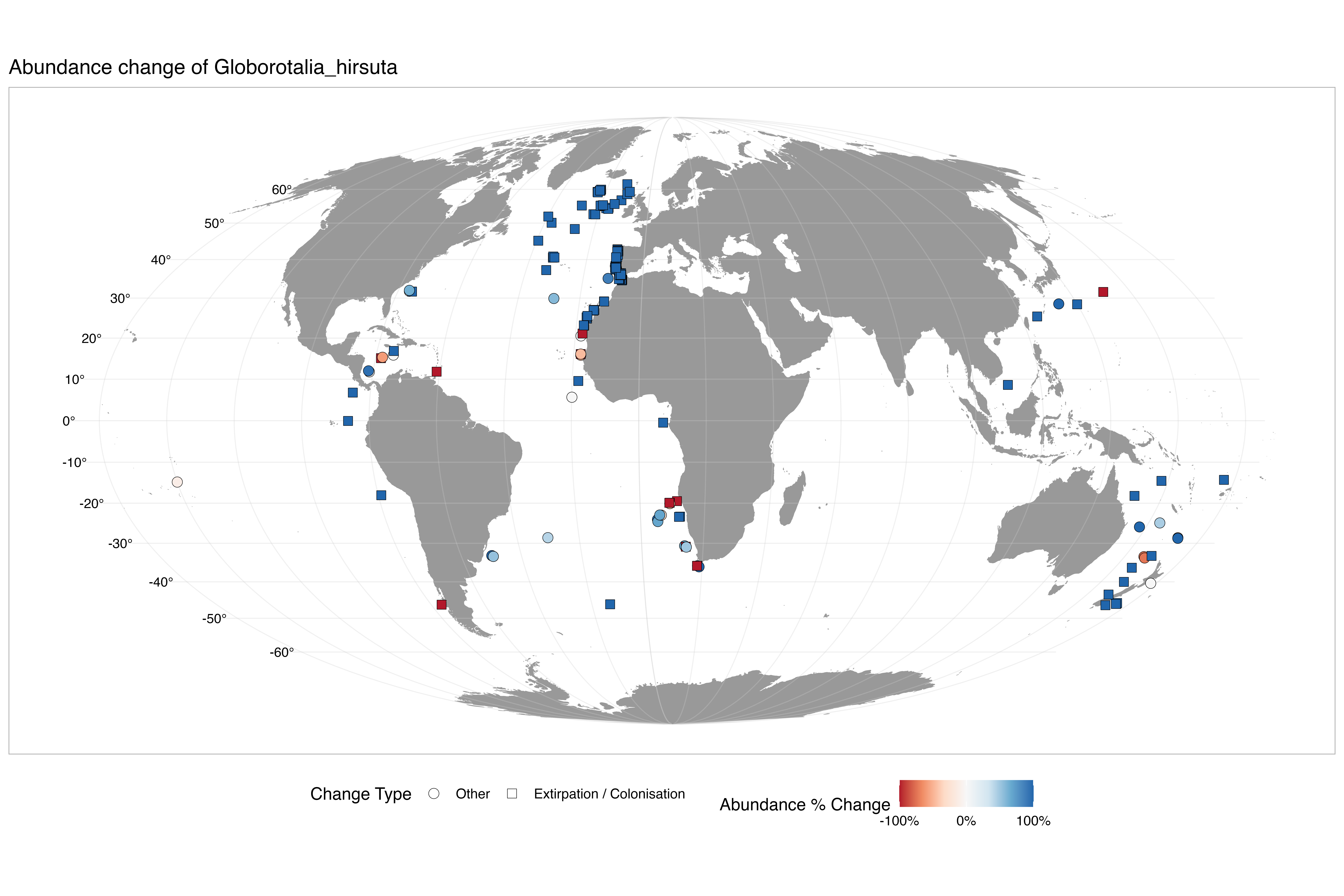

### abundance_change_Globorotalia_menardii.png

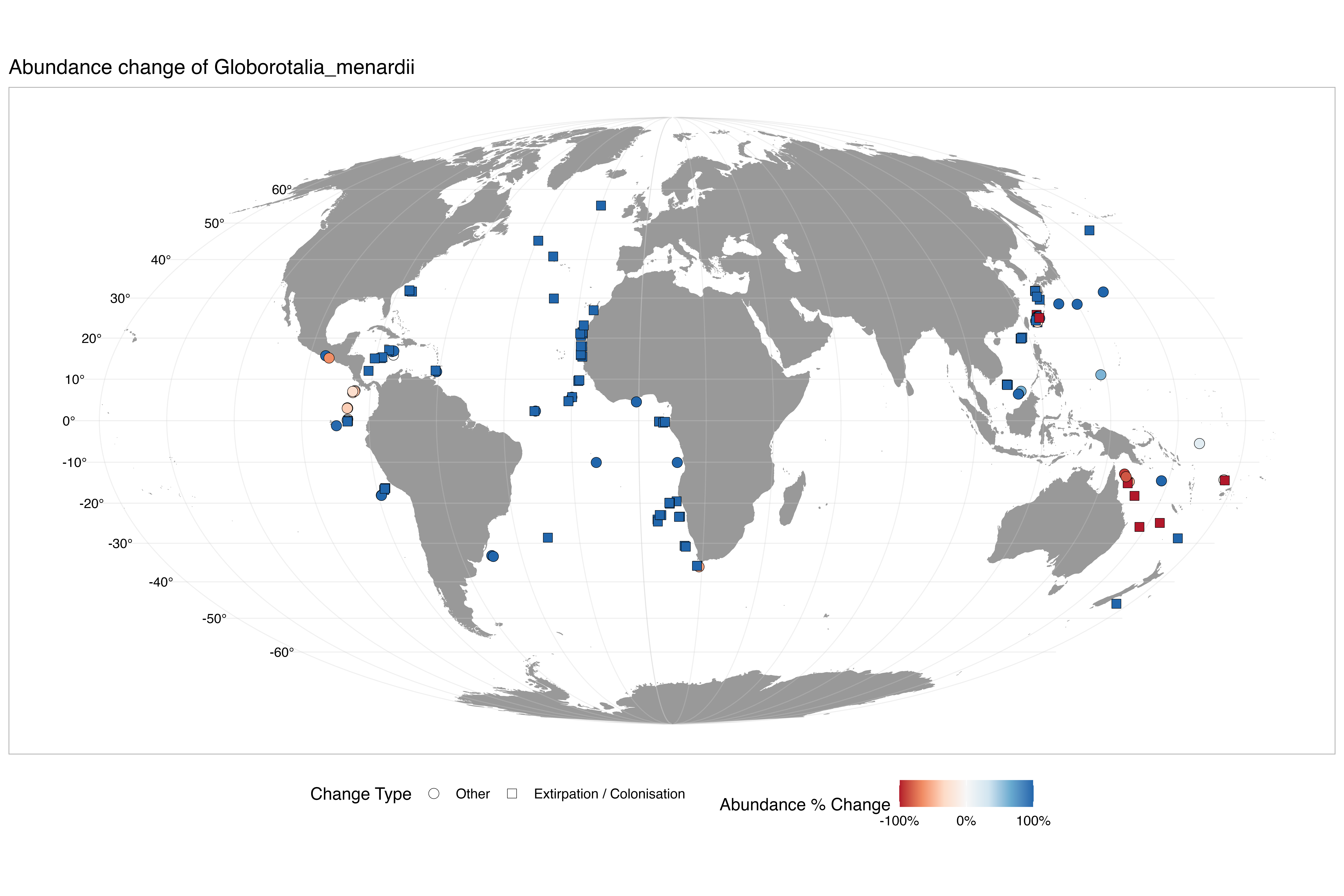

### abundance_change_Globorotalia_scitula.png

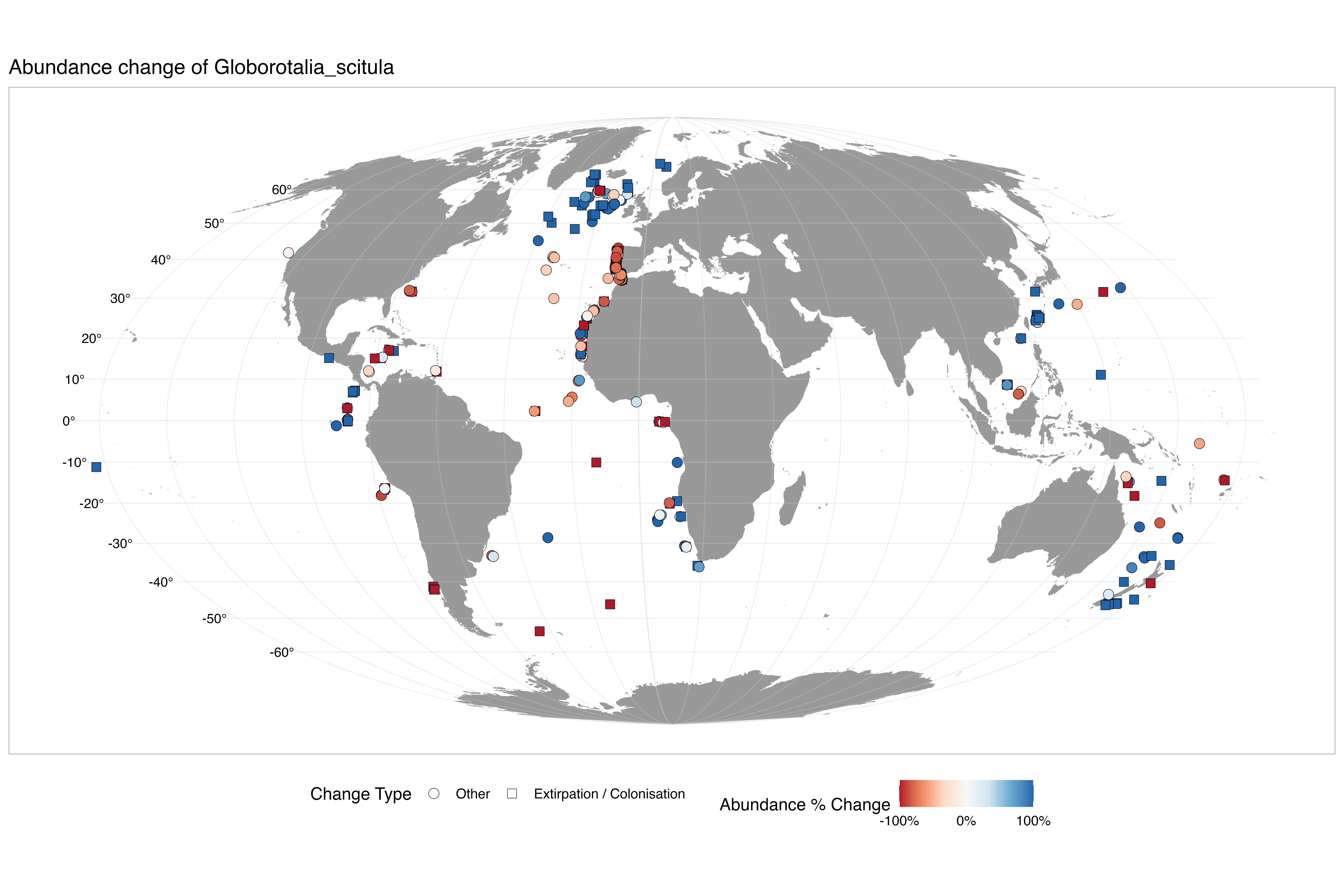

### abundance_change_Globorotalia_theyeri.png

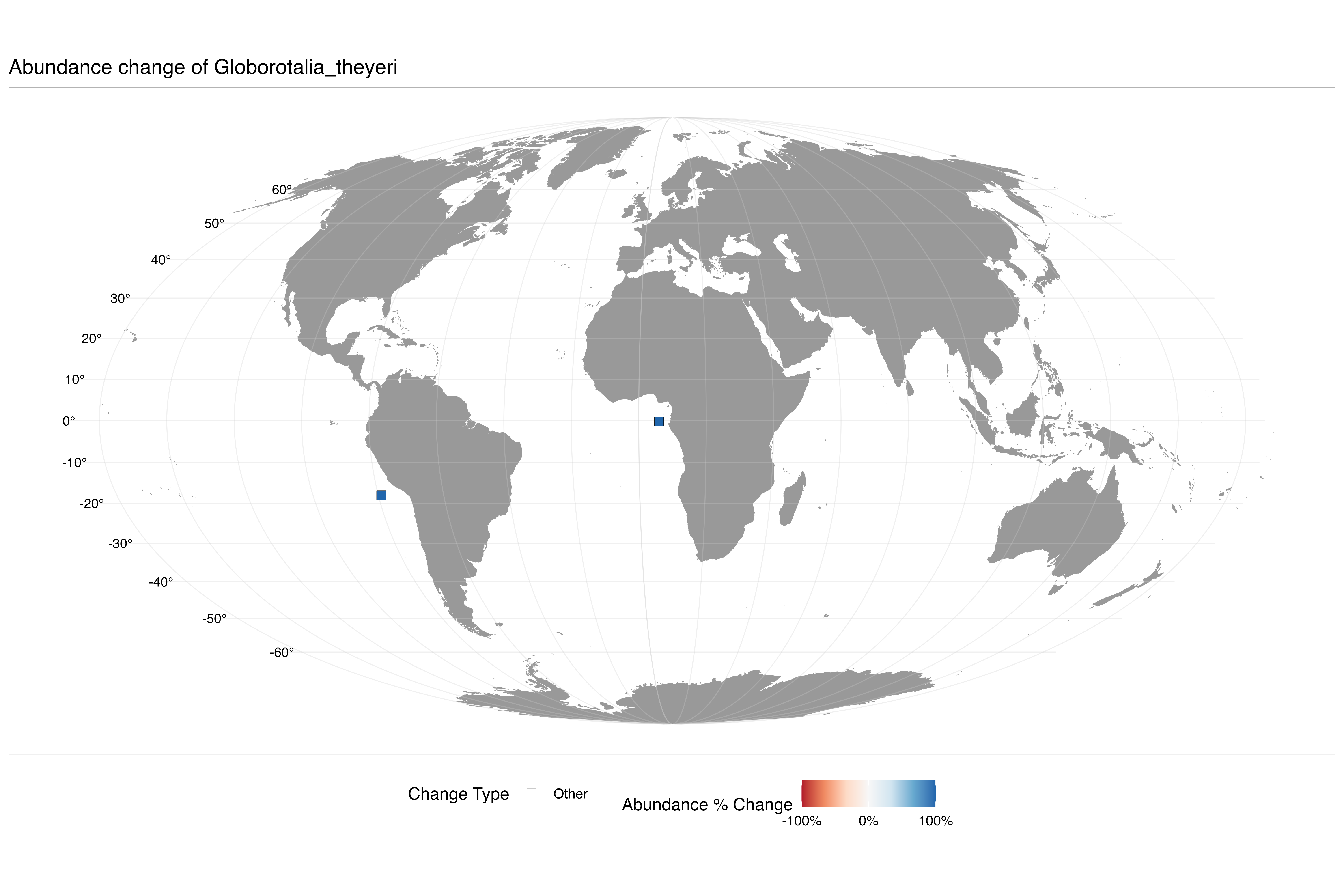

### abundance_change_Globorotalia_truncatulinoides.png

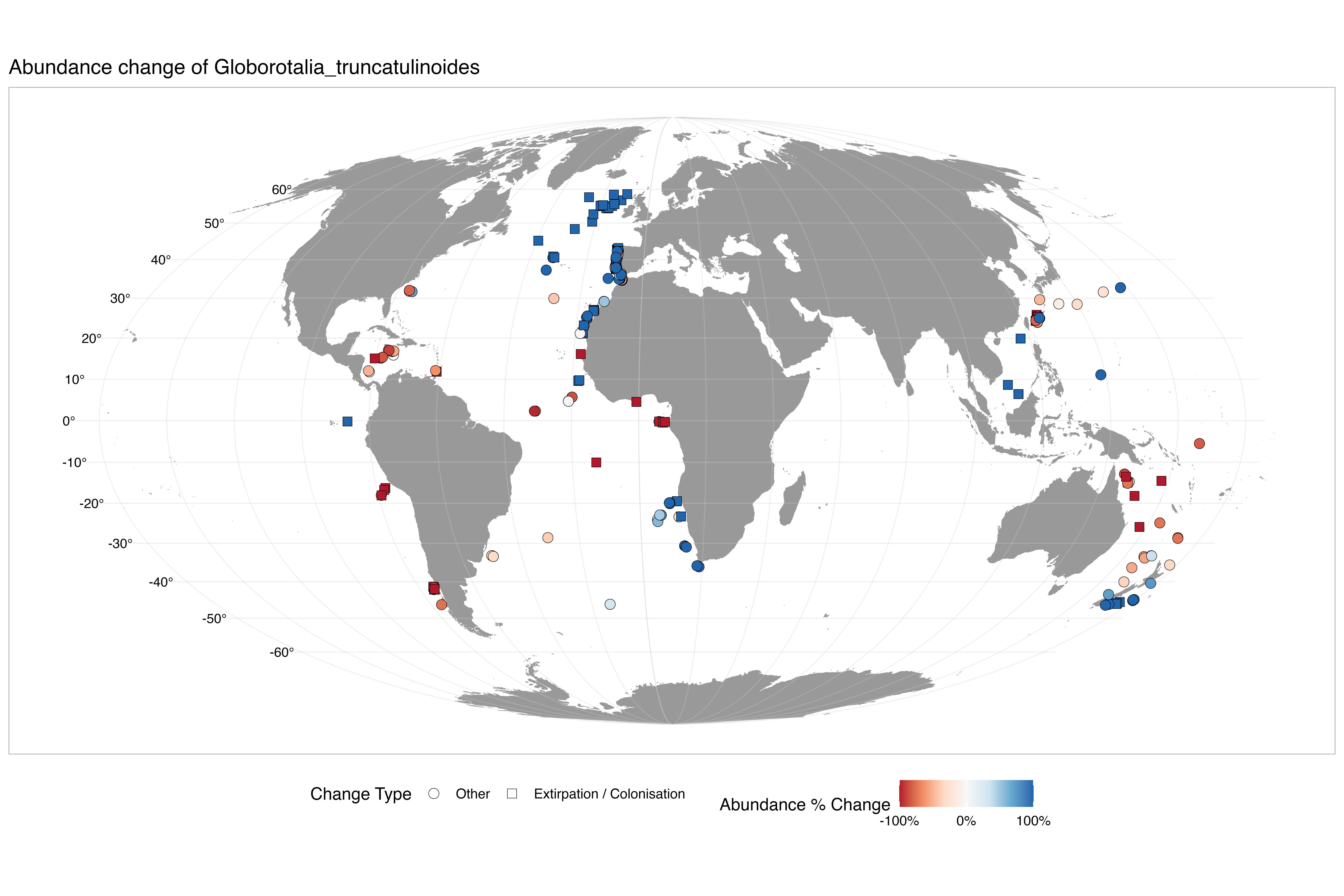

### abundance_change_Globorotalia_tumida.png

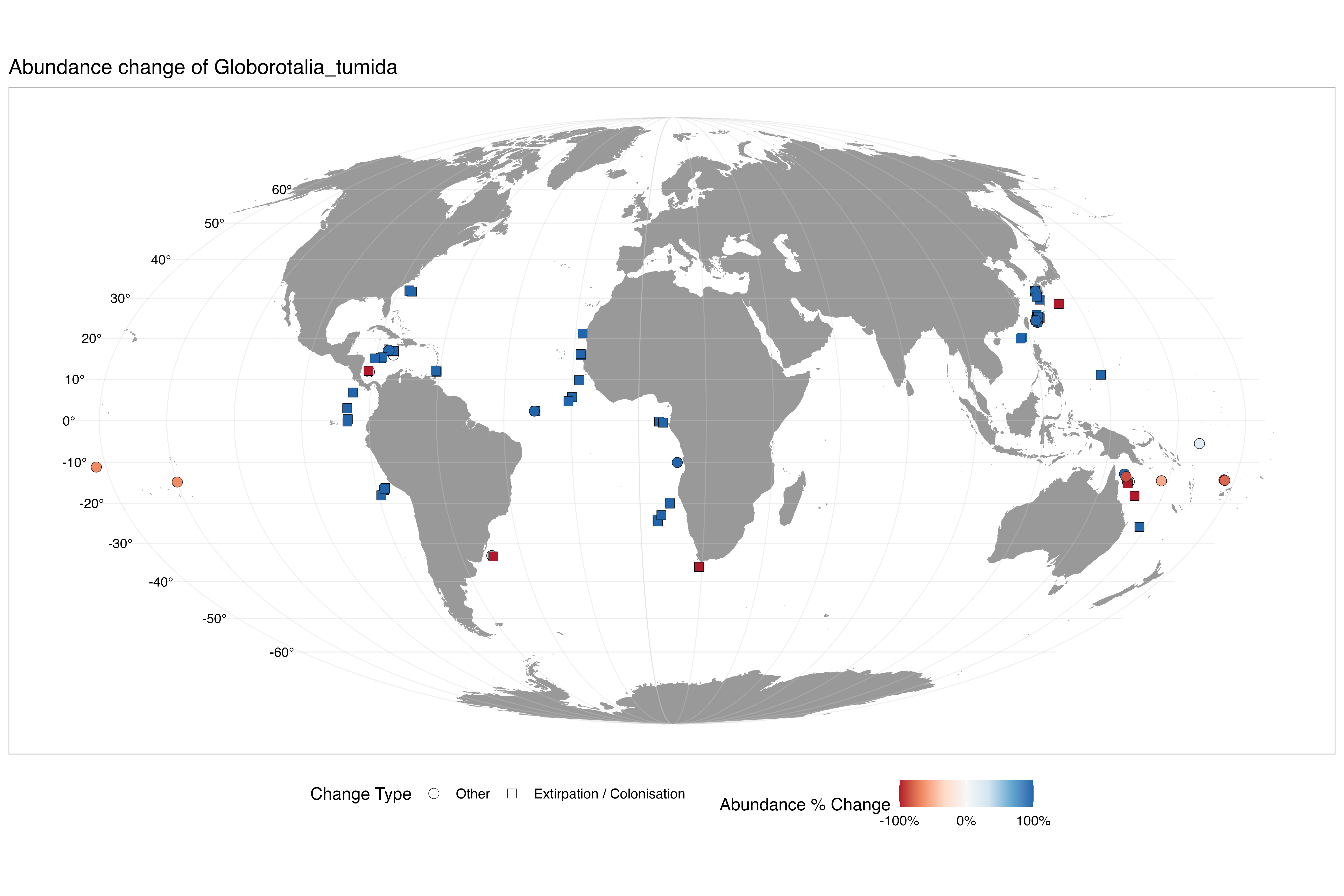

### abundance_change_Globorotalia_ungulata.png

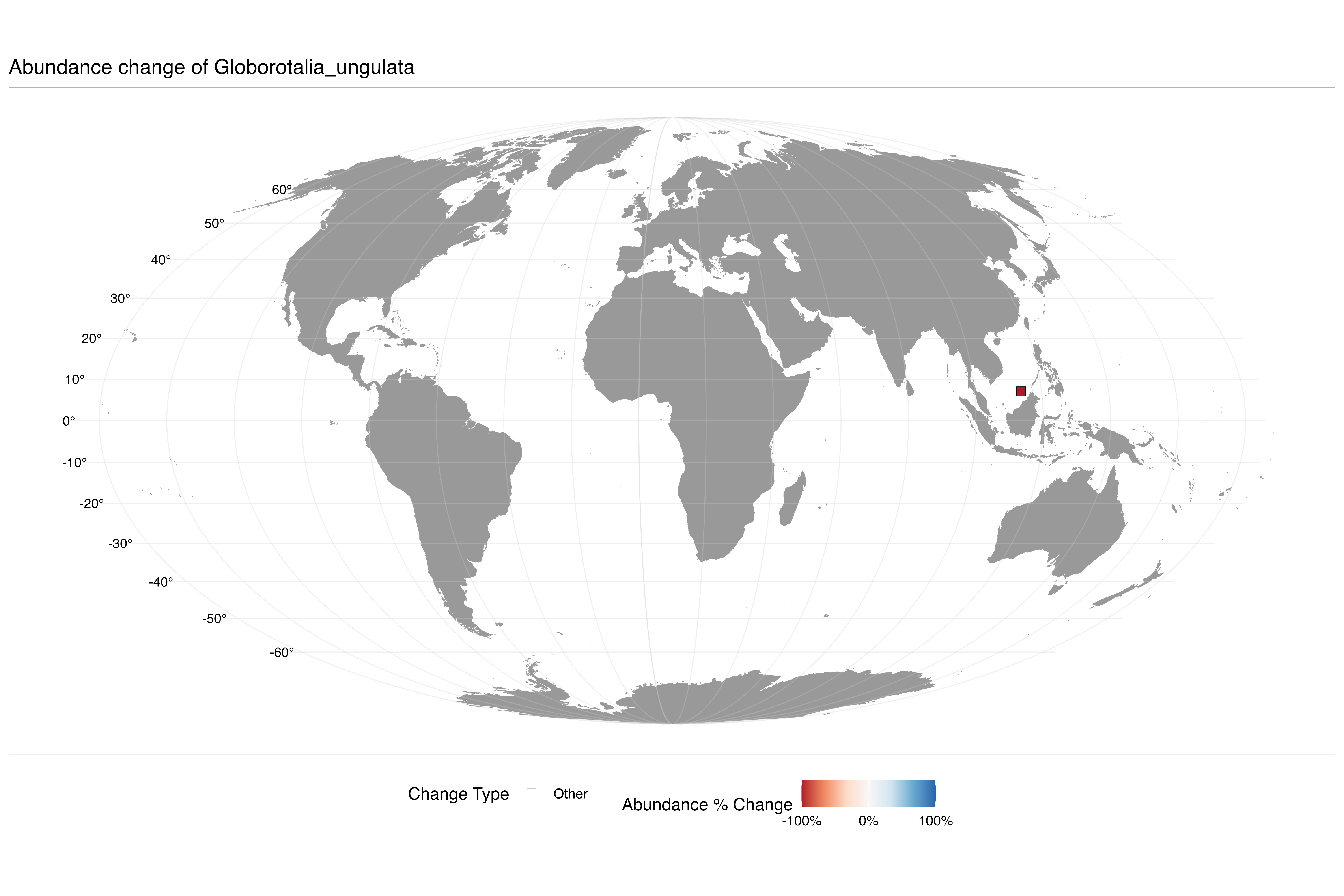

### abundance_change_Globorotaloides_hexagonus.png

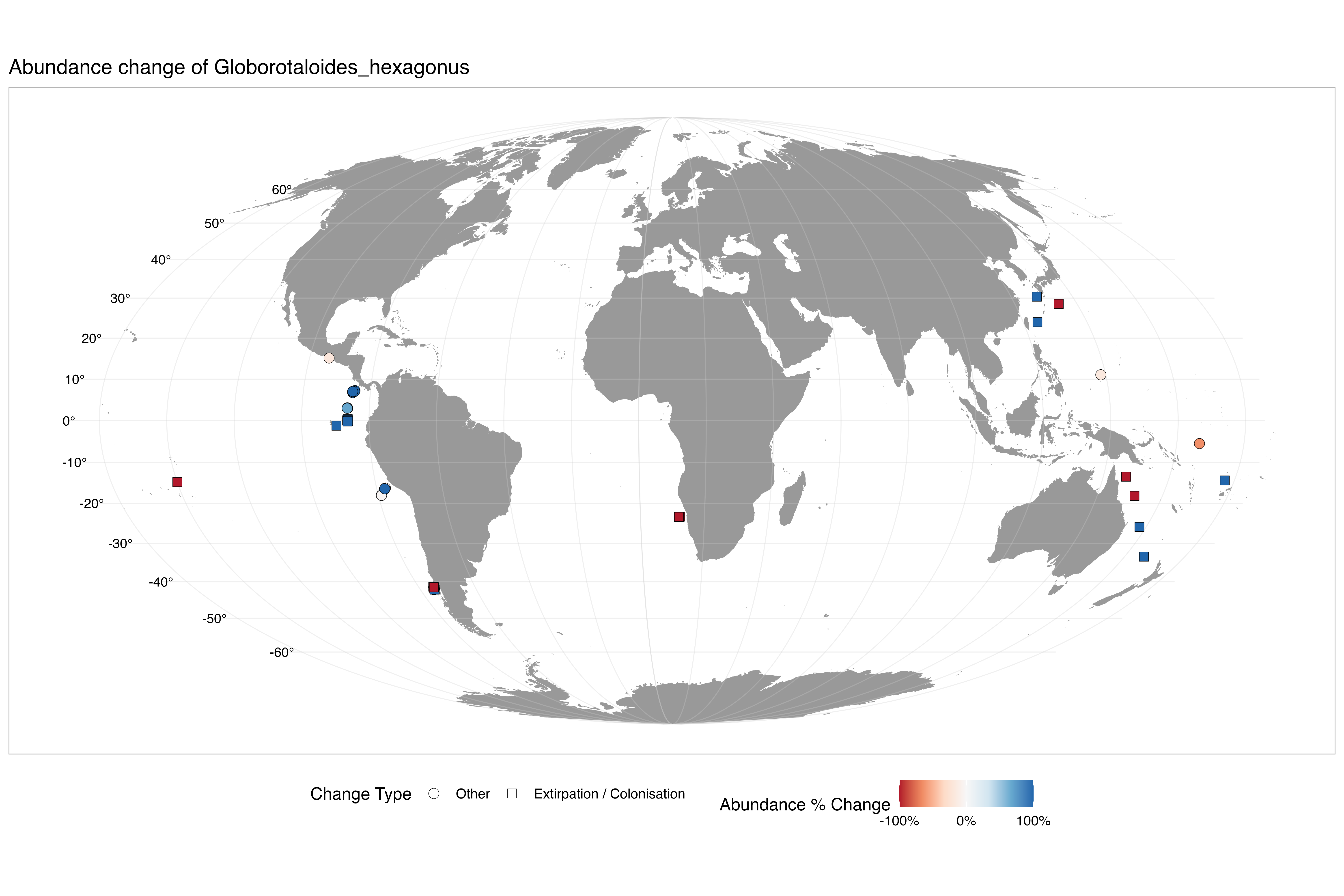

### abundance_change_Globoturborotalita_rubescens.png

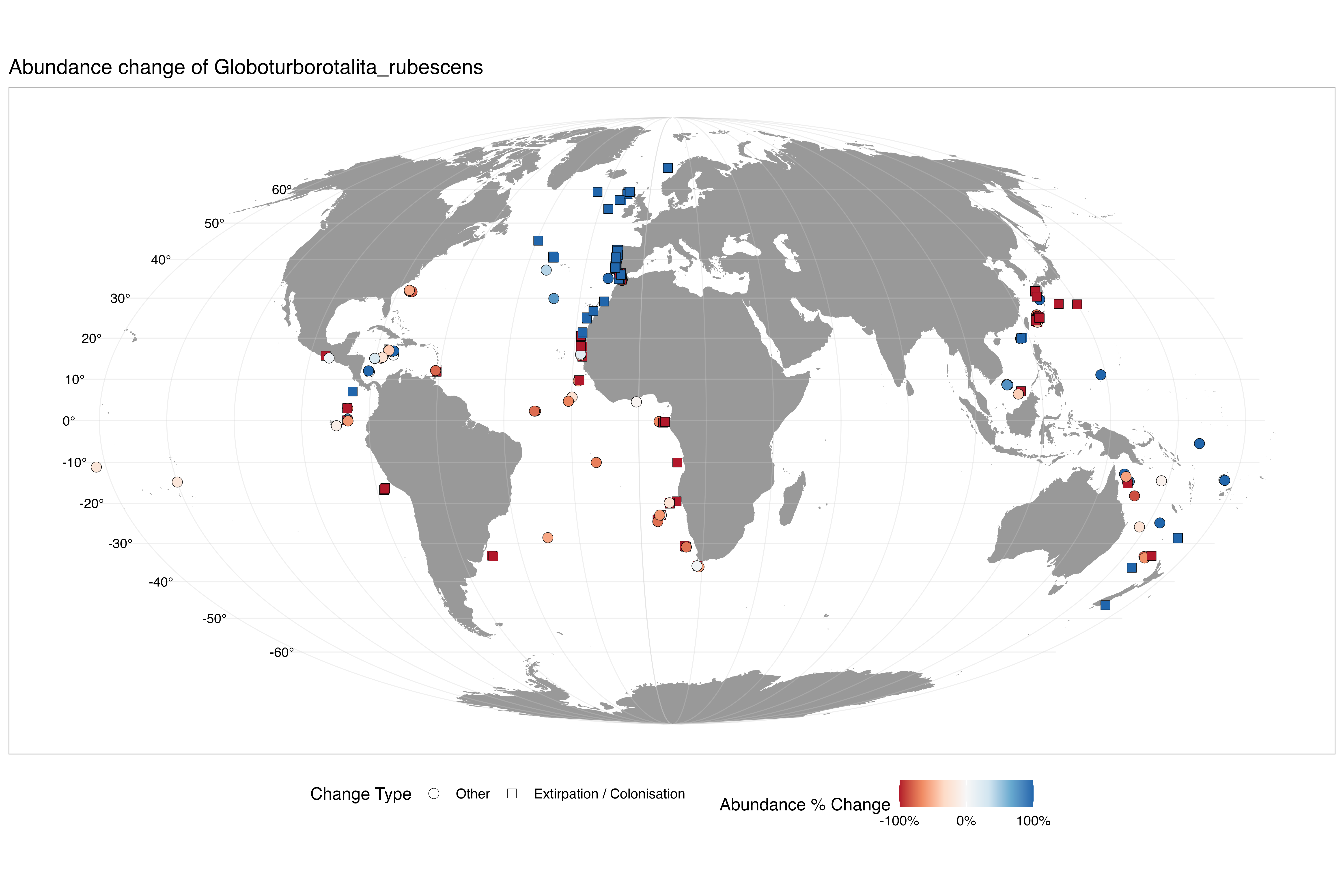

### abundance_change_Neogloboquadrina_dutertrei.png

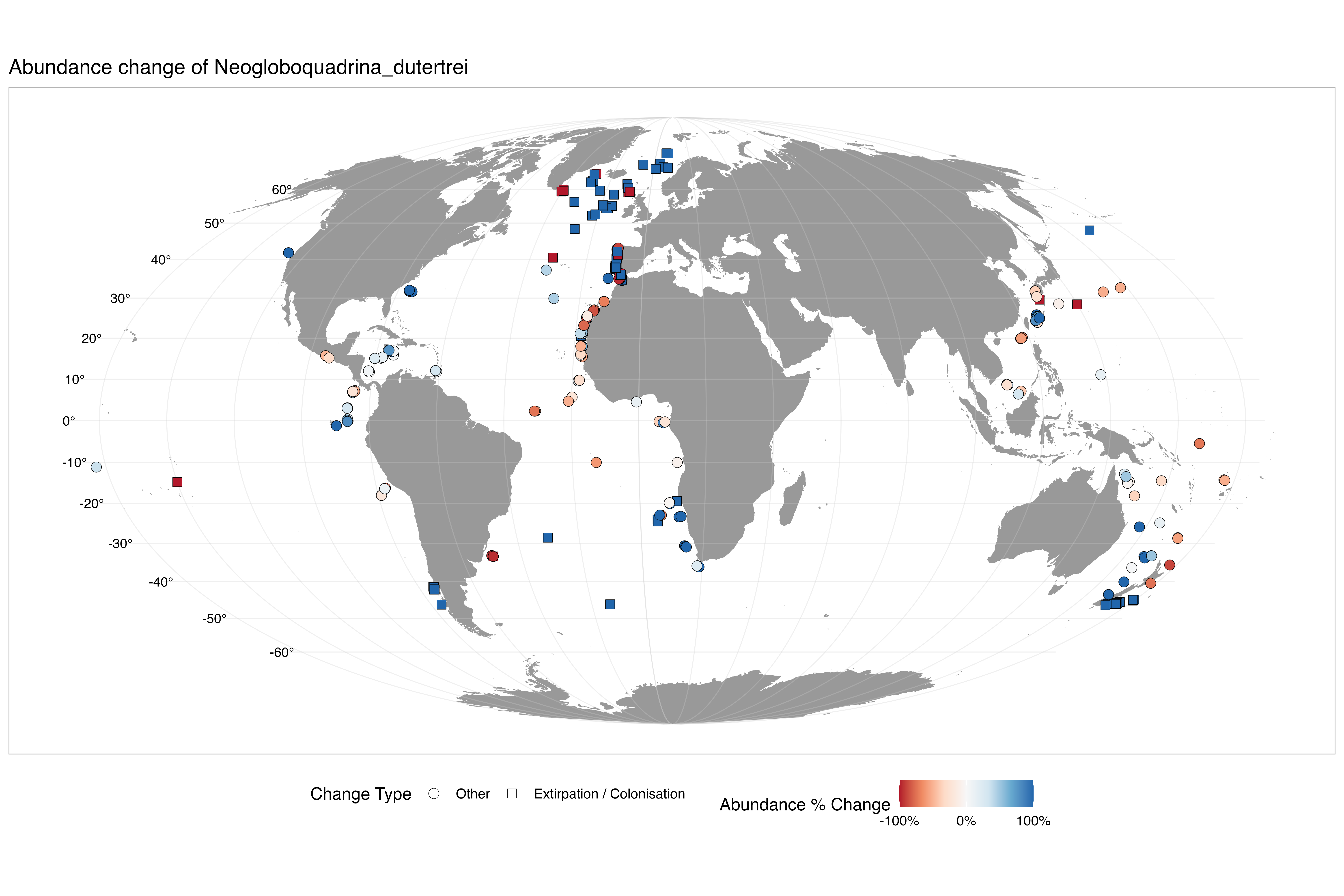

### abundance_change_Neogloboquadrina_incompta.png

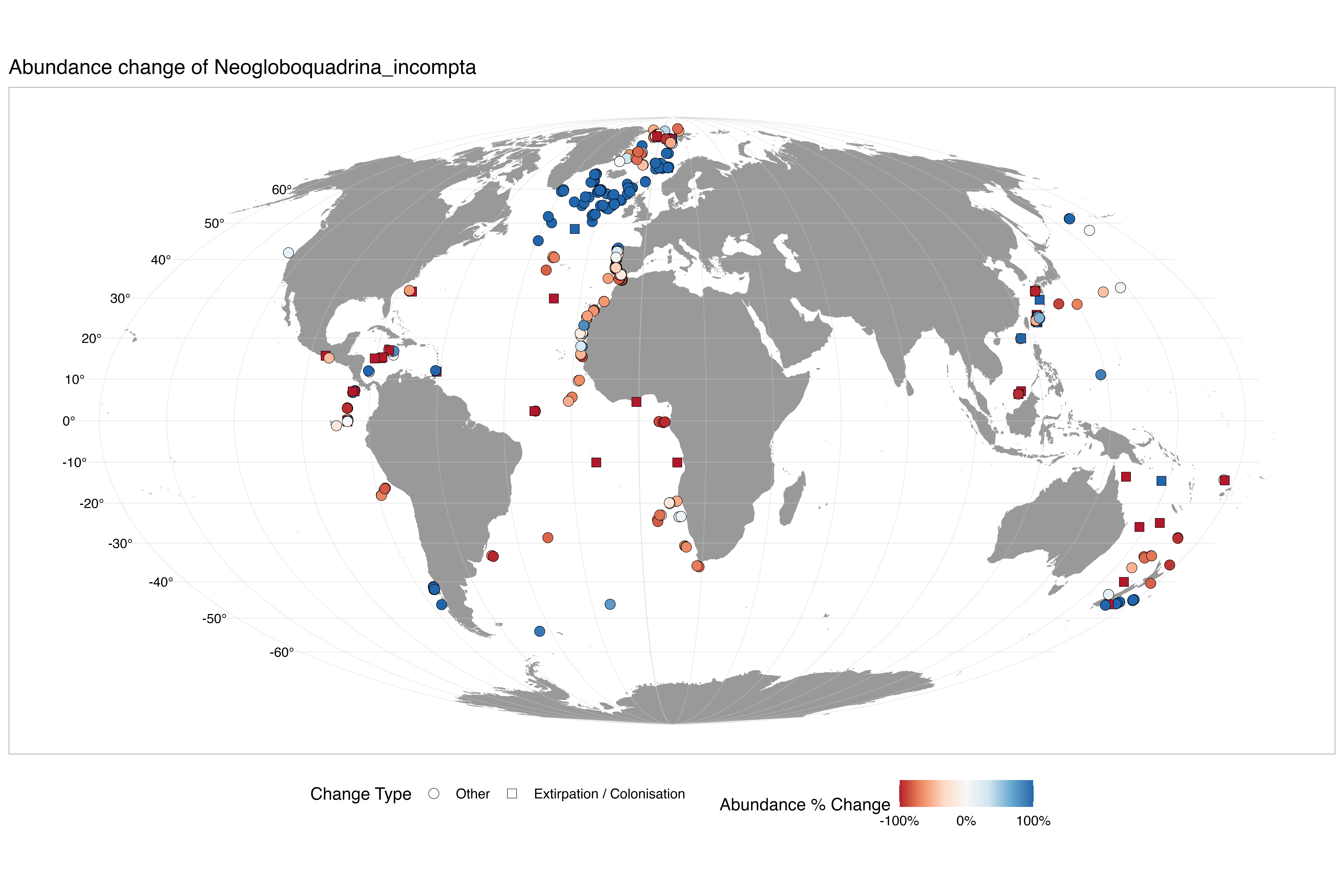

### abundance_change_Neogloboquadrina_pachyderma.png

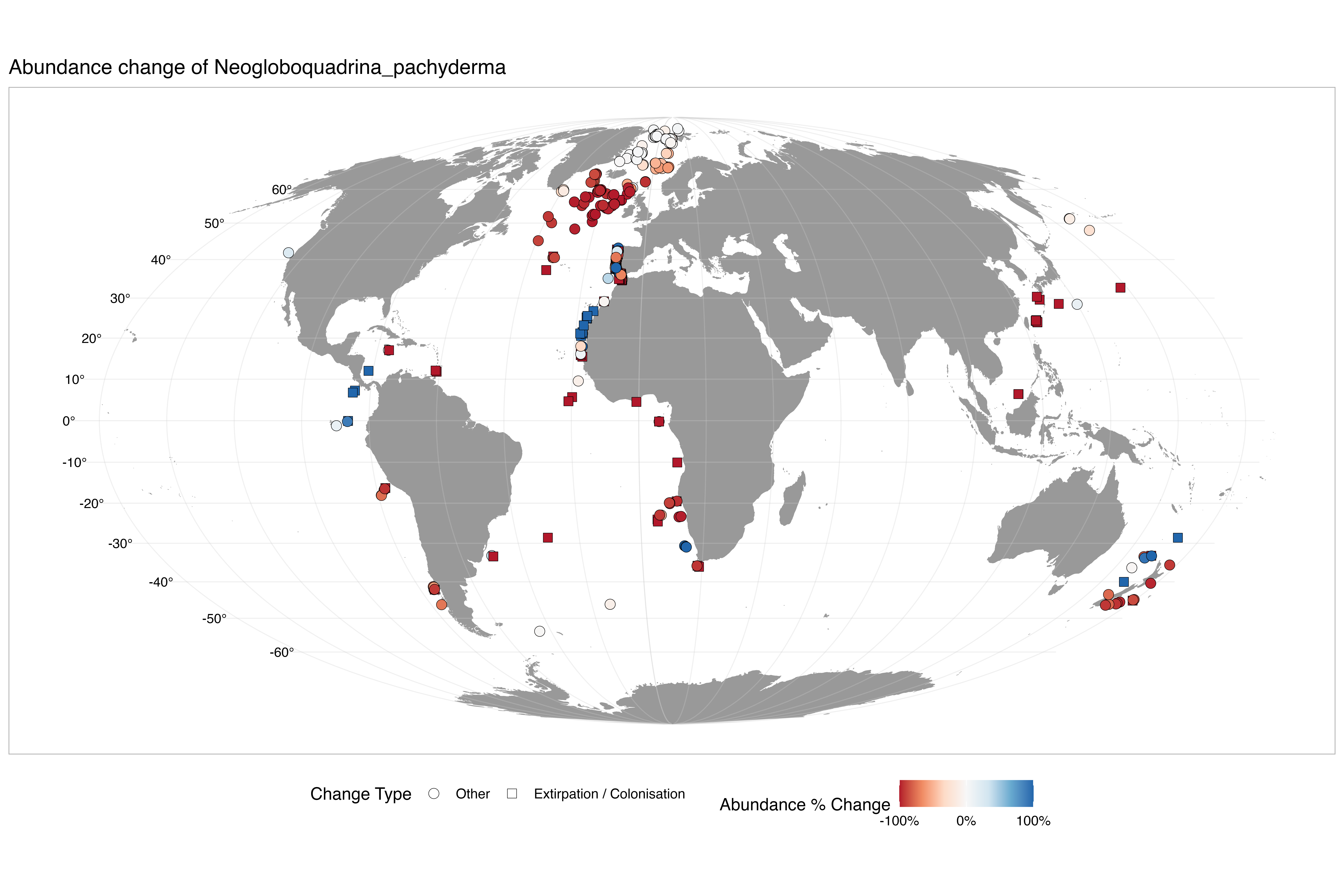

### abundance_change_Orbulina_universa.png

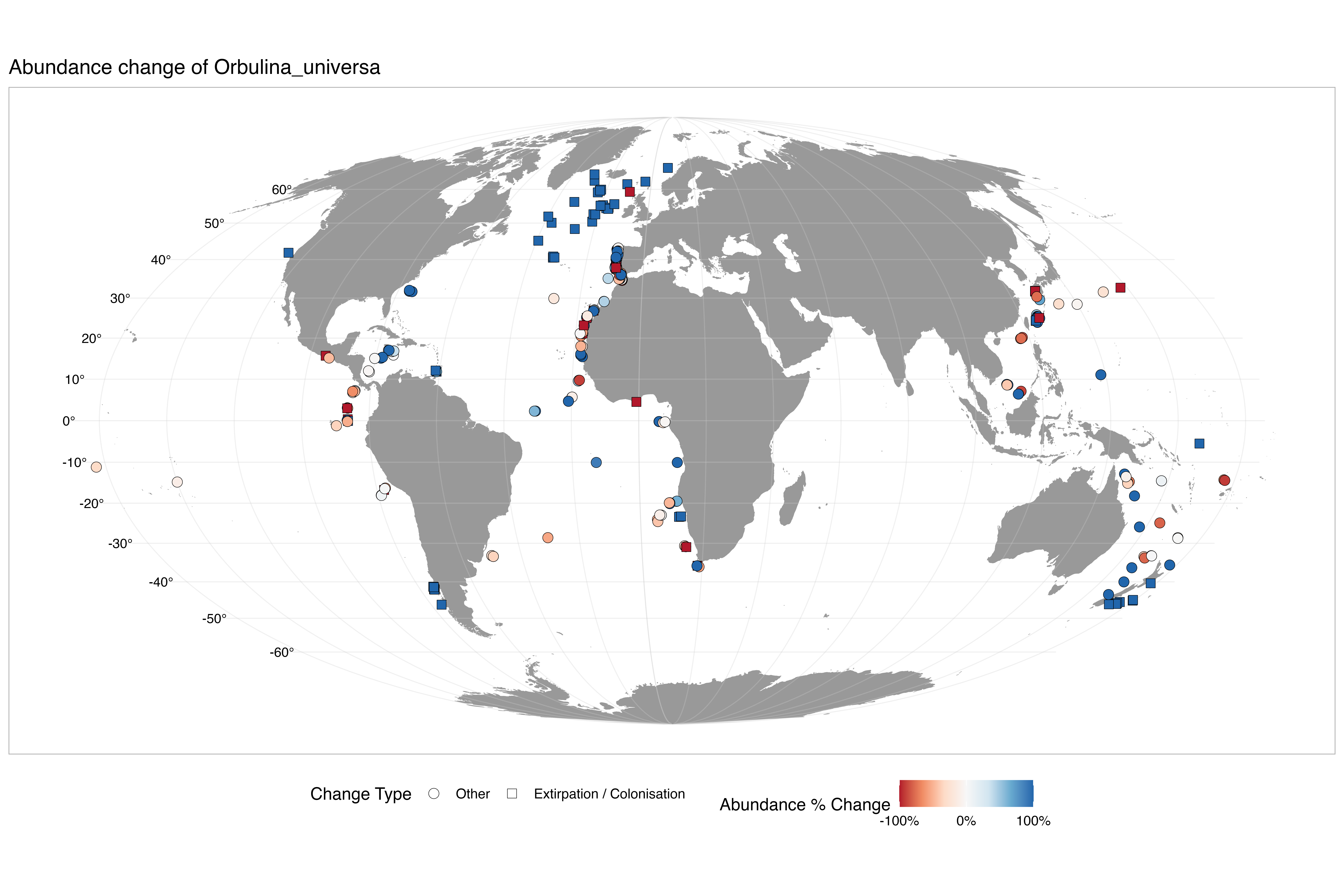

### abundance_change_Pulleniatina_obliquiloculata.png

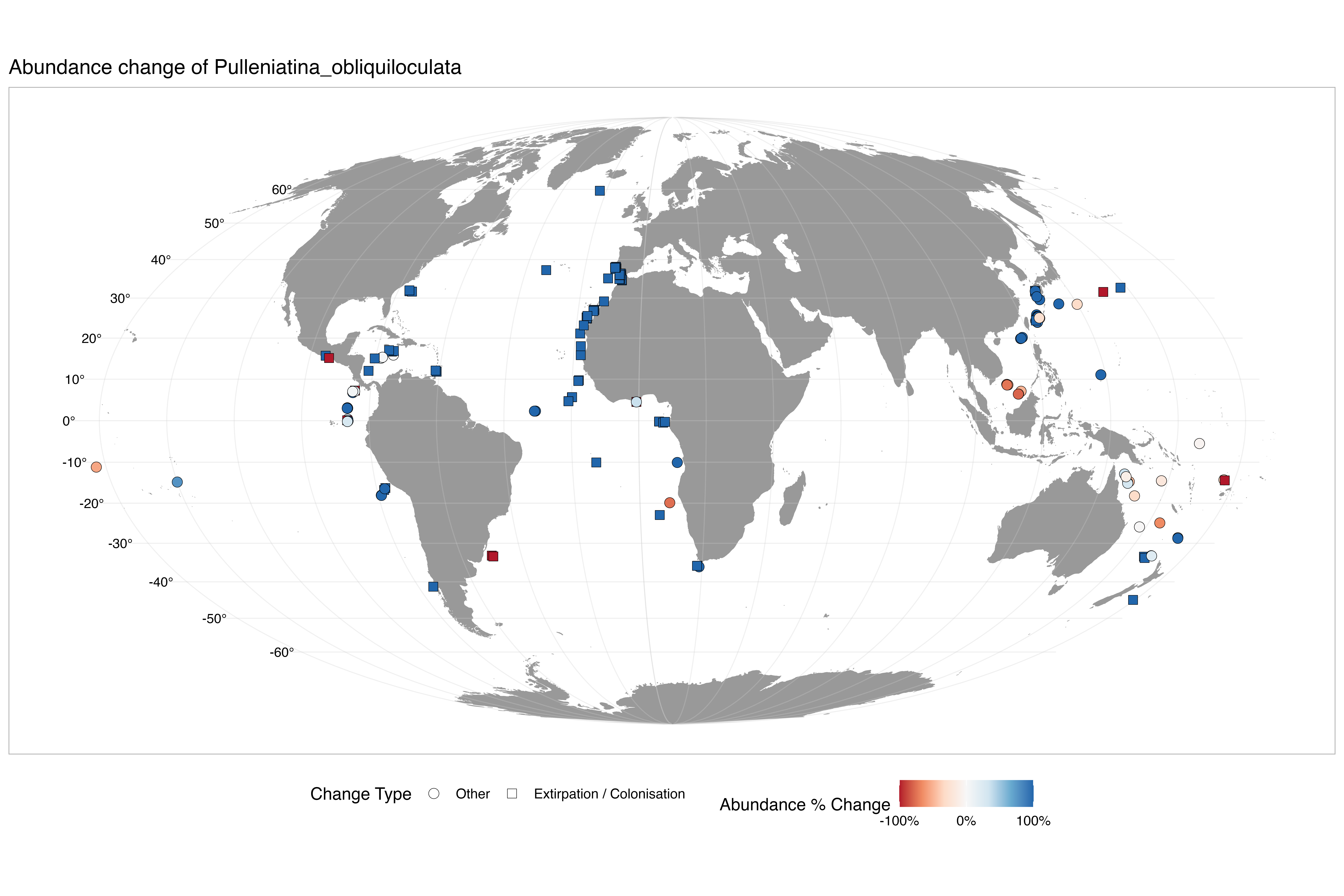

### abundance_change_Sphaeroidinella_dehiscens.png

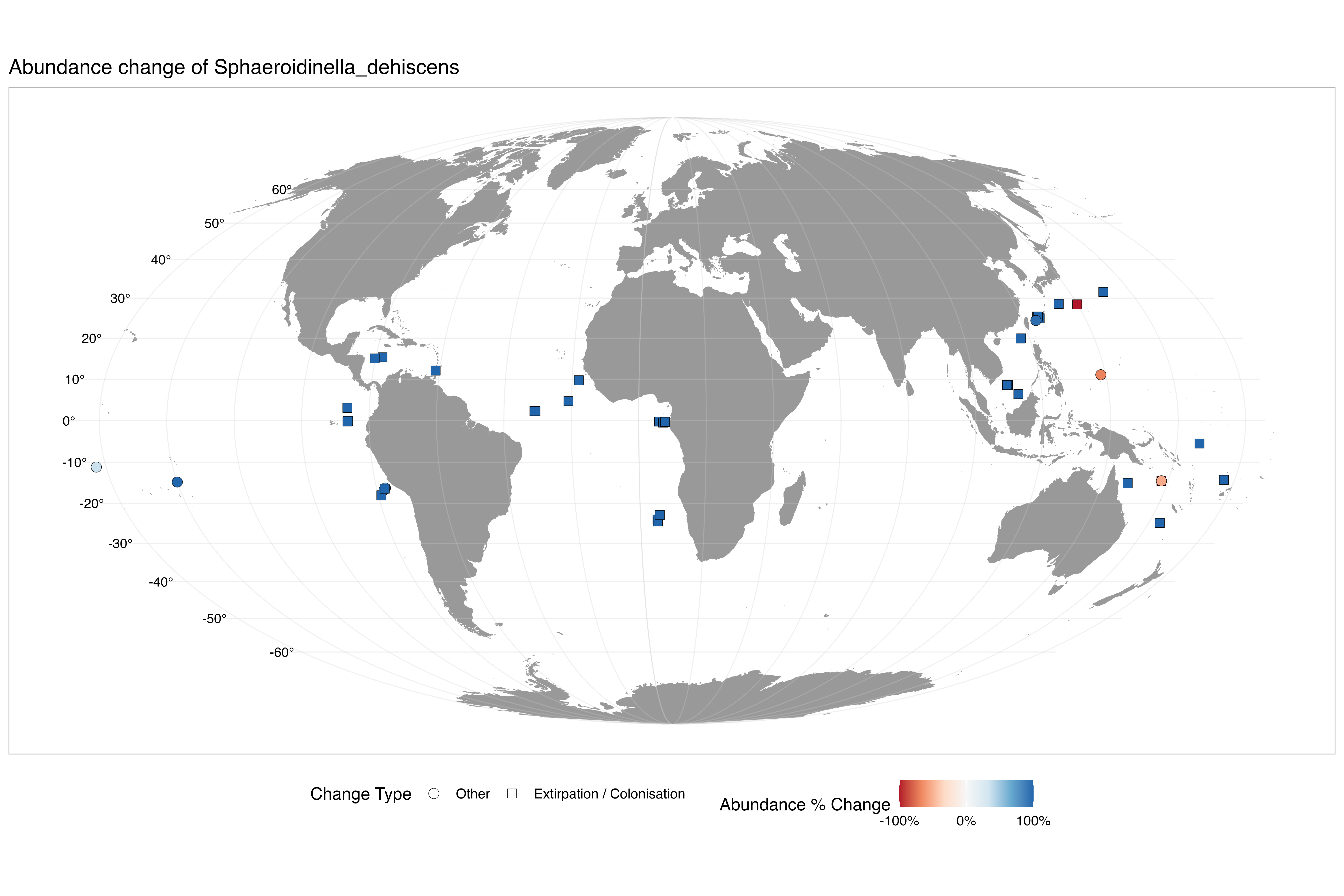

### abundance_change_Tenuitella_iota.png

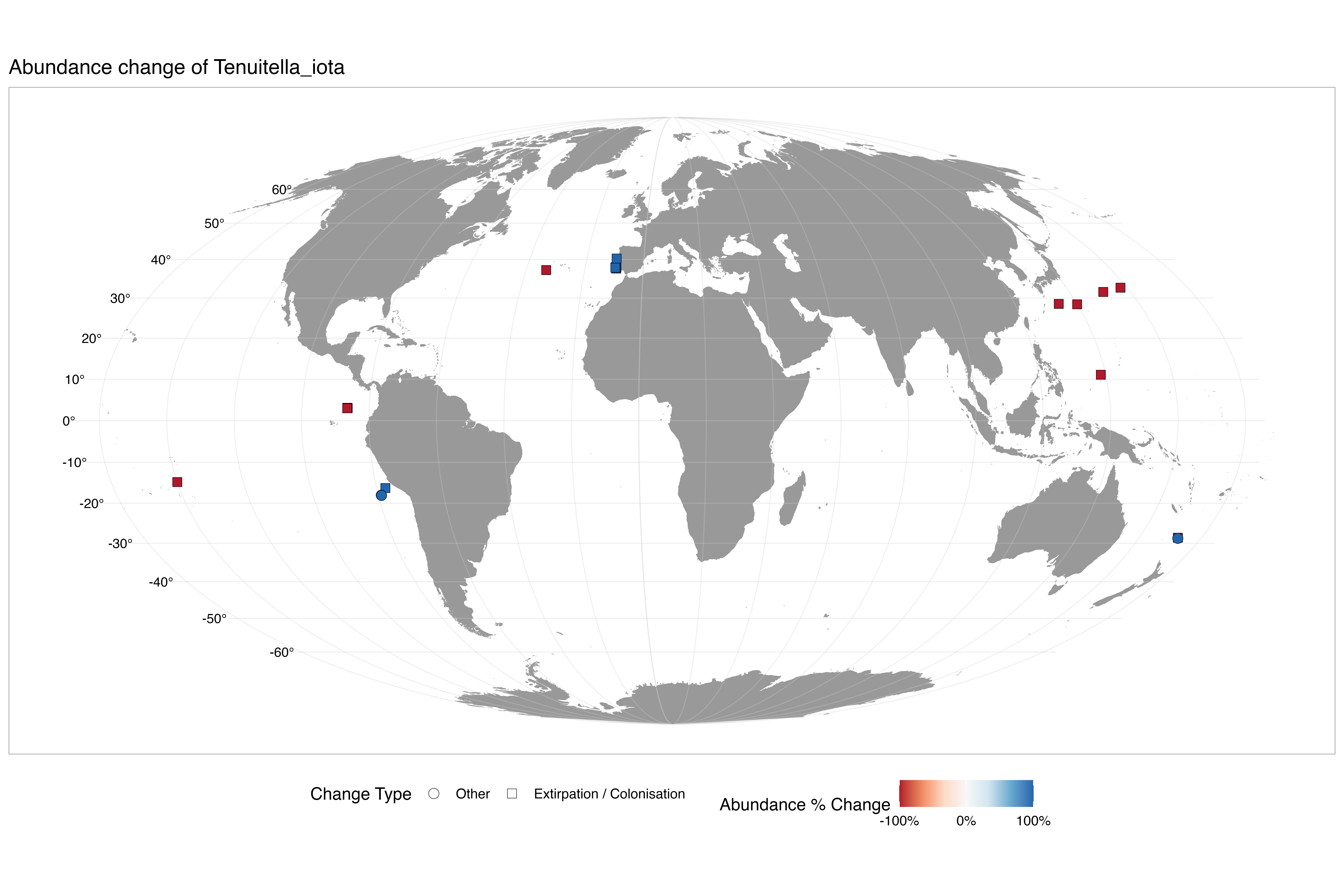

### abundance_change_Trilobatus_sacculifer.png

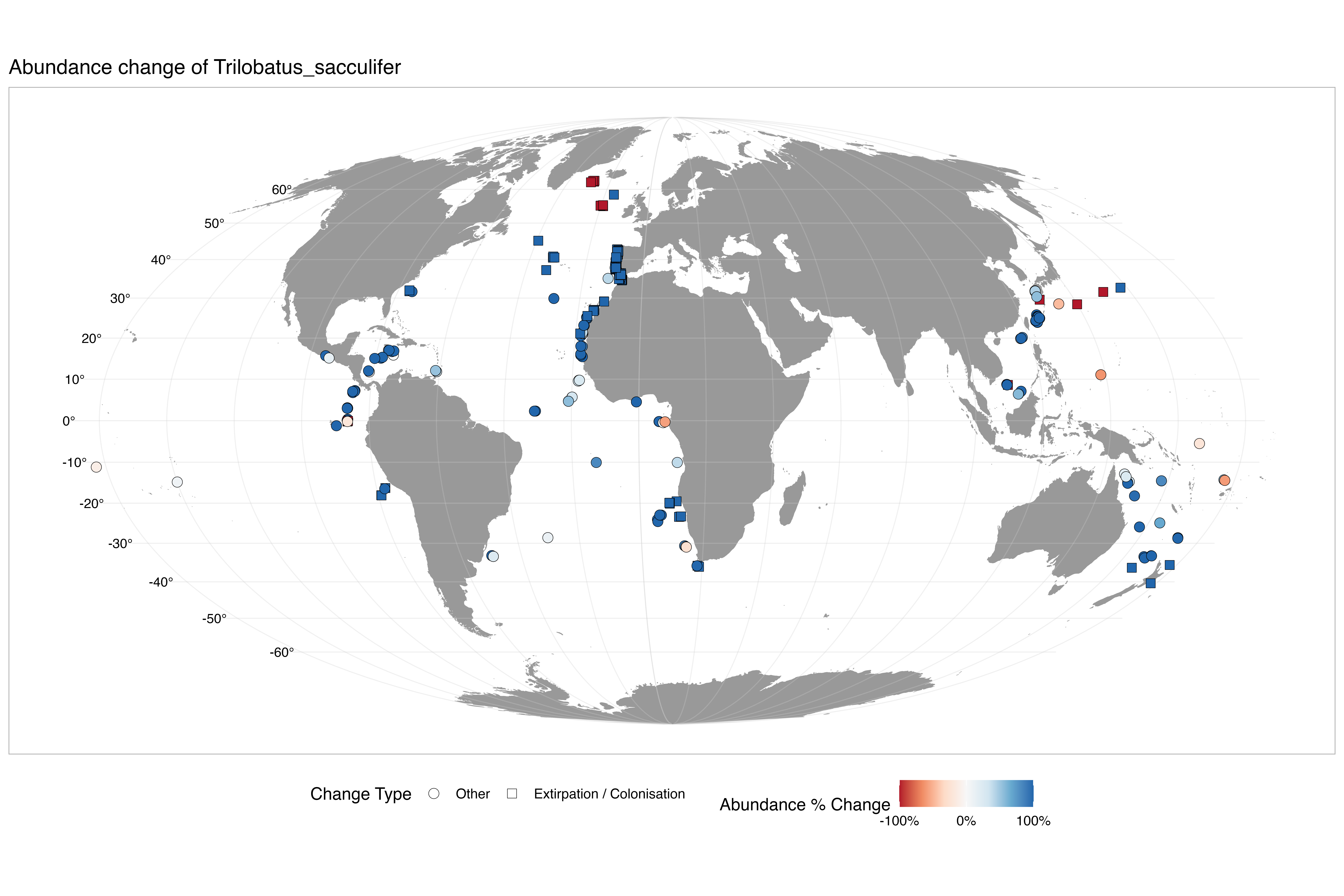

### abundance_change_Turborotalita_humilis.png

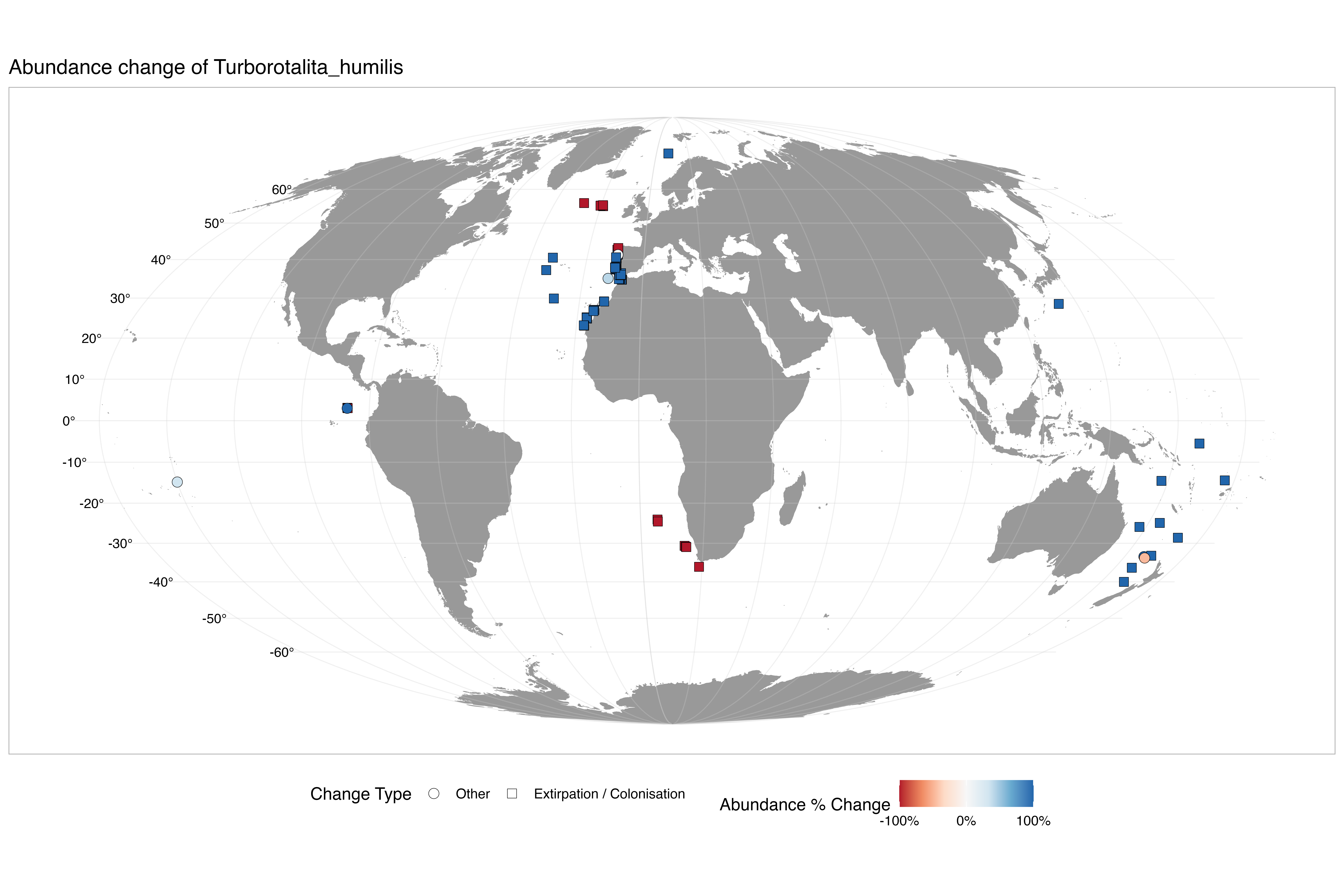

### abundance_change_Turborotalita_quinqueloba.png

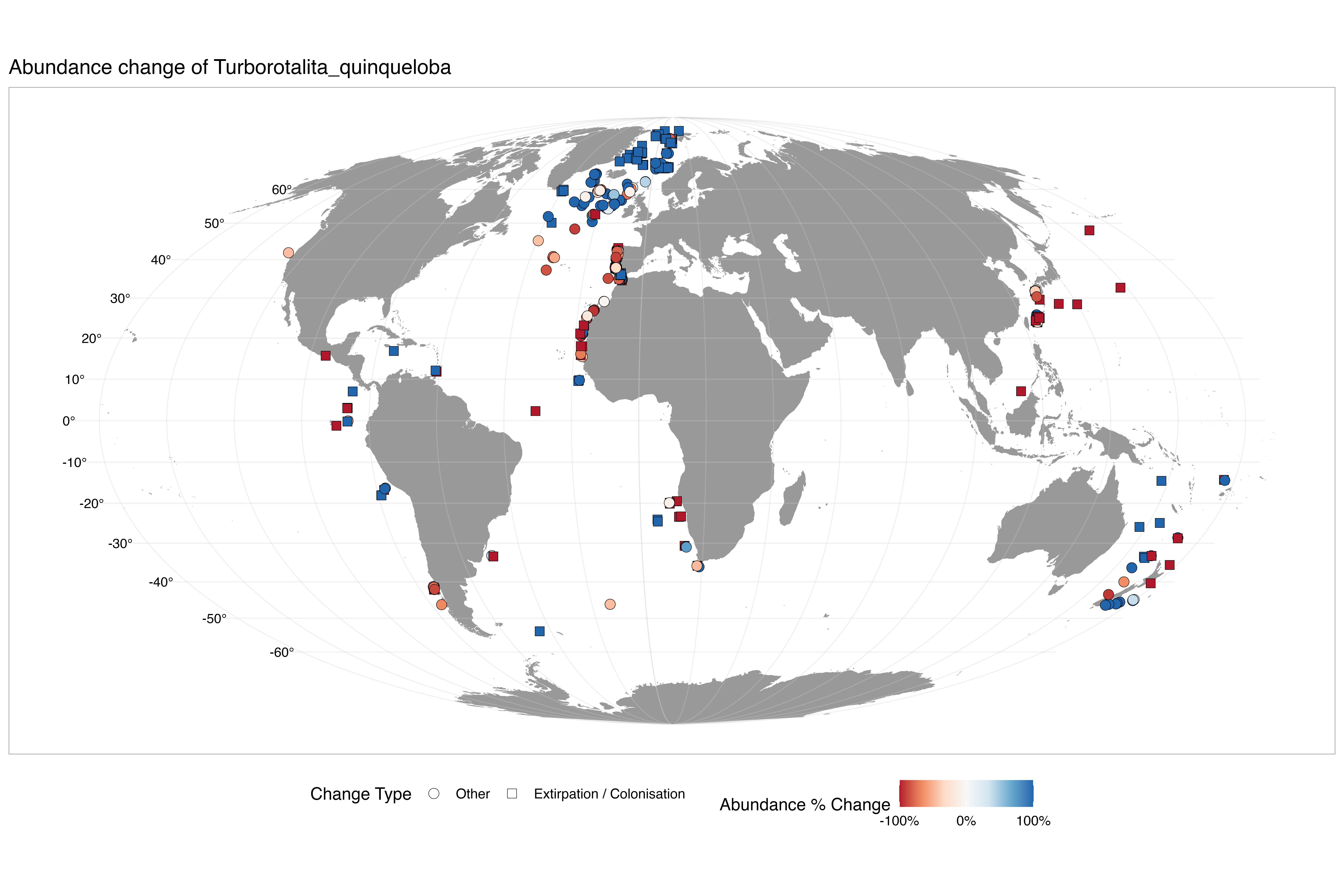

### interpolated_abundance_change_Beella_digitata.png

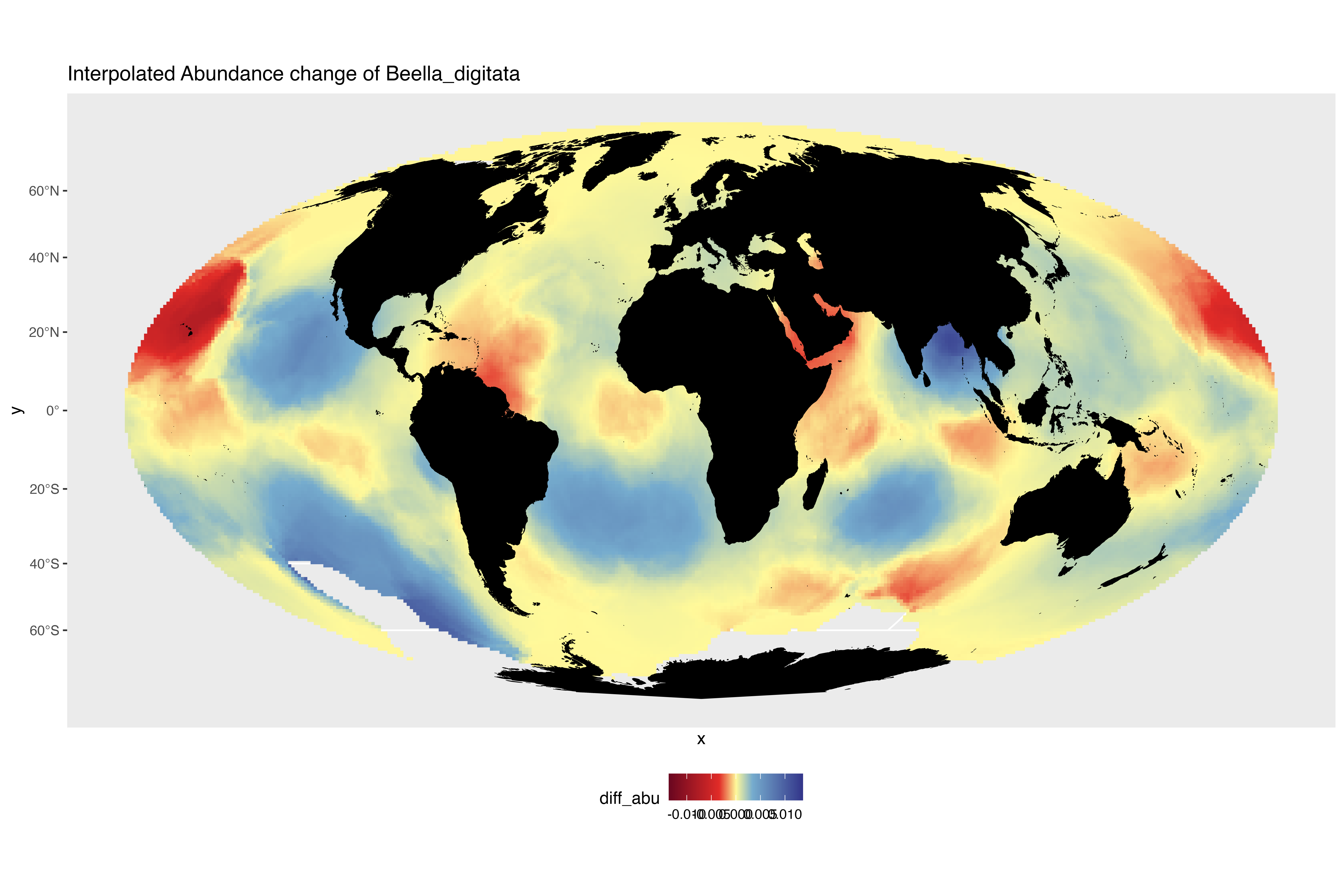

### interpolated_abundance_change_Candeina_nitida.png

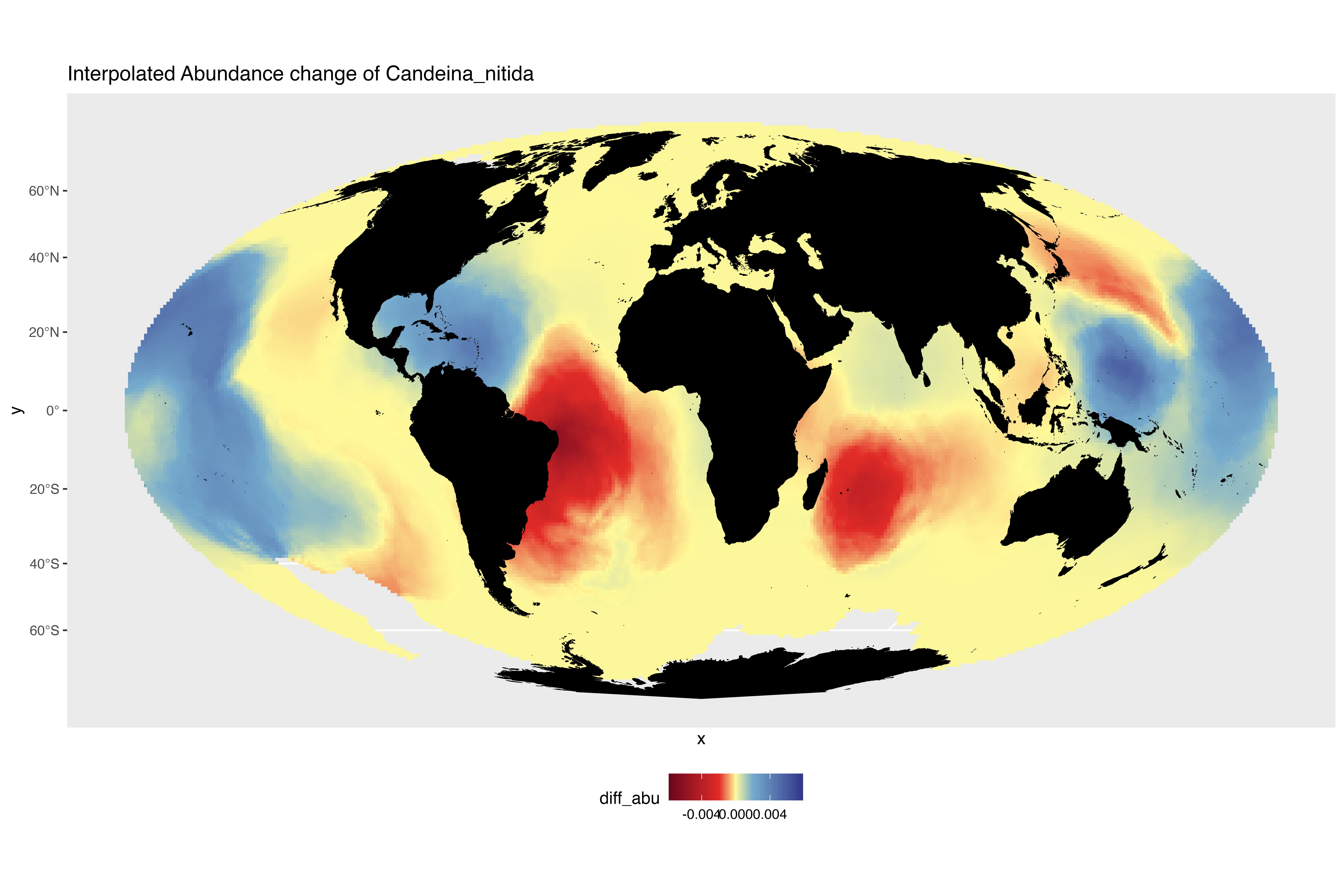

### interpolated_abundance_change_Dentigloborotalia_anfracta.png

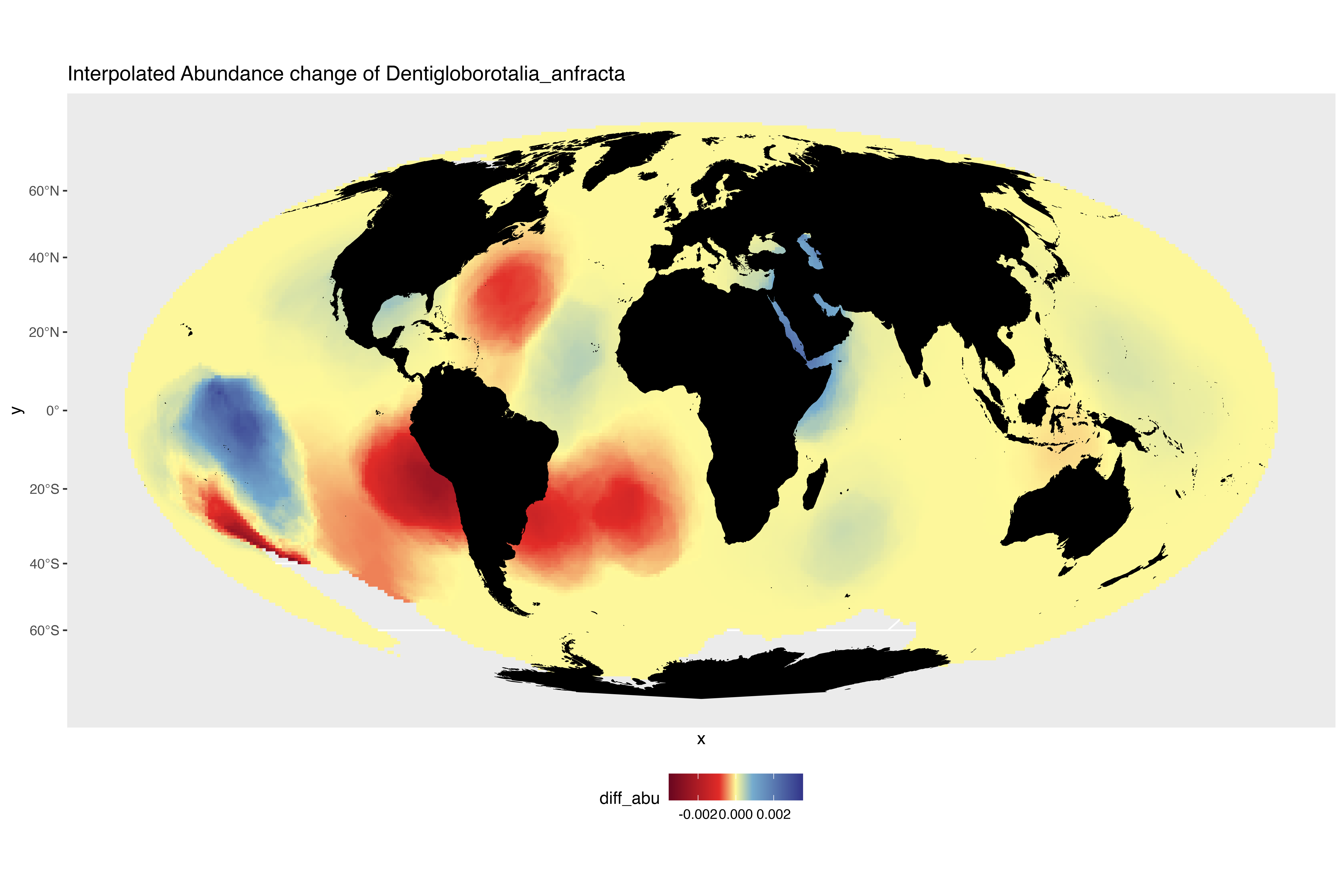

### interpolated_abundance_change_Globigerina_bulloides.png

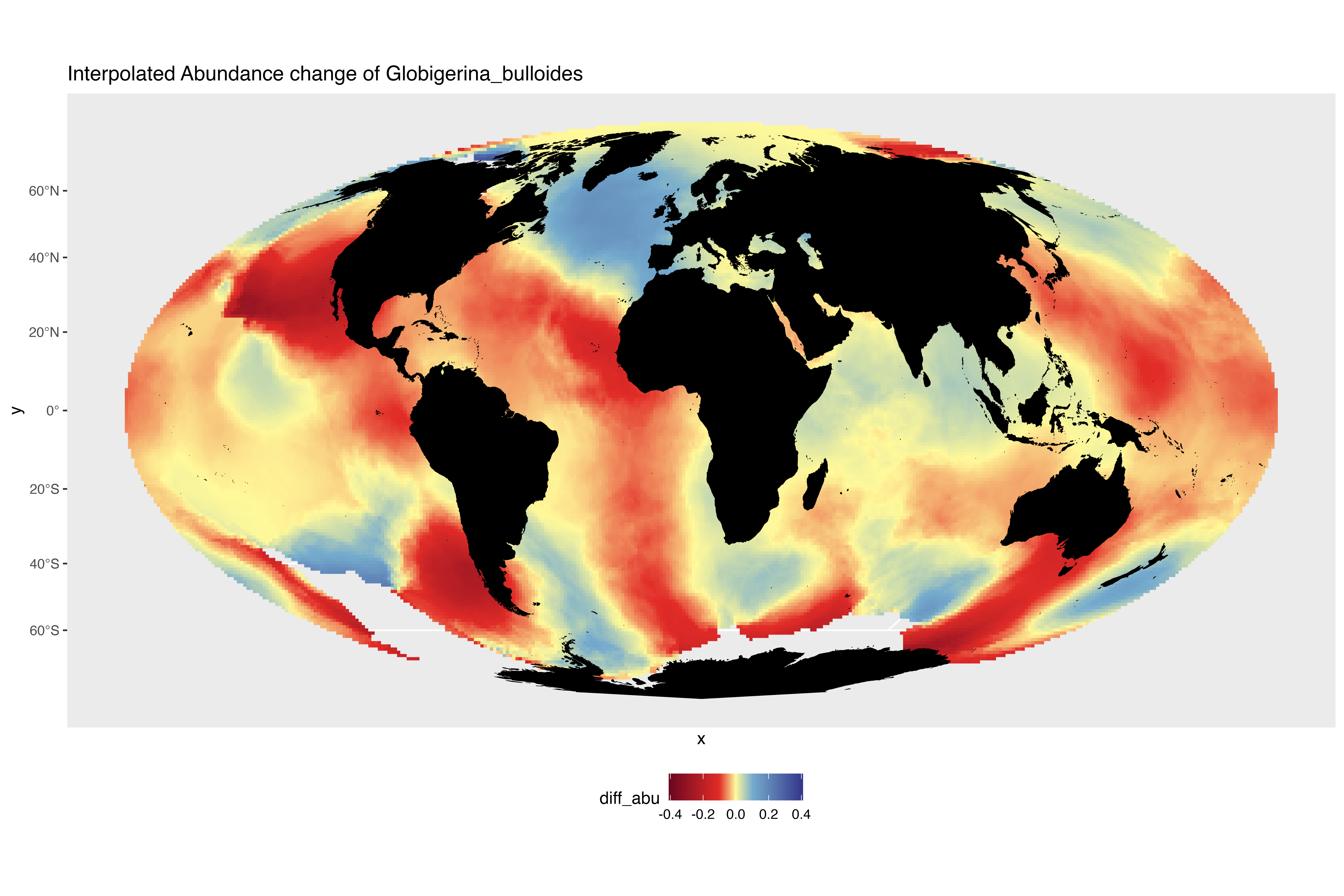

### interpolated_abundance_change_Globigerina_falconensis.png

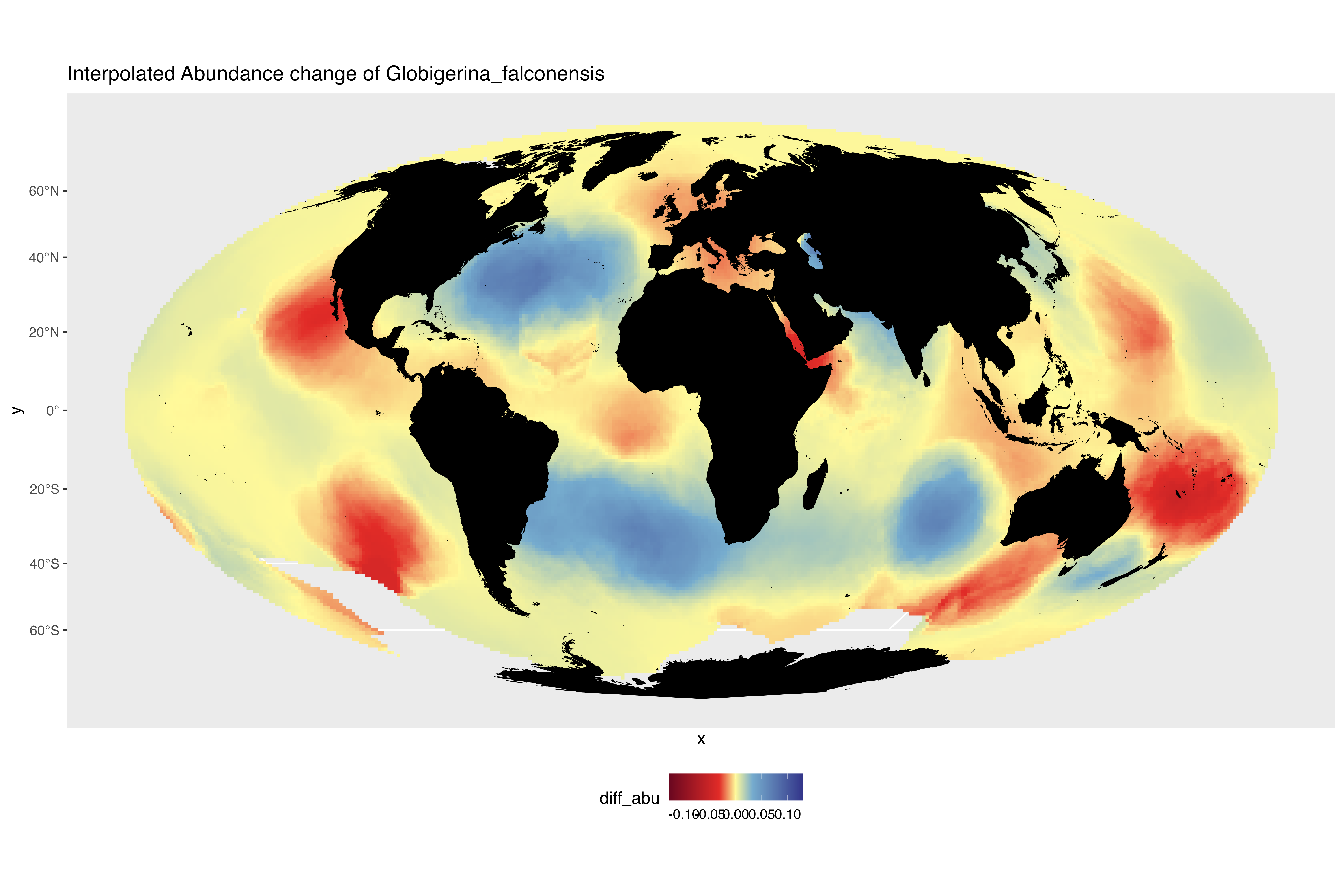

### interpolated_abundance_change_Globigerinella_adamsi.png

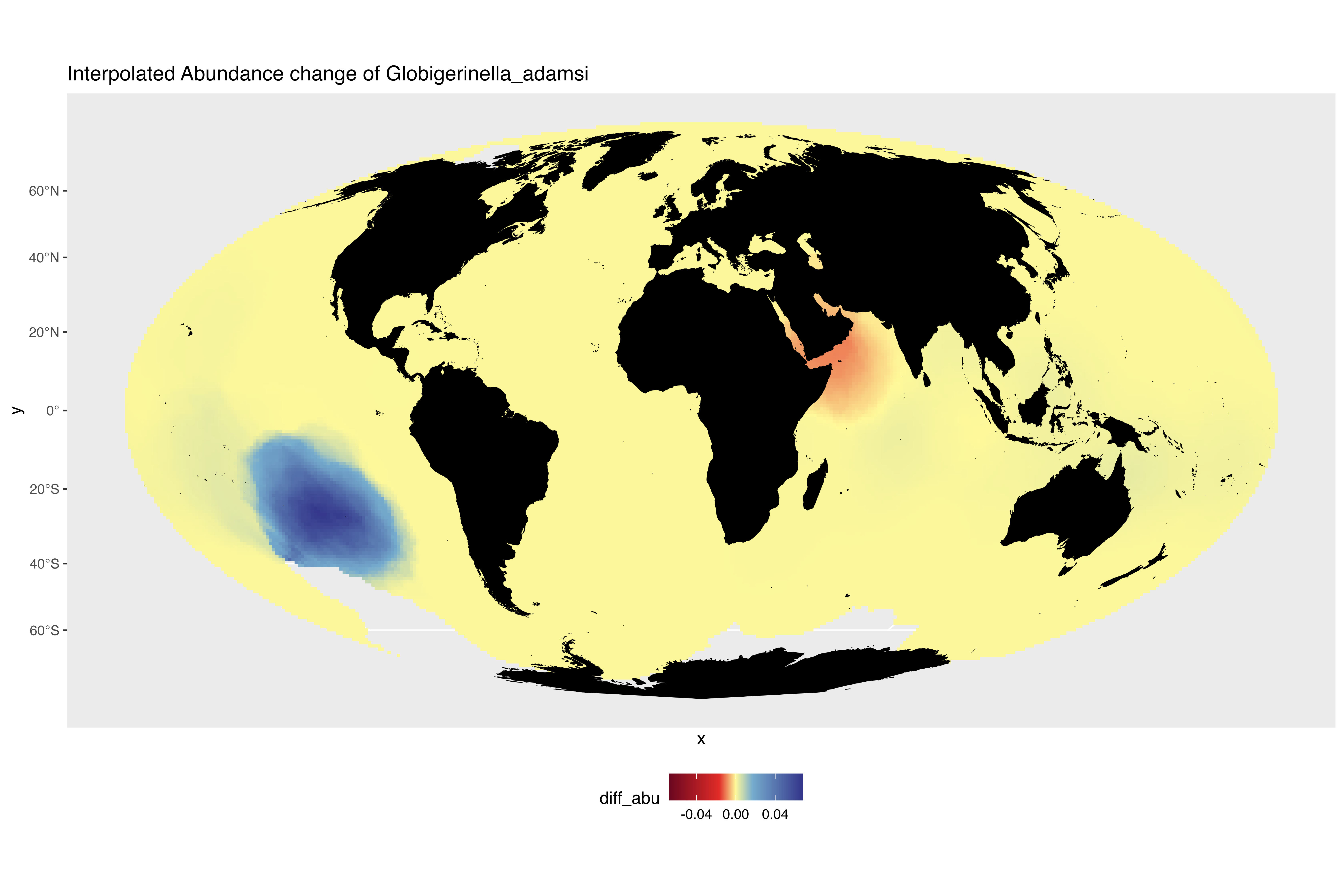

### interpolated_abundance_change_Globigerinella_calida.png

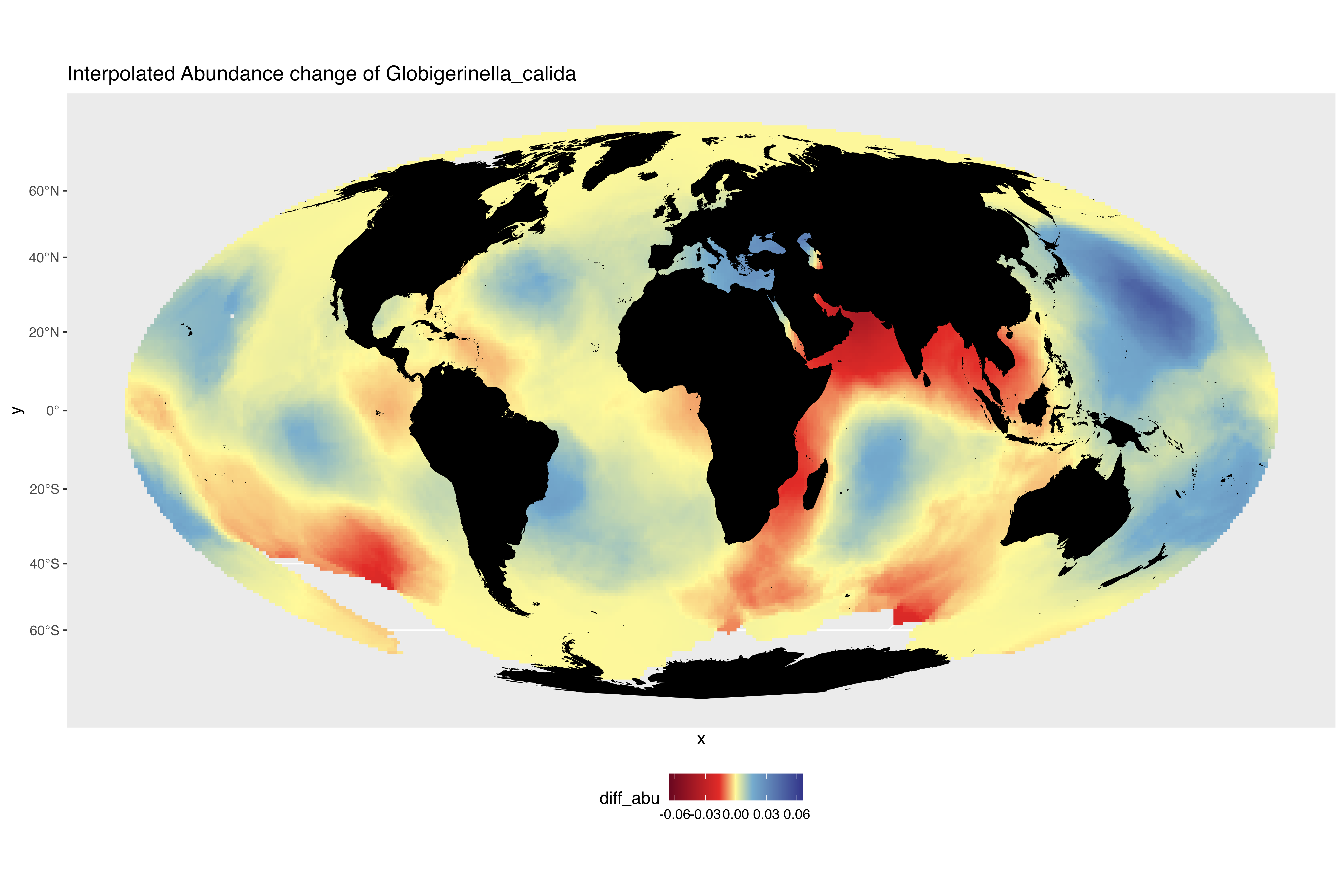

### interpolated_abundance_change_Globigerinella_siphonifera.png

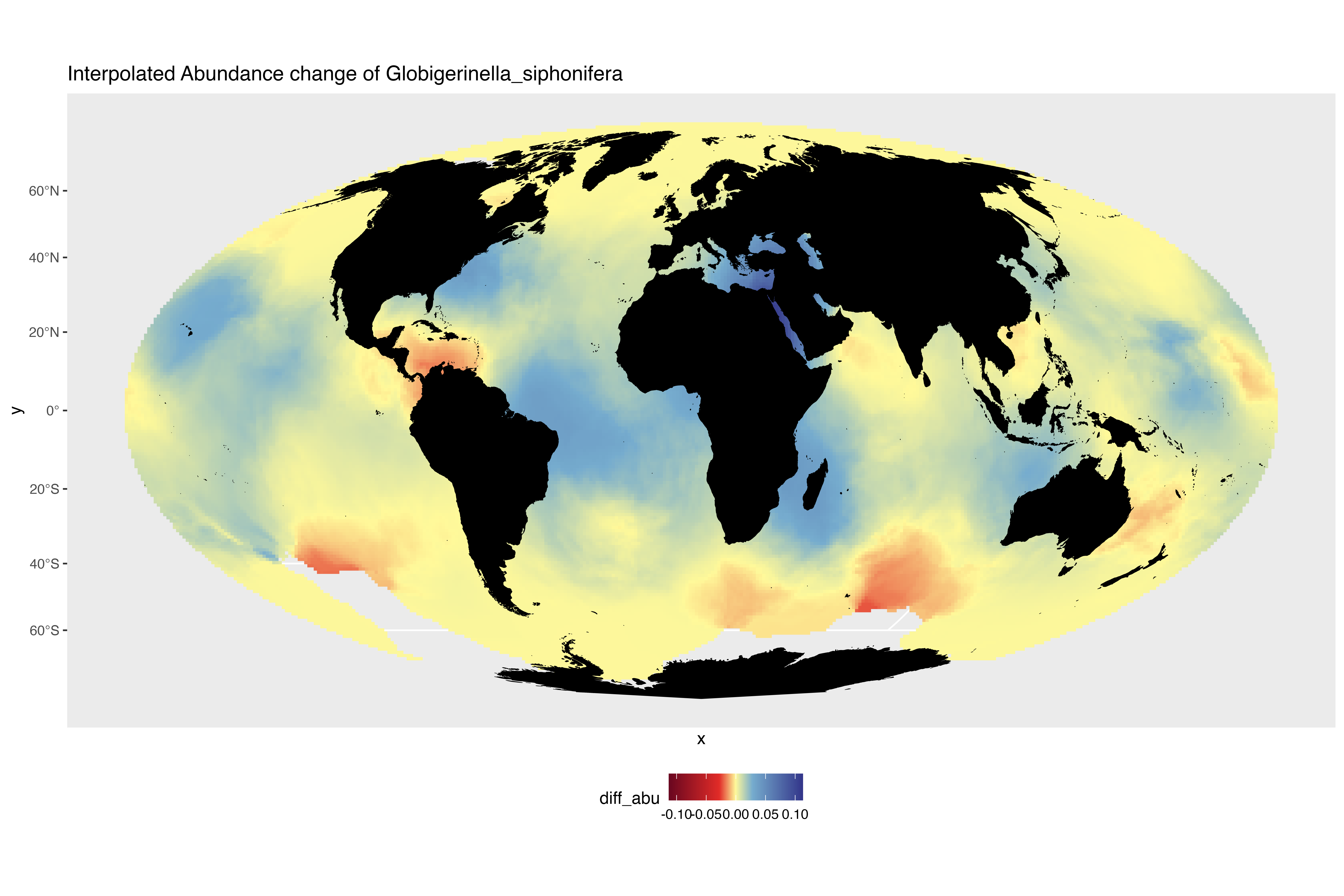

### interpolated_abundance_change_Globigerinita_glutinata.png

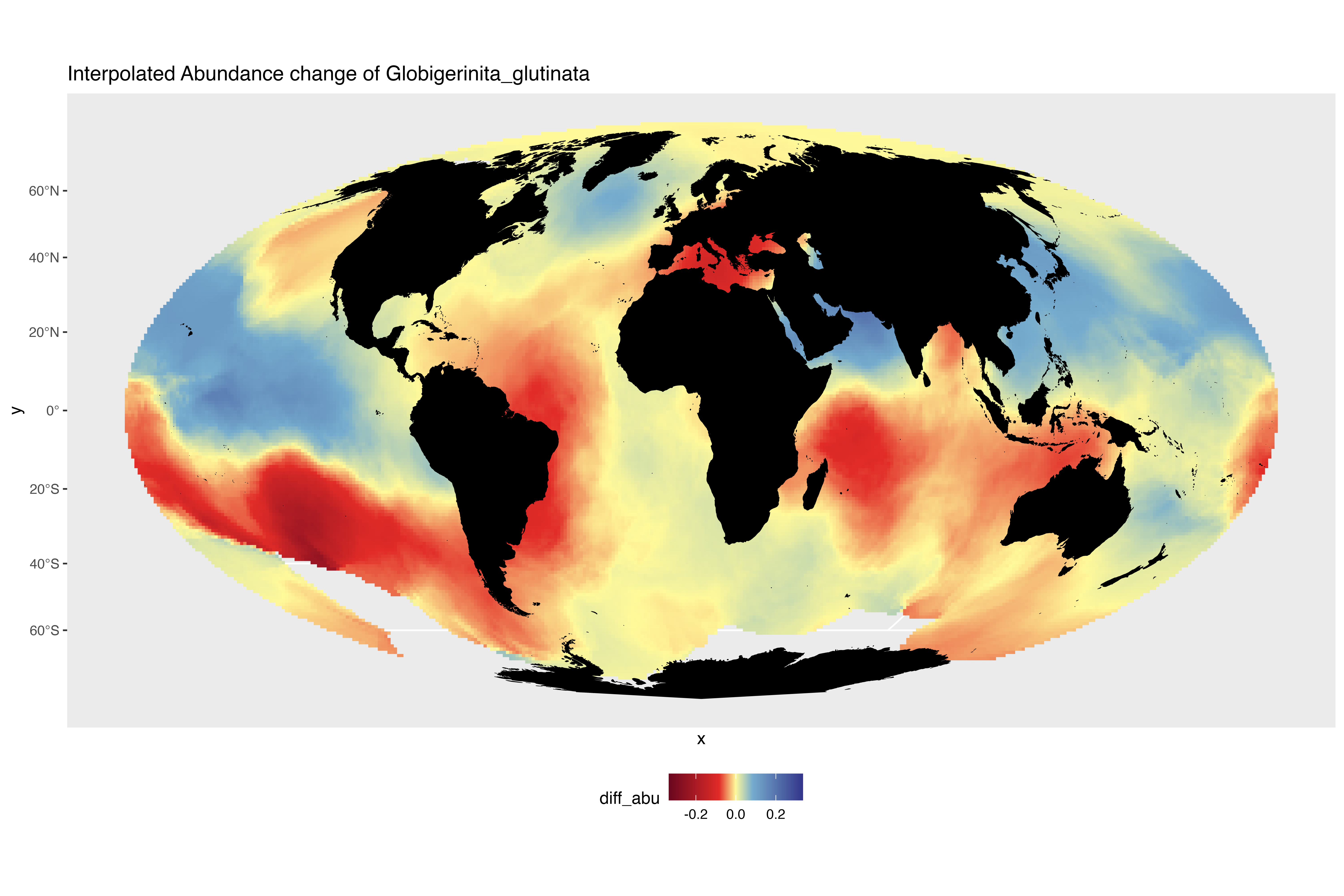

### interpolated_abundance_change_Globigerinita_uvula.png

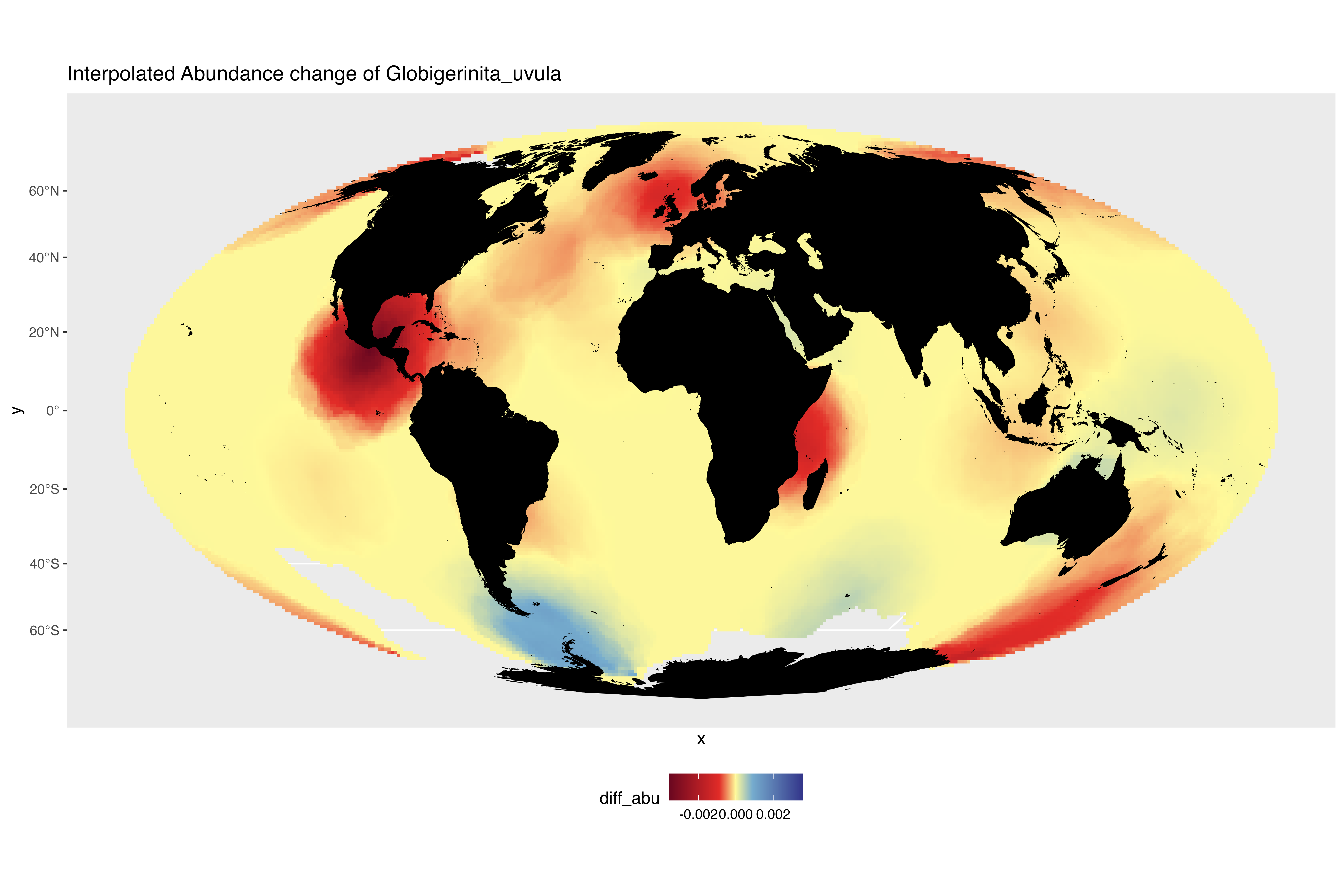

### interpolated_abundance_change_Globigerinoides_conglobatus.png

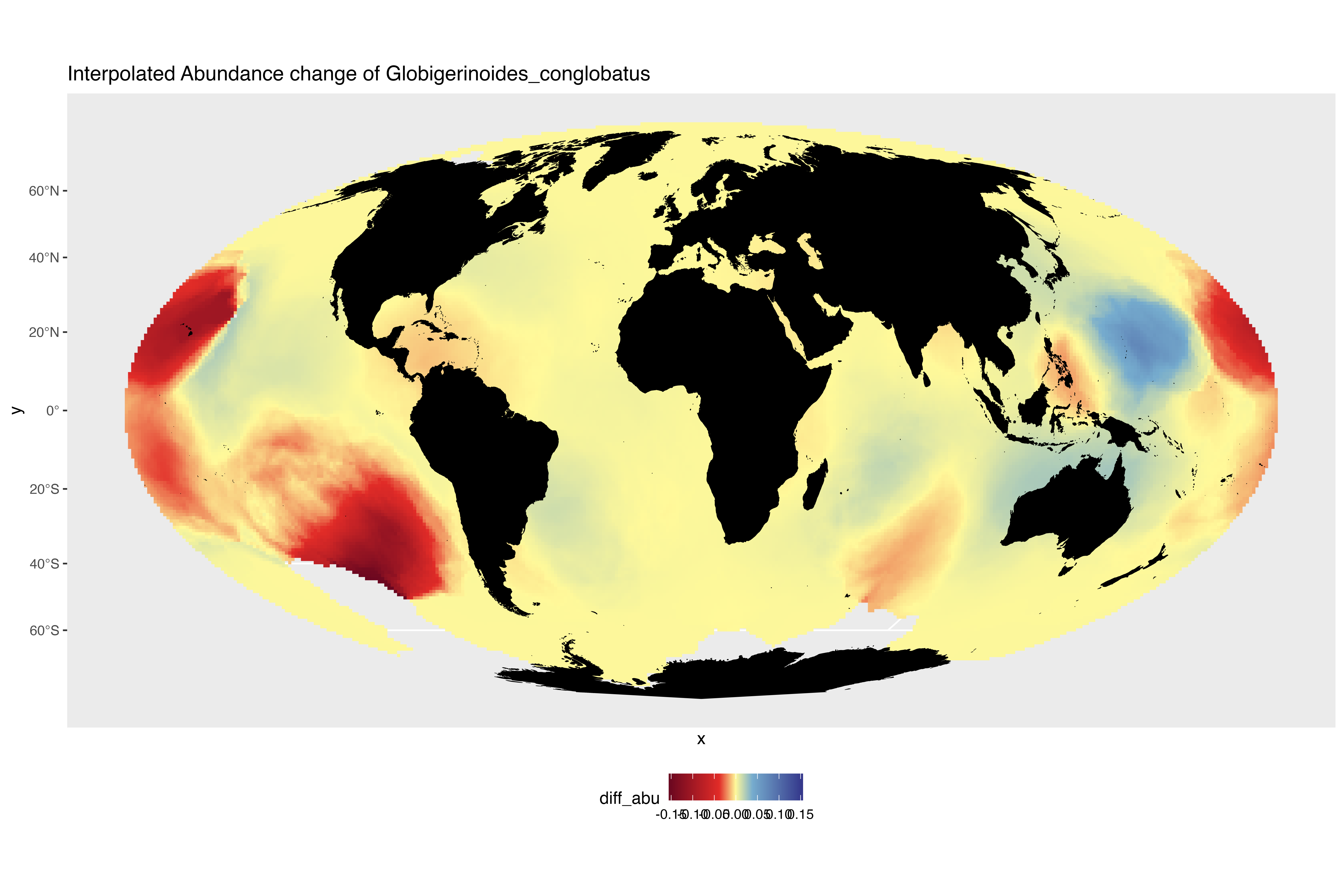

### interpolated_abundance_change_Globigerinoides_ruber.png

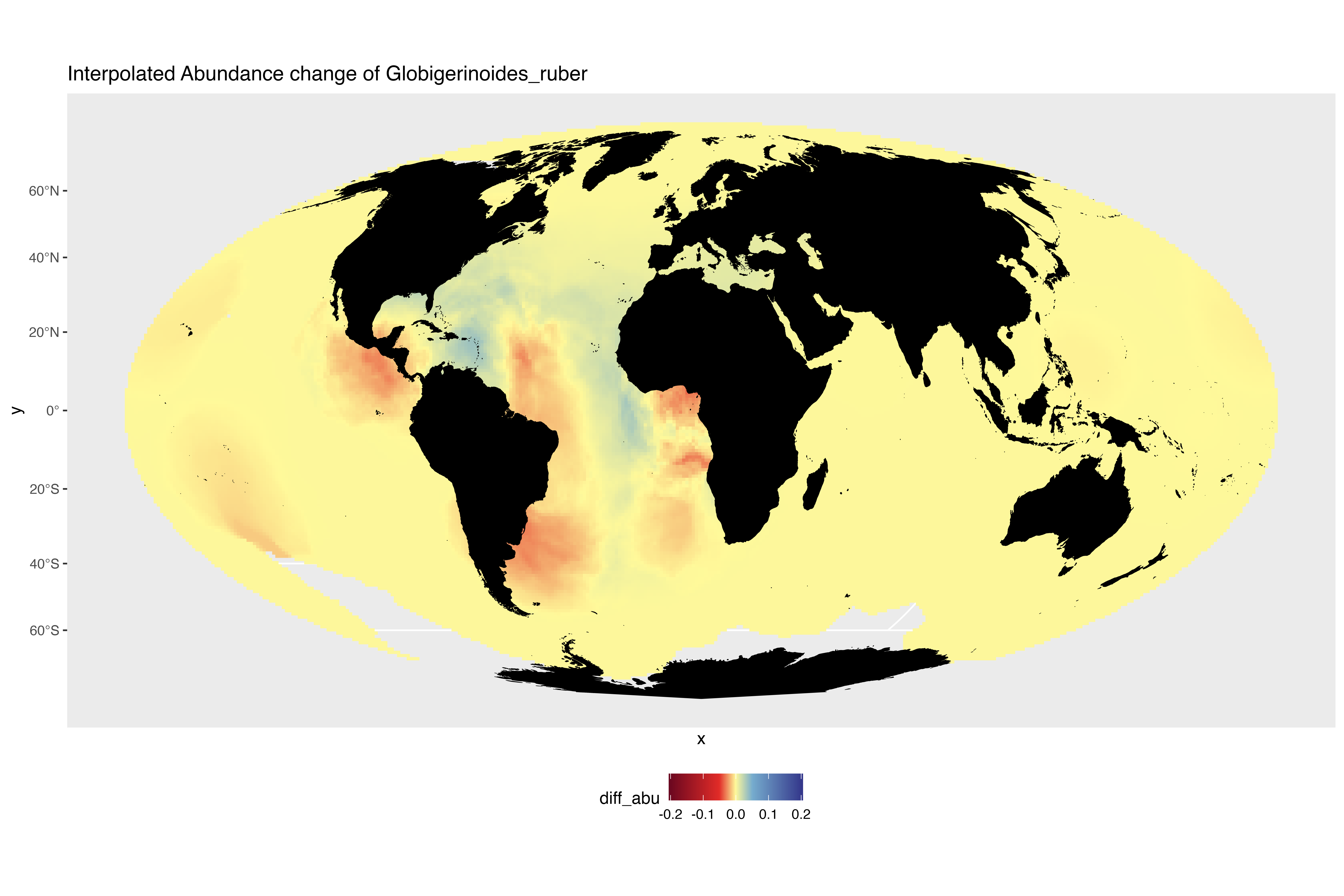

### interpolated_abundance_change_Globigerinoides_tenellus.png

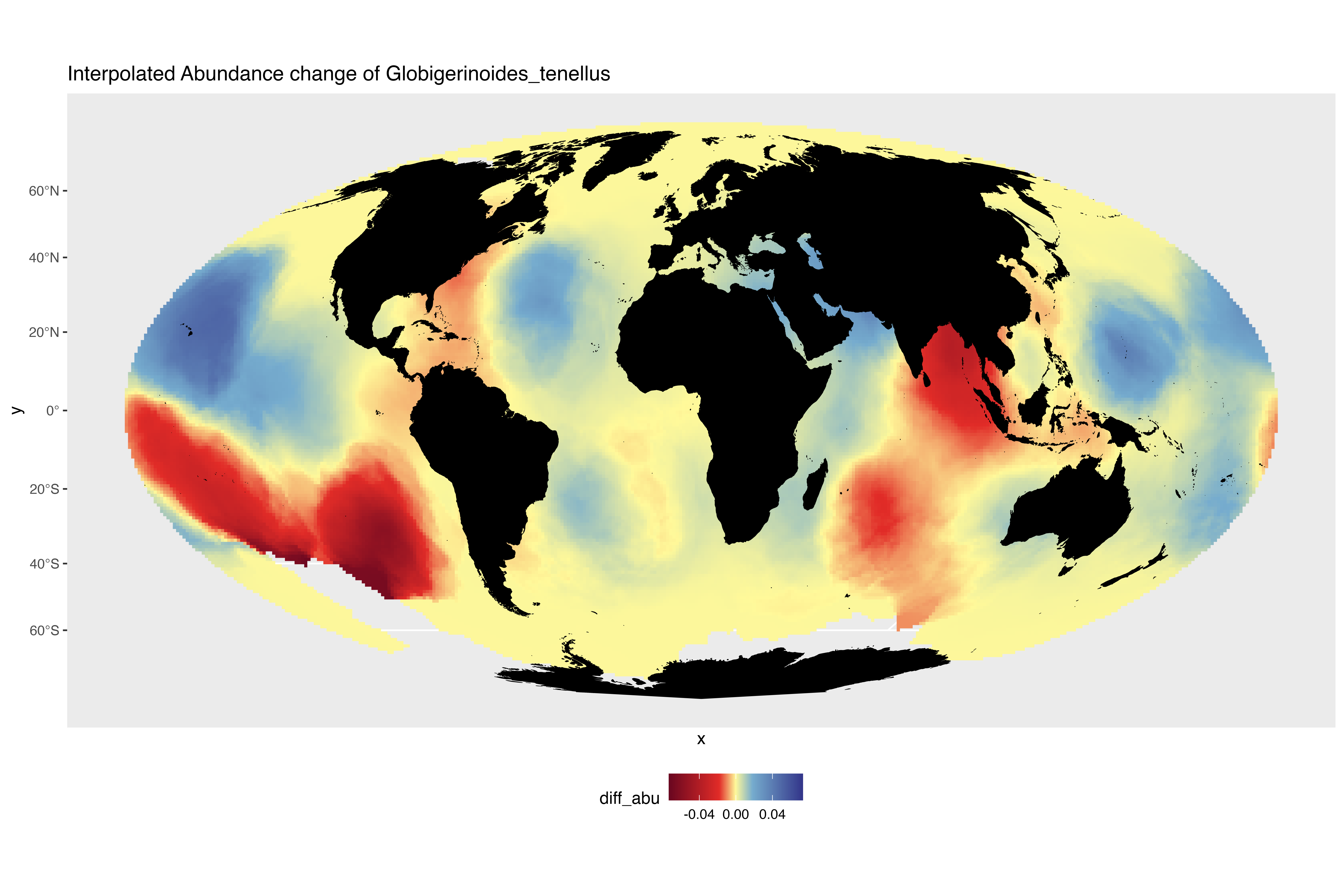

### interpolated_abundance_change_Globigerinoides_white.png

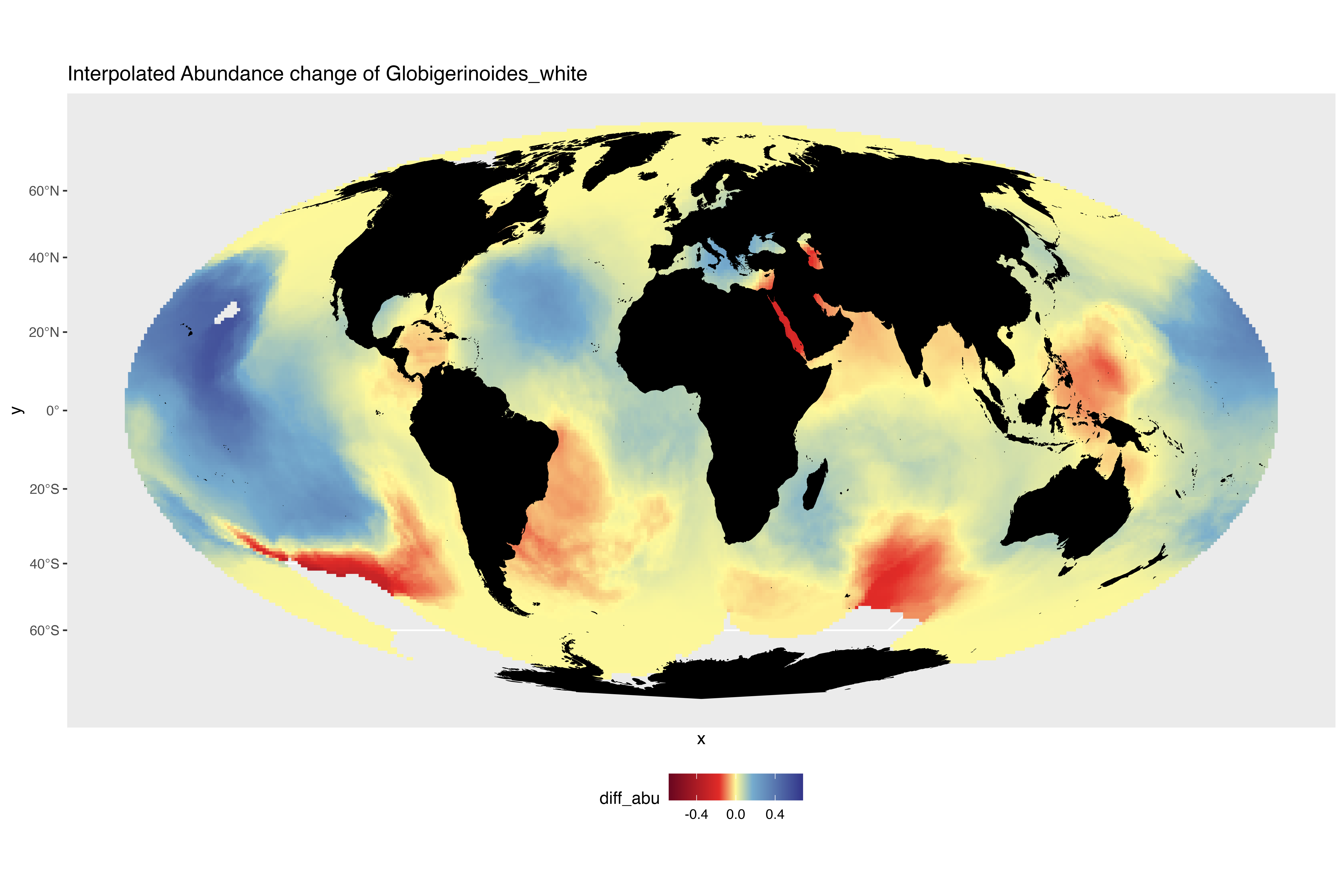

### interpolated_abundance_change_Globoconella_inflata.png

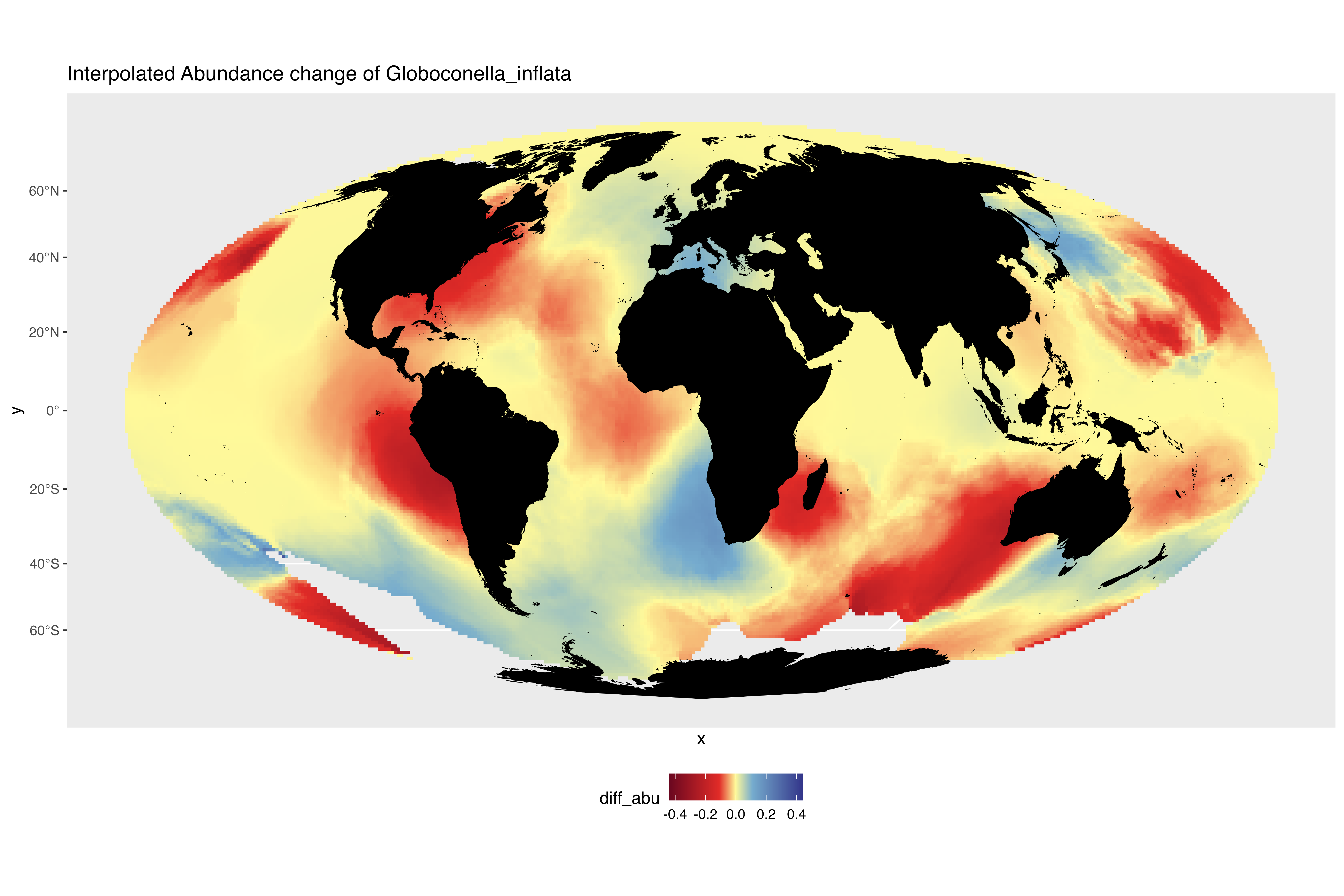

### interpolated_abundance_change_Globoquadrina_conglomerata.png

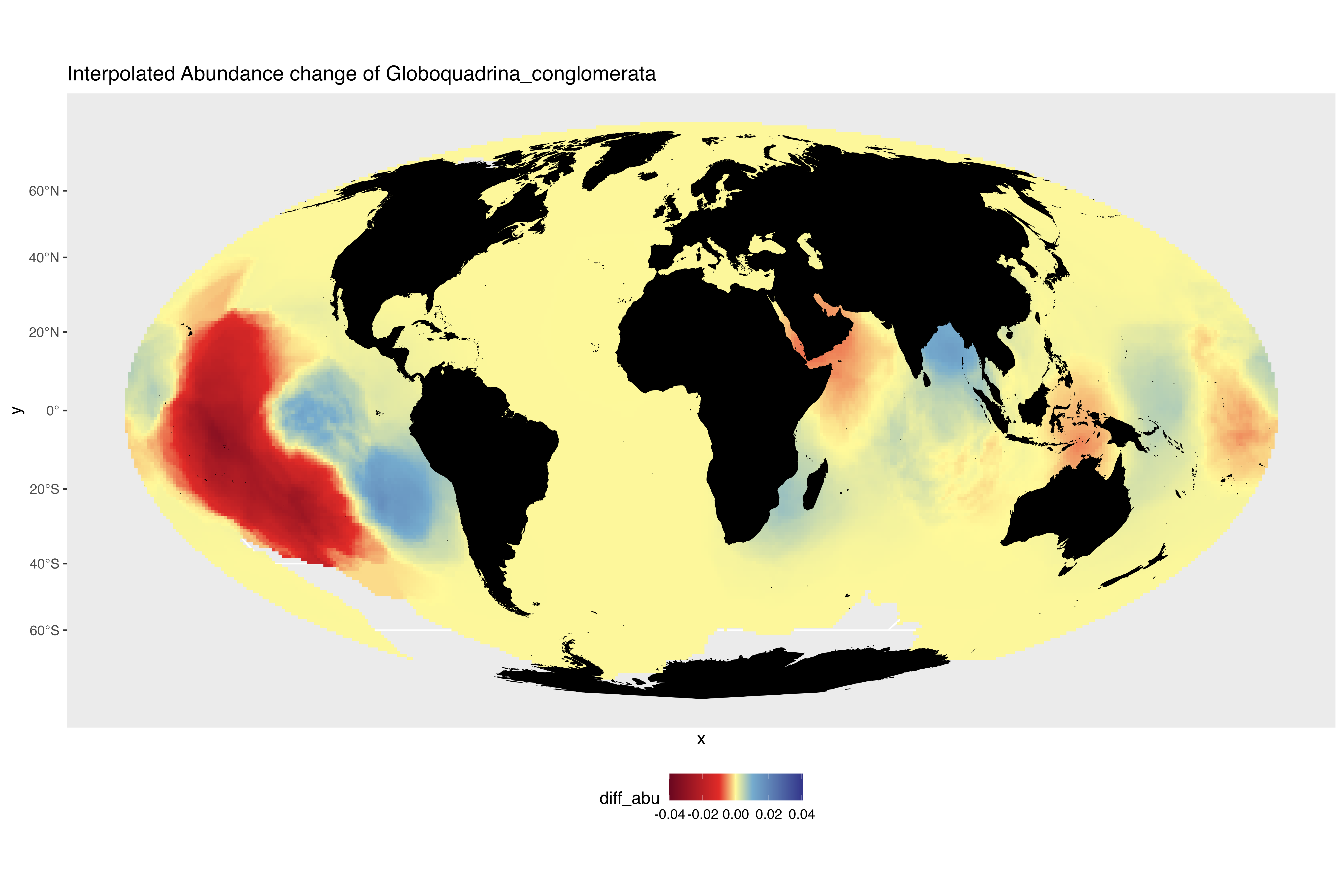

### interpolated_abundance_change_Globorotalia_crassaformis.png

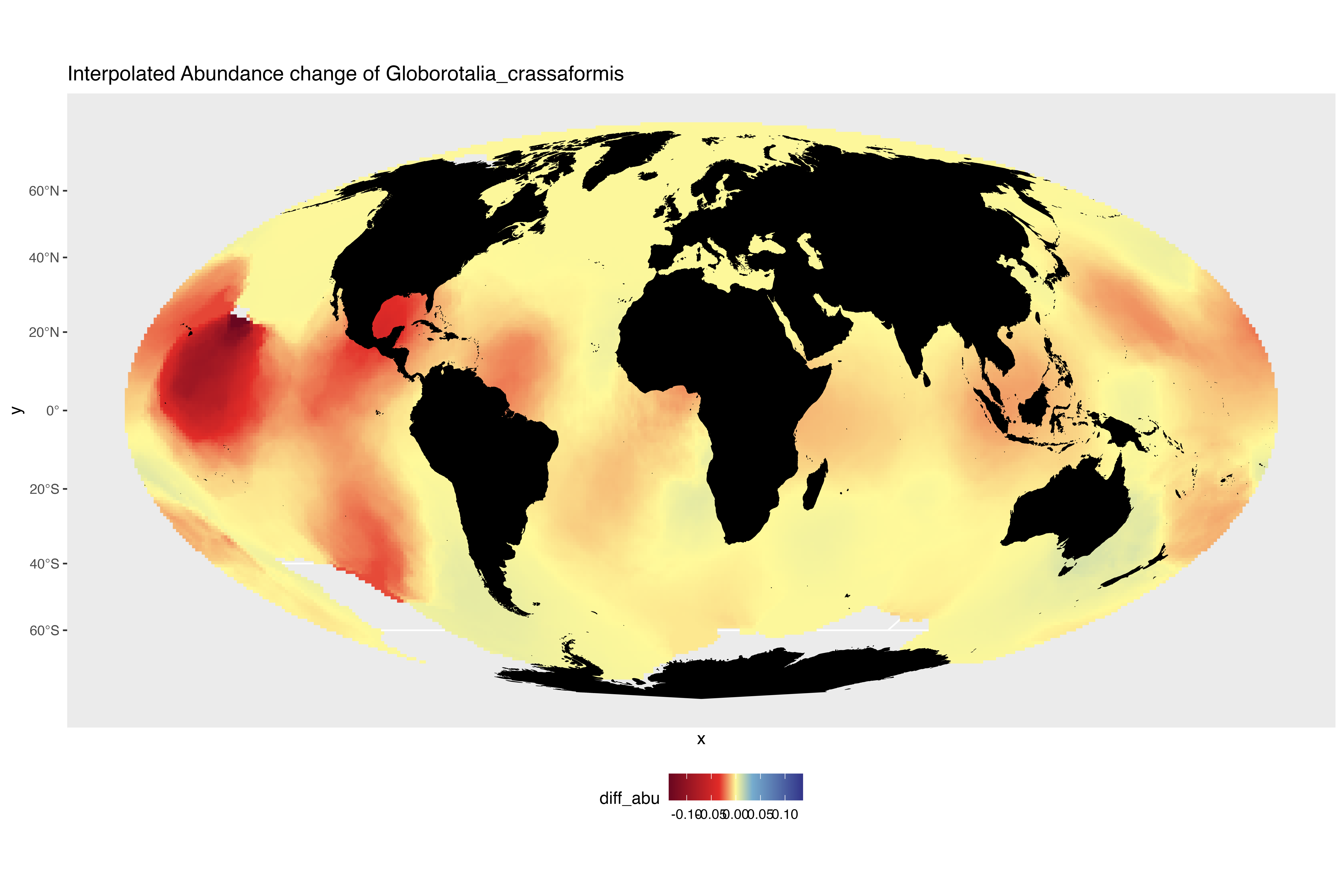

### interpolated_abundance_change_Globorotalia_hirsuta.png

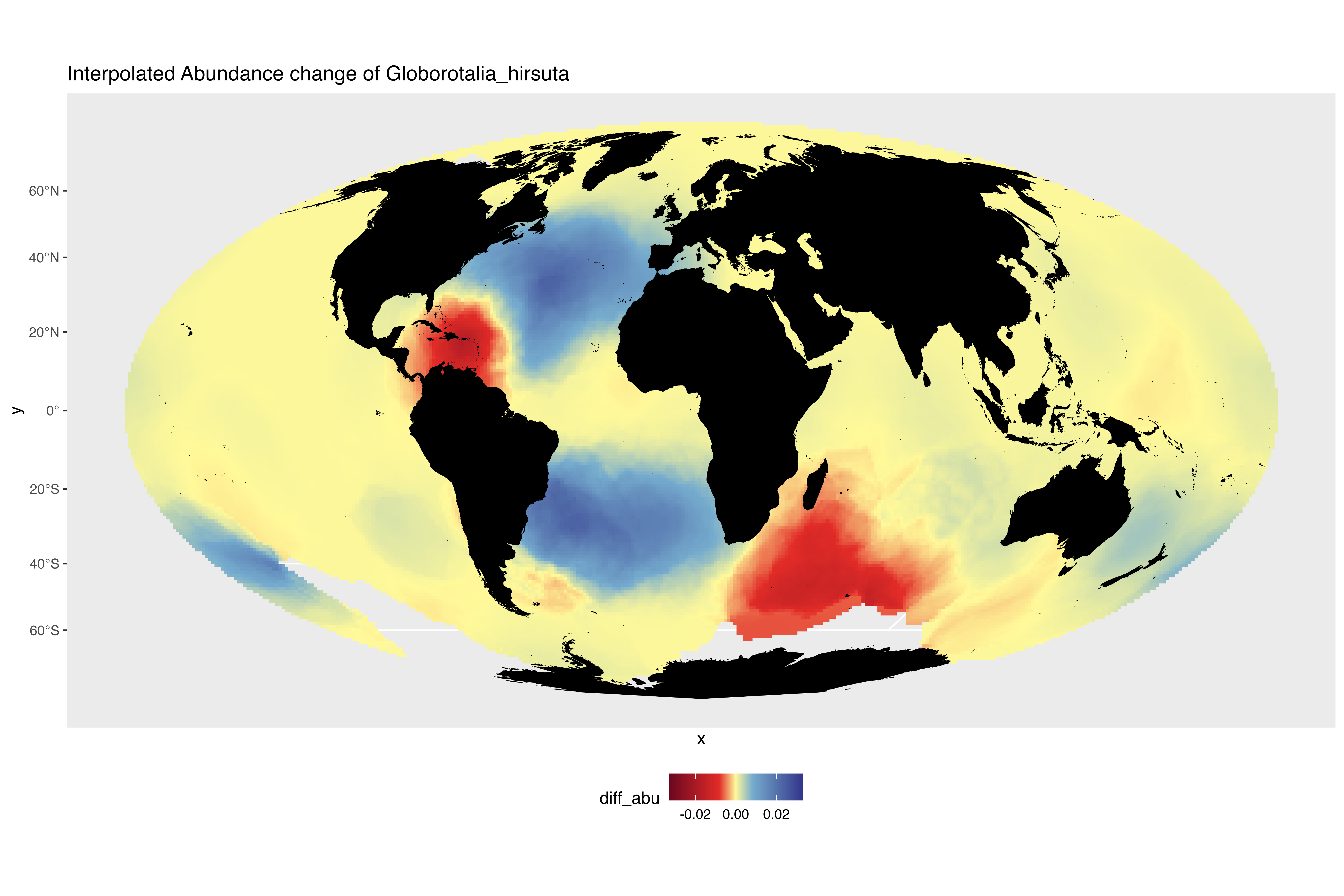

### interpolated_abundance_change_Globorotalia_menardii.png

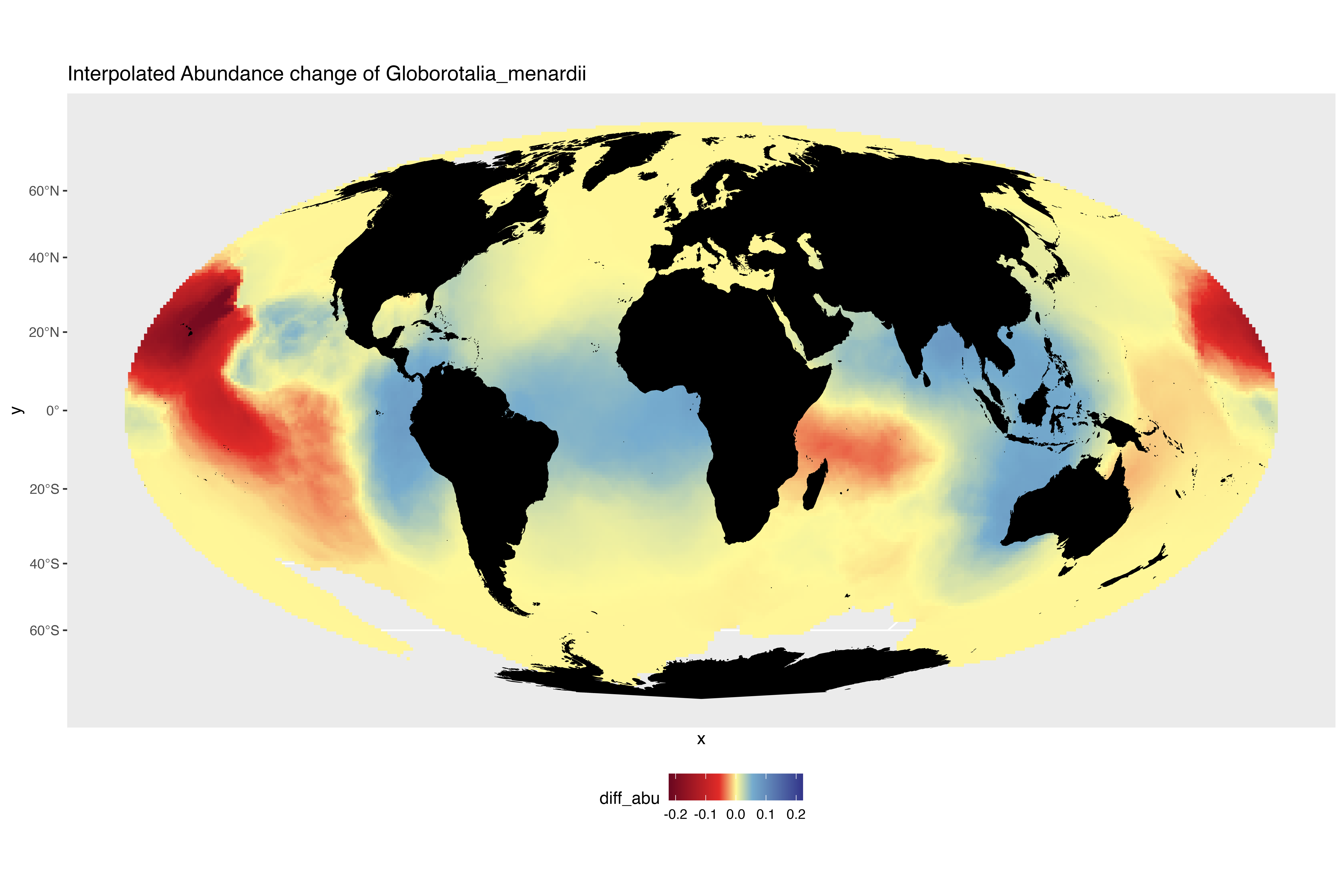

### interpolated_abundance_change_Globorotalia_scitula.png

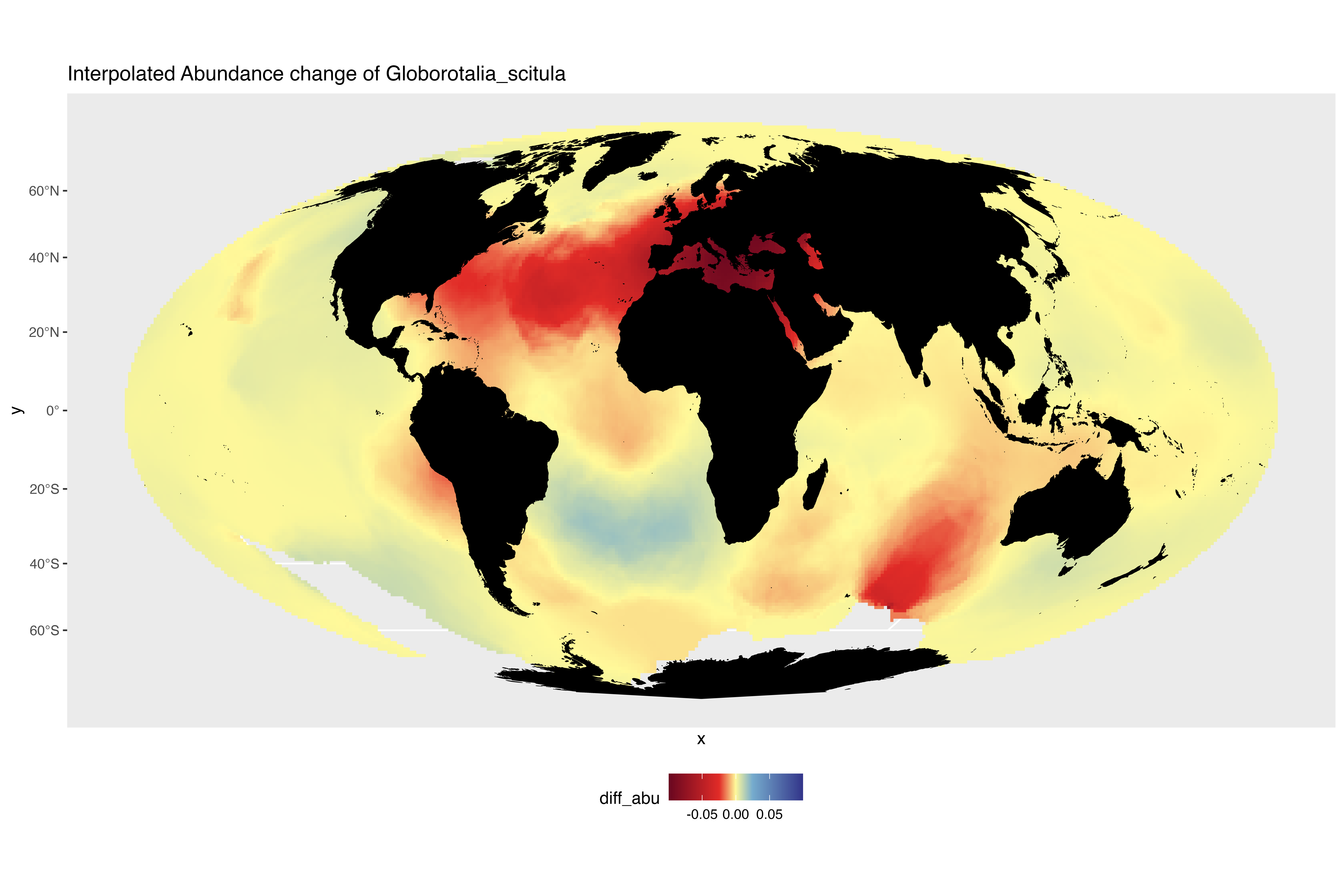

### interpolated_abundance_change_Globorotalia_theyeri.png

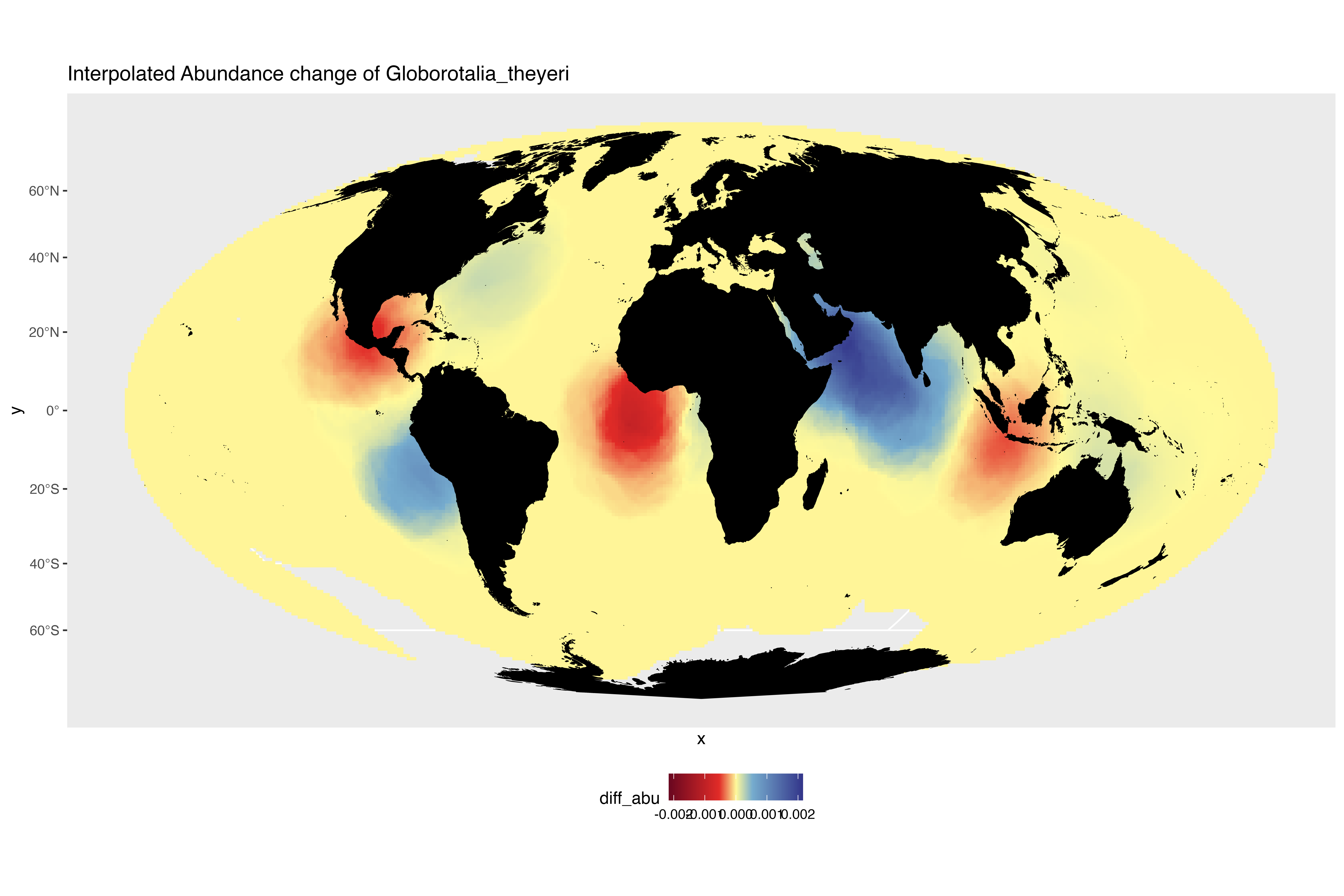

### interpolated_abundance_change_Globorotalia_truncatulinoides.png

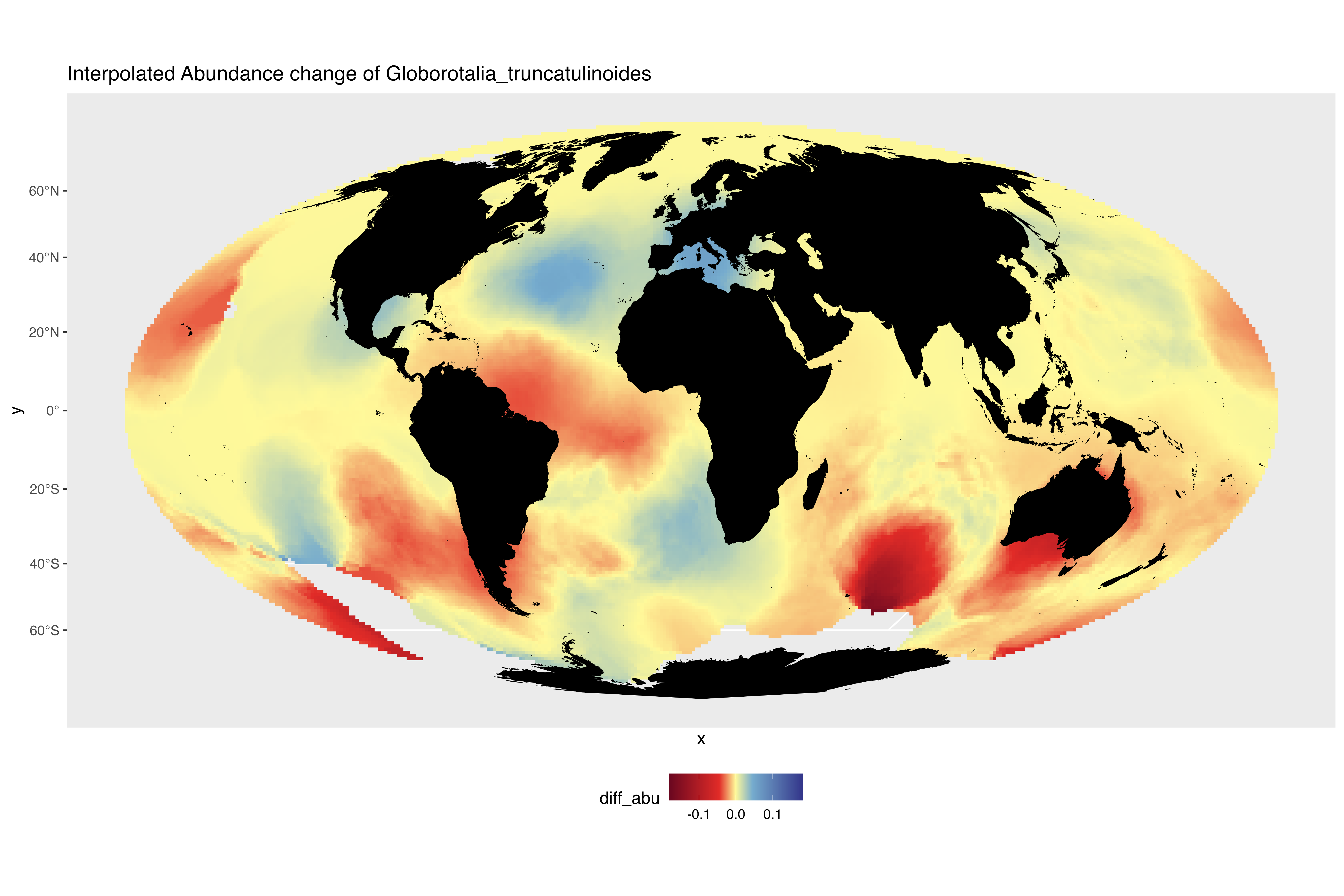

### interpolated_abundance_change_Globorotalia_tumida.png

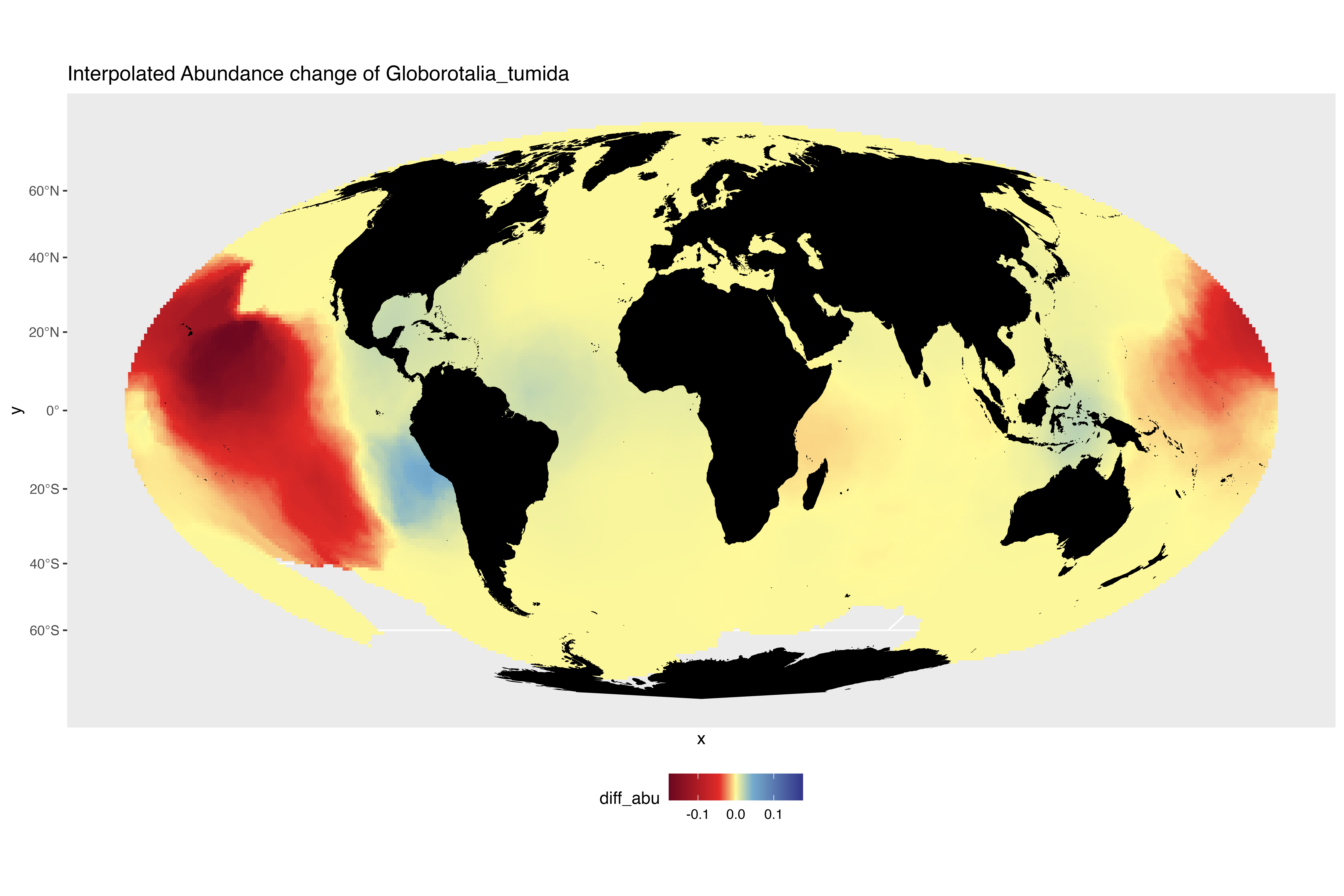

### interpolated_abundance_change_Globorotalia_ungulata.png

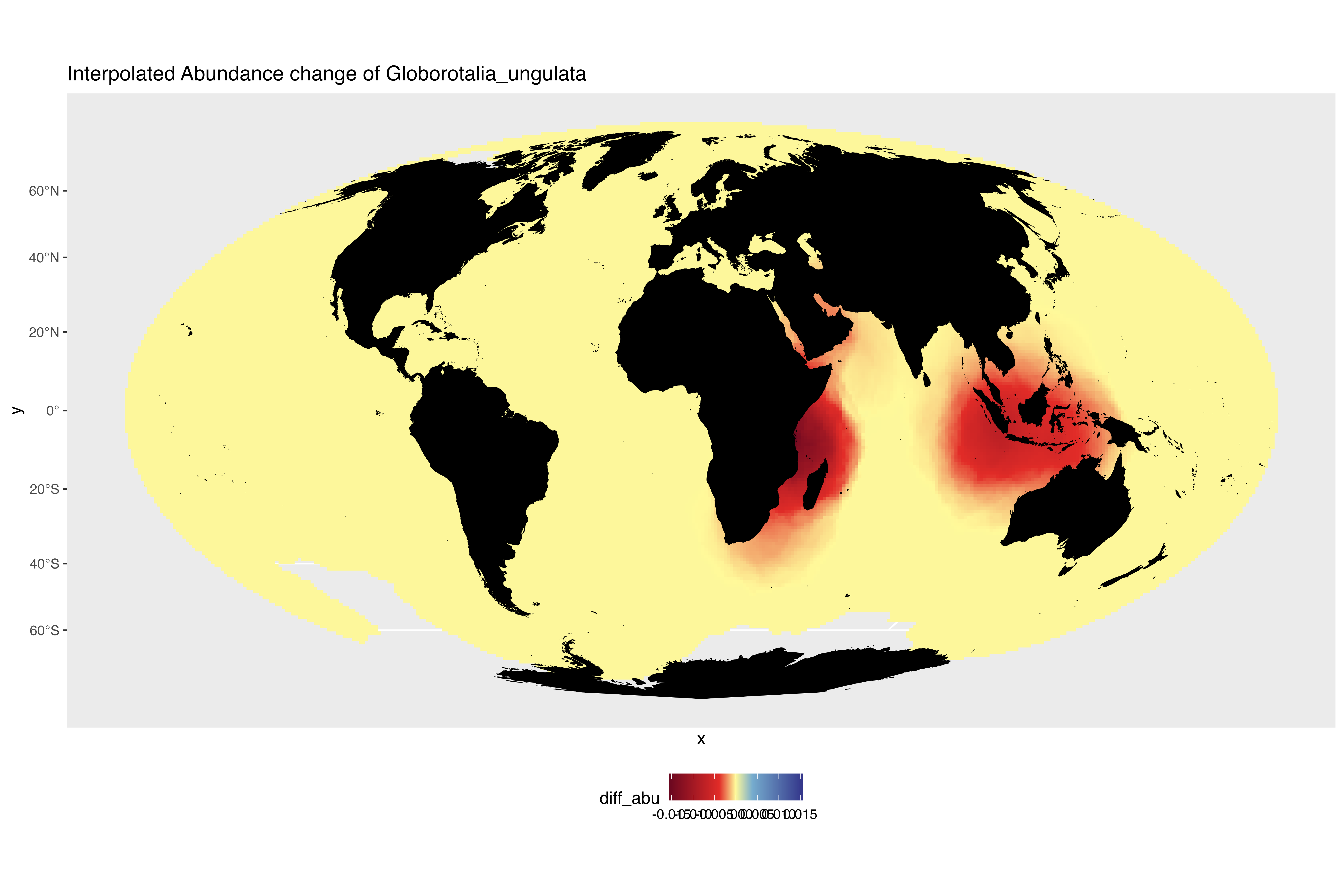

### interpolated_abundance_change_Globorotaloides_hexagonus.png

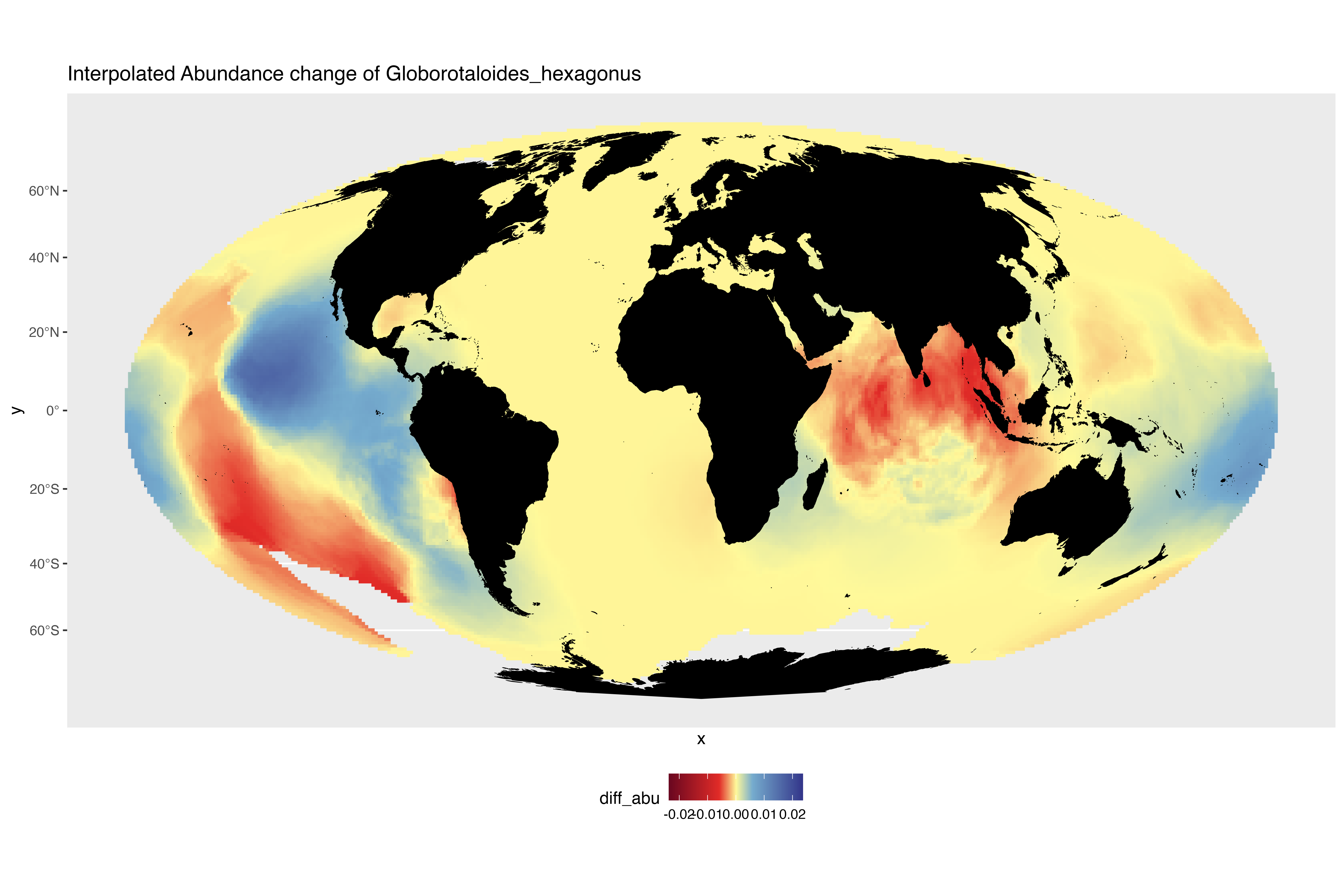

### interpolated_abundance_change_Globoturborotalita_rubescens.png

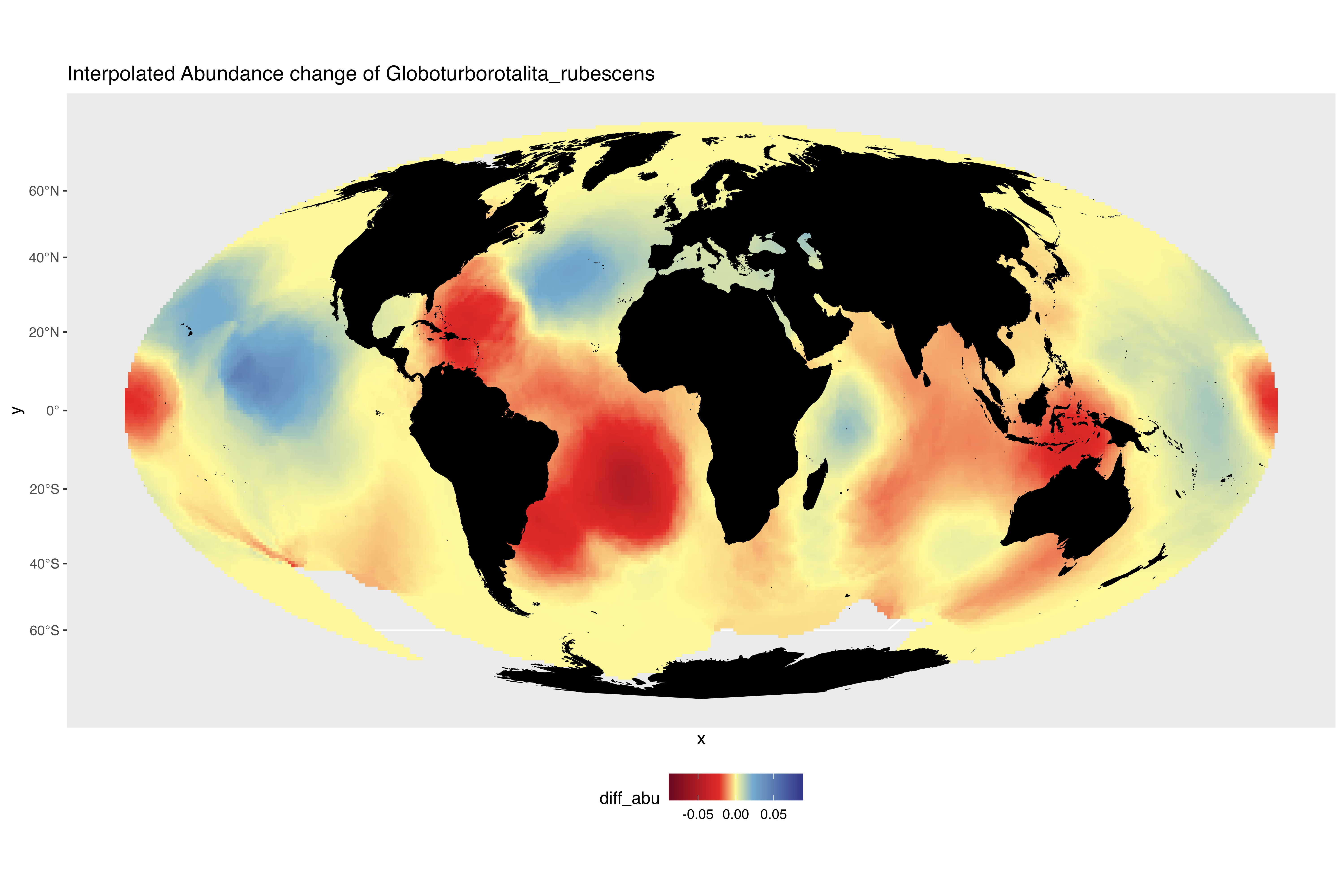

### interpolated_abundance_change_Neogloboquadrina_dutertrei.png

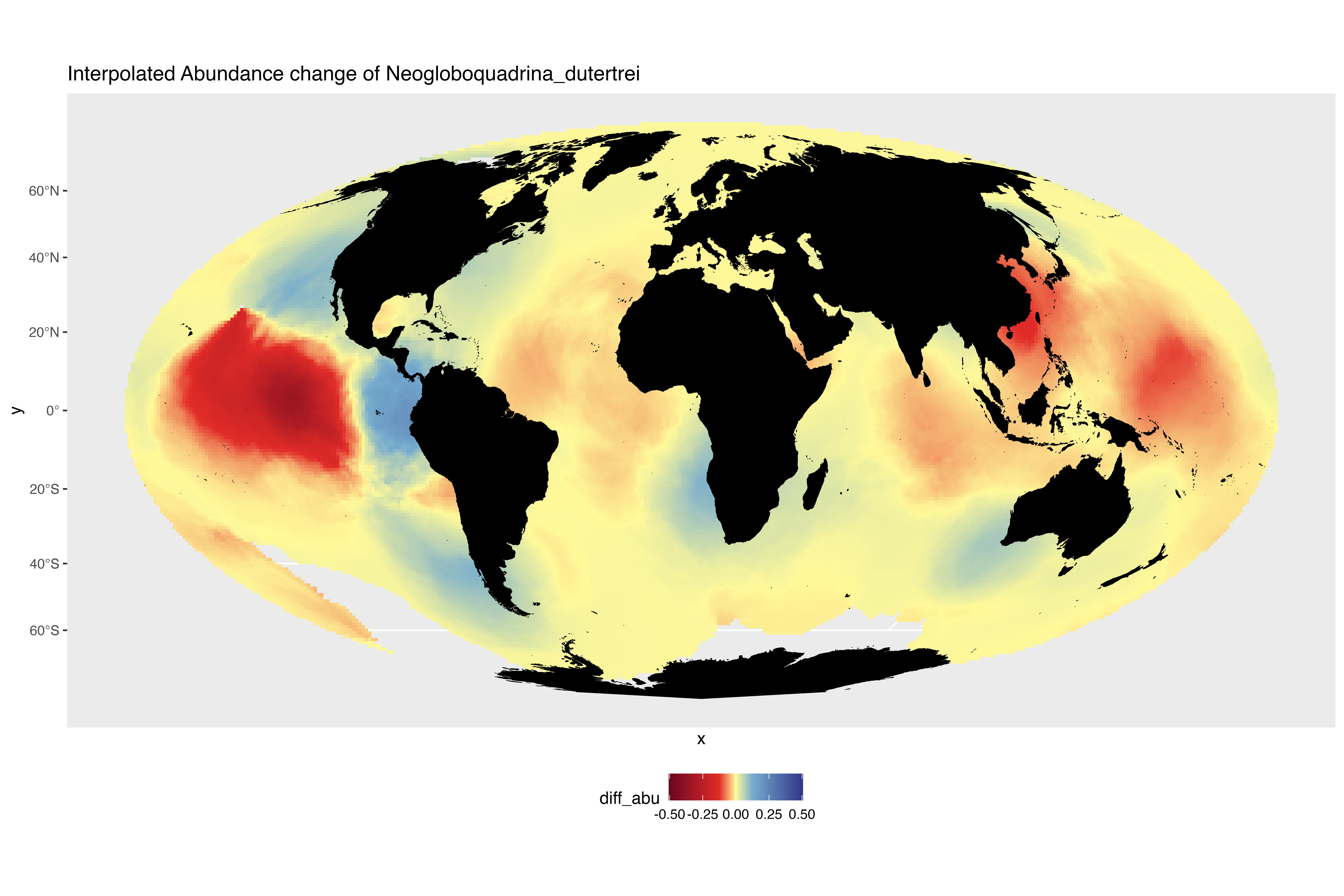

### interpolated_abundance_change_Neogloboquadrina_incompta.png

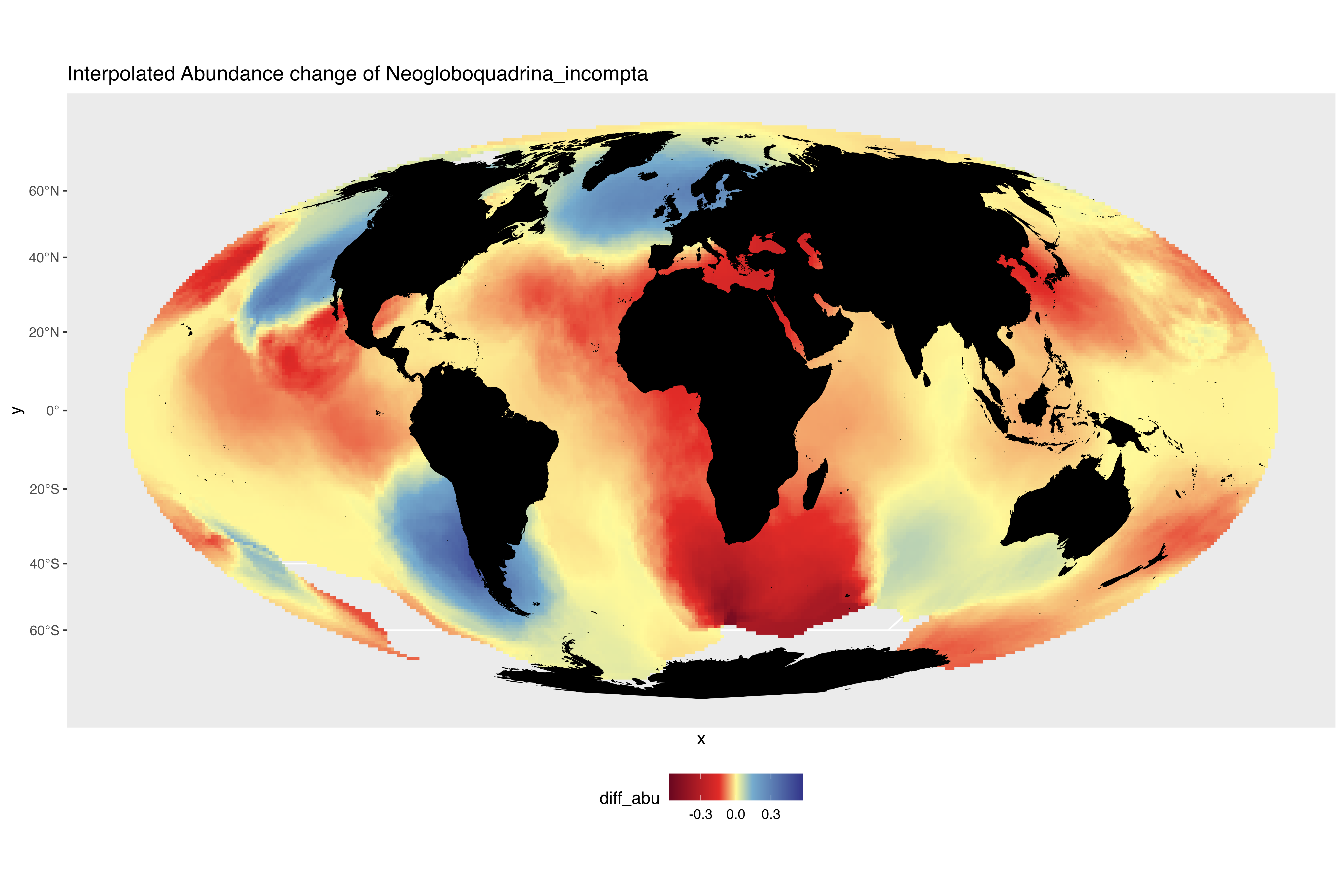

### interpolated_abundance_change_Neogloboquadrina_pachyderma.png

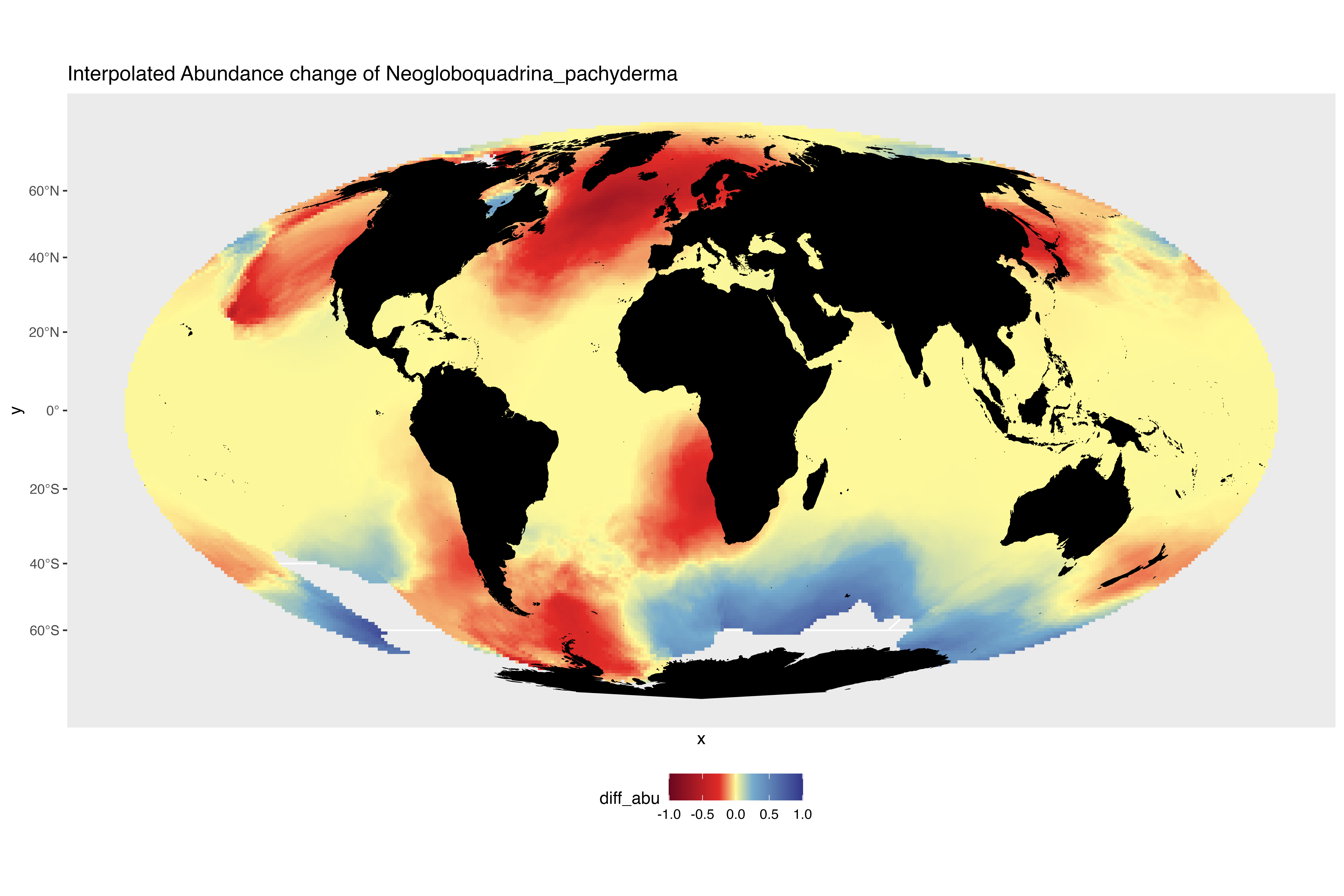

### interpolated_abundance_change_Orbulina_universa.png

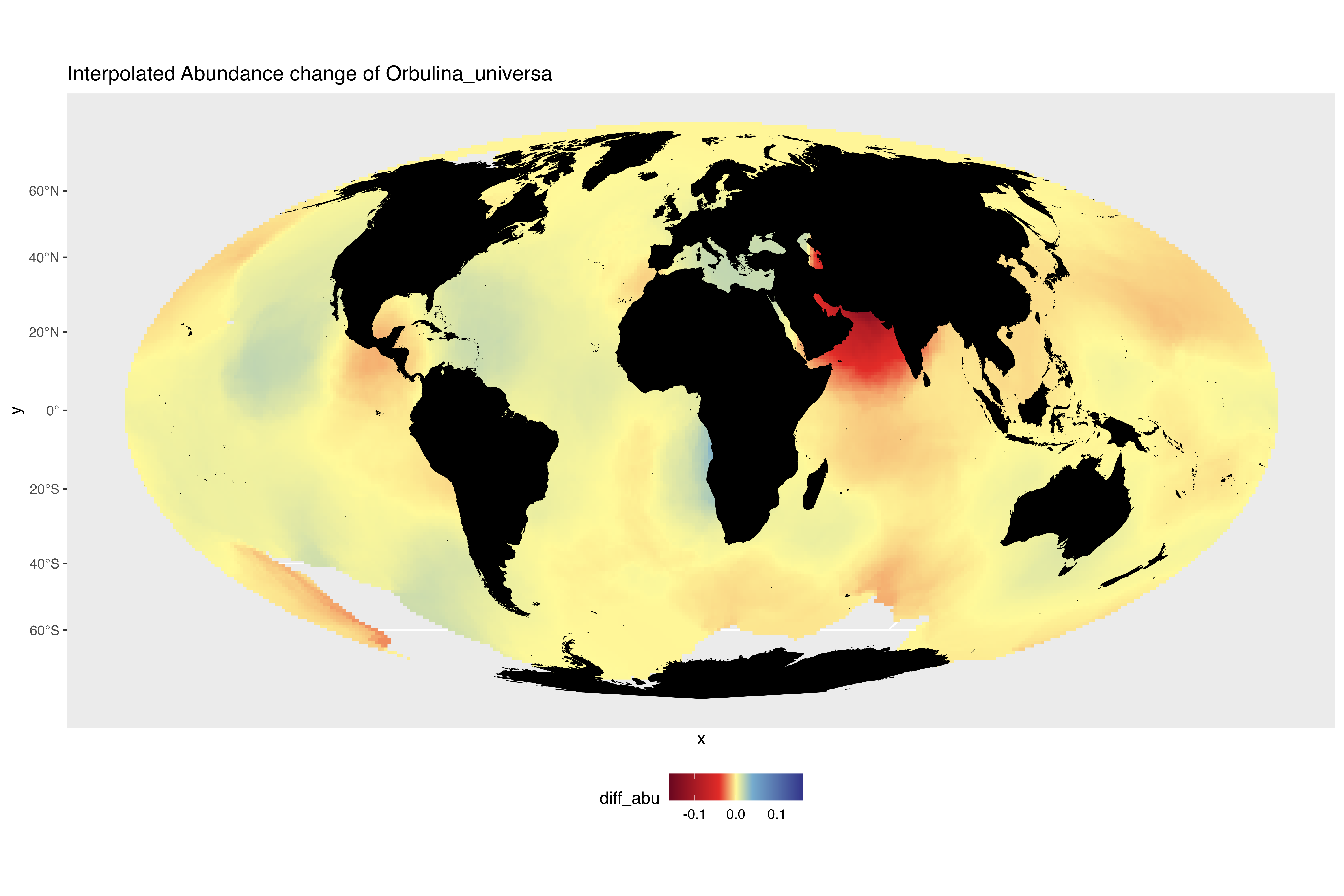

### interpolated_abundance_change_Pulleniatina_obliquiloculata.png

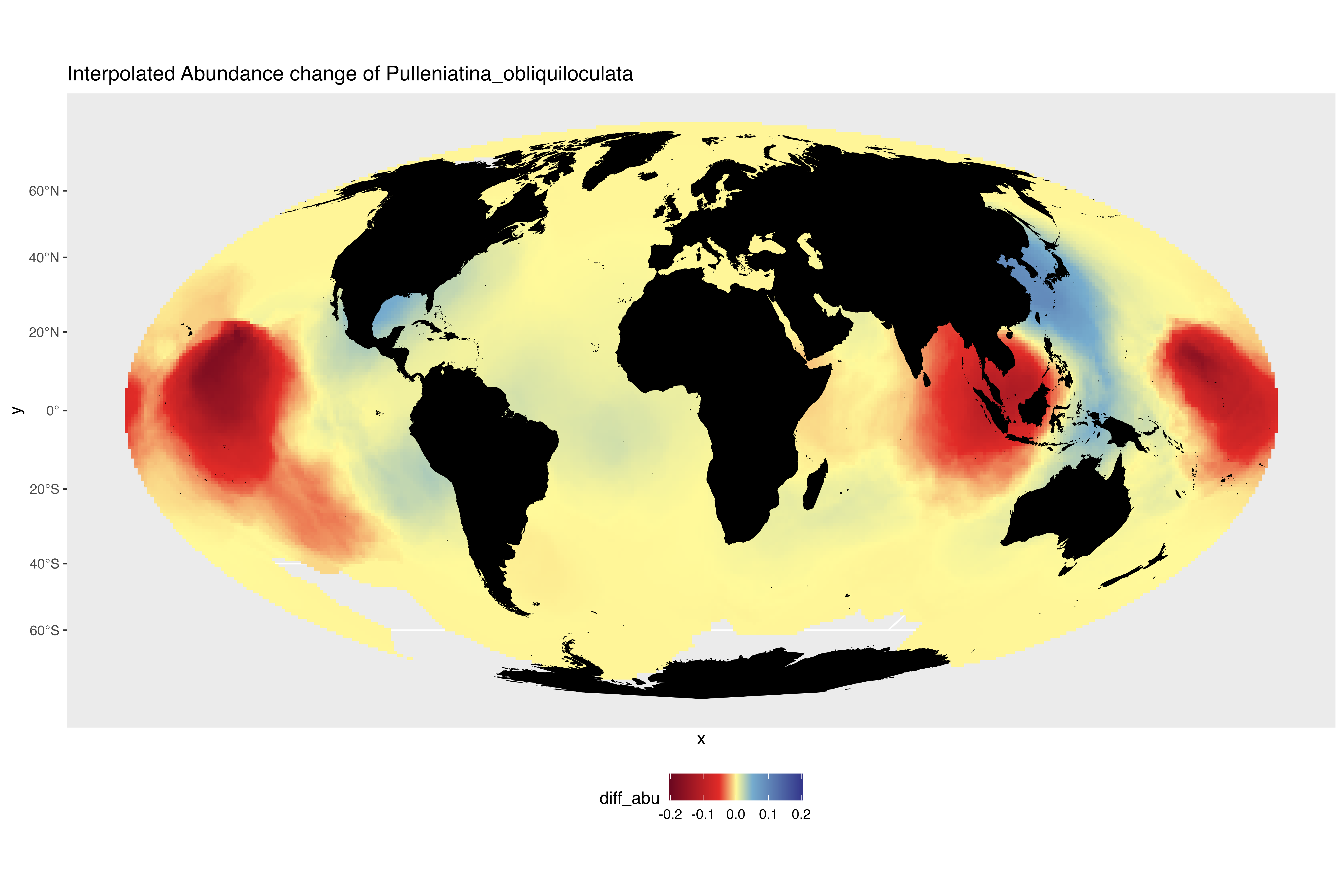

### interpolated_abundance_change_Sphaeroidinella_dehiscens.png

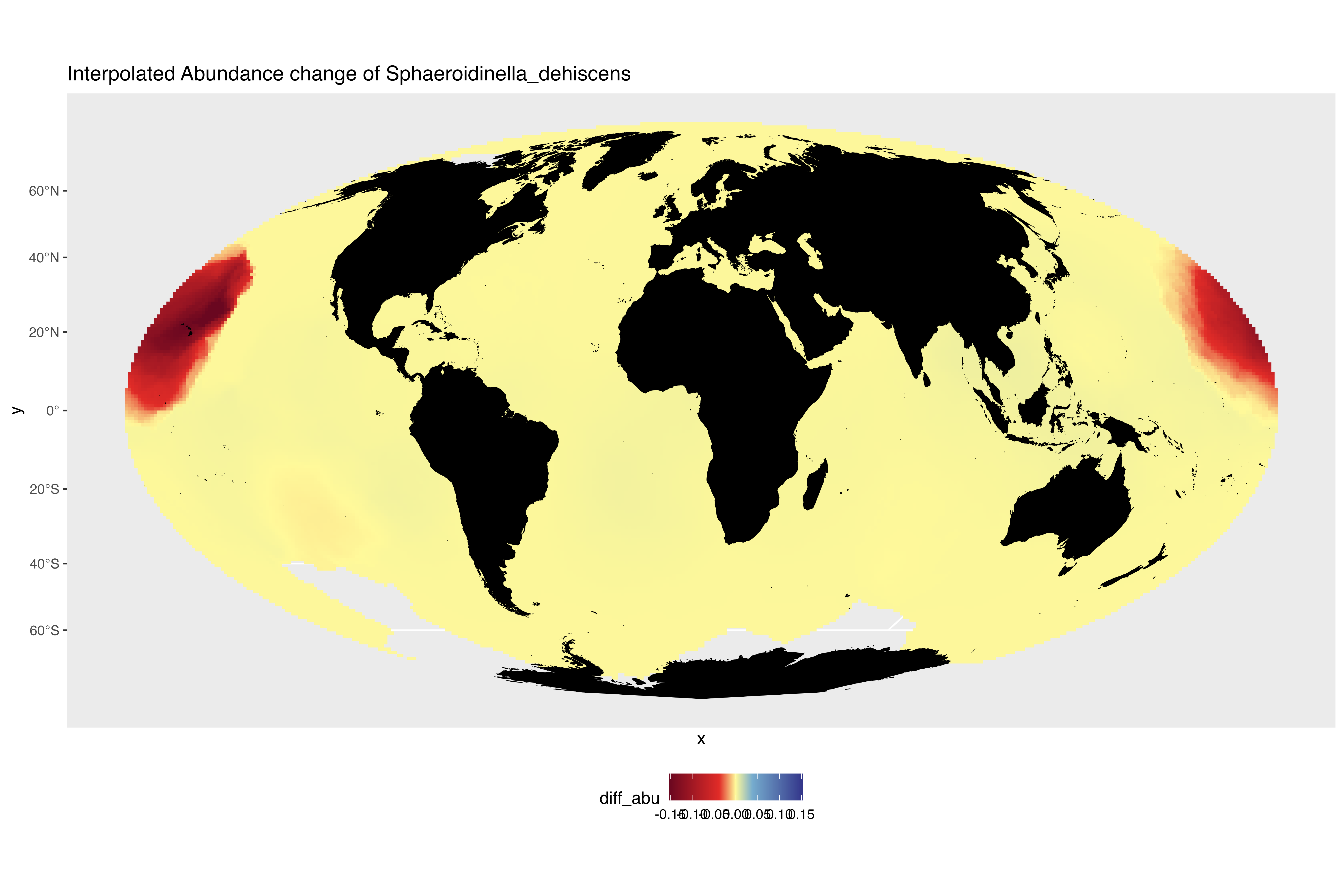

### interpolated_abundance_change_Tenuitella_iota.png

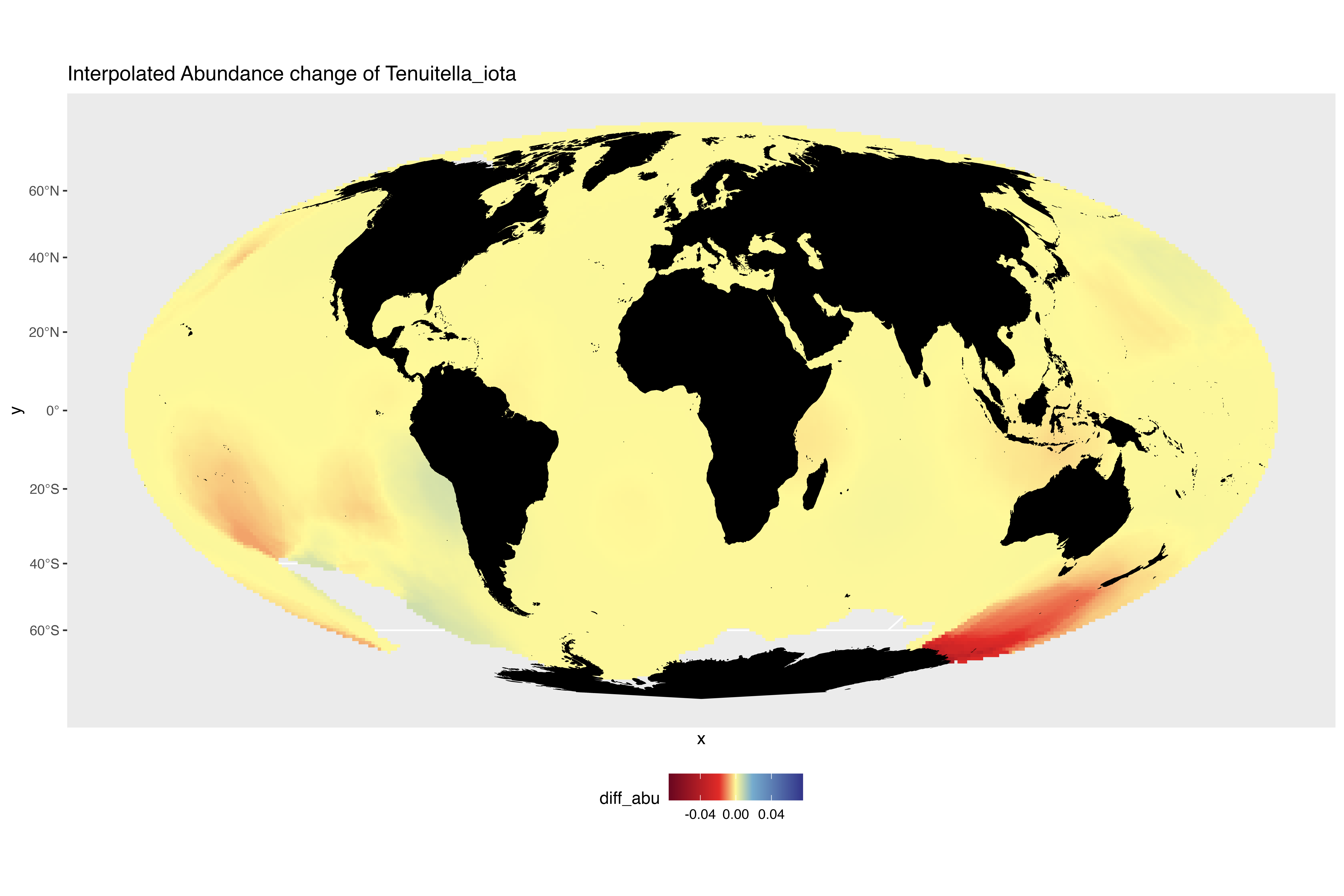

### interpolated_abundance_change_Trilobatus_sacculifer.png

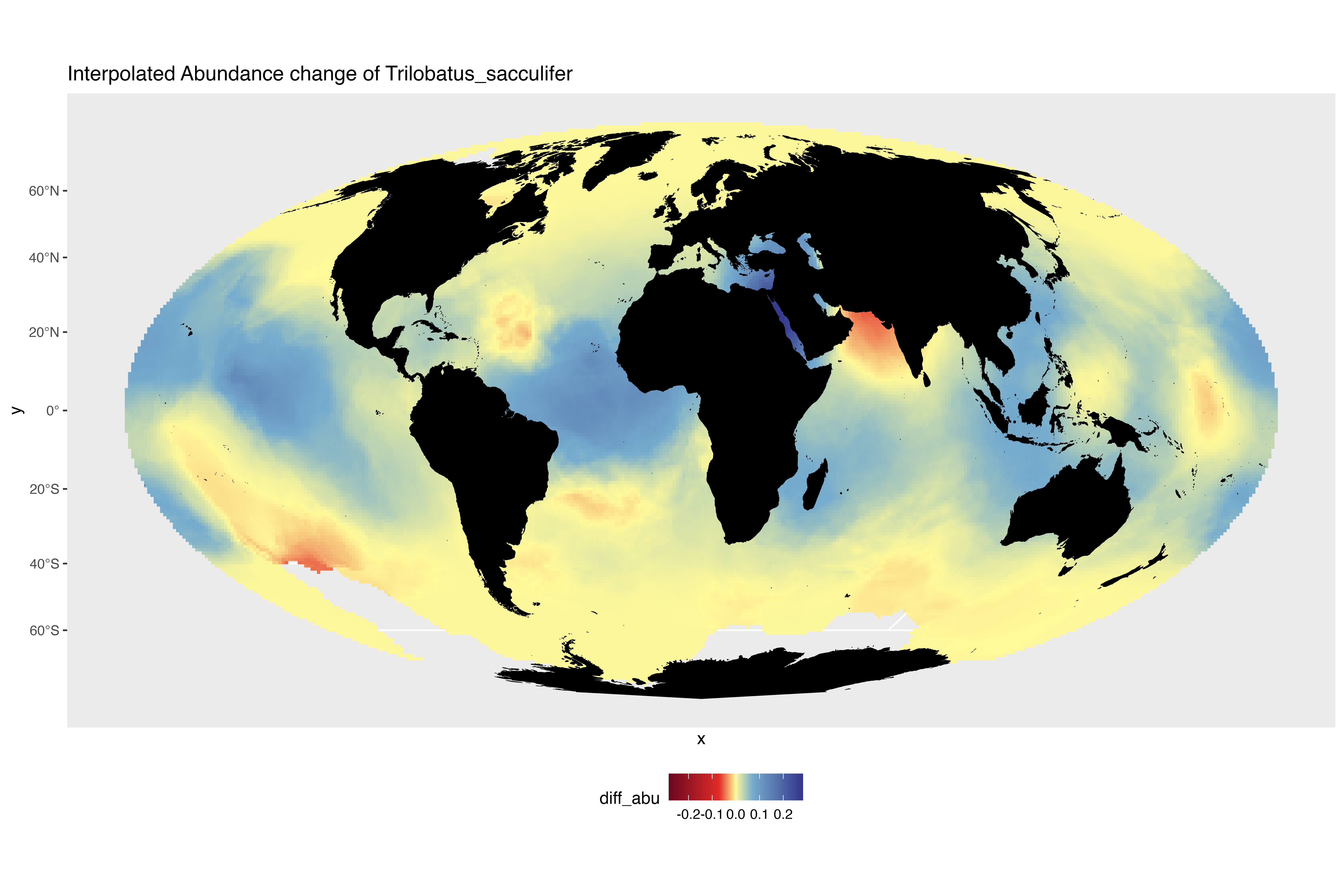

### interpolated_abundance_change_Turborotalita_humilis.png

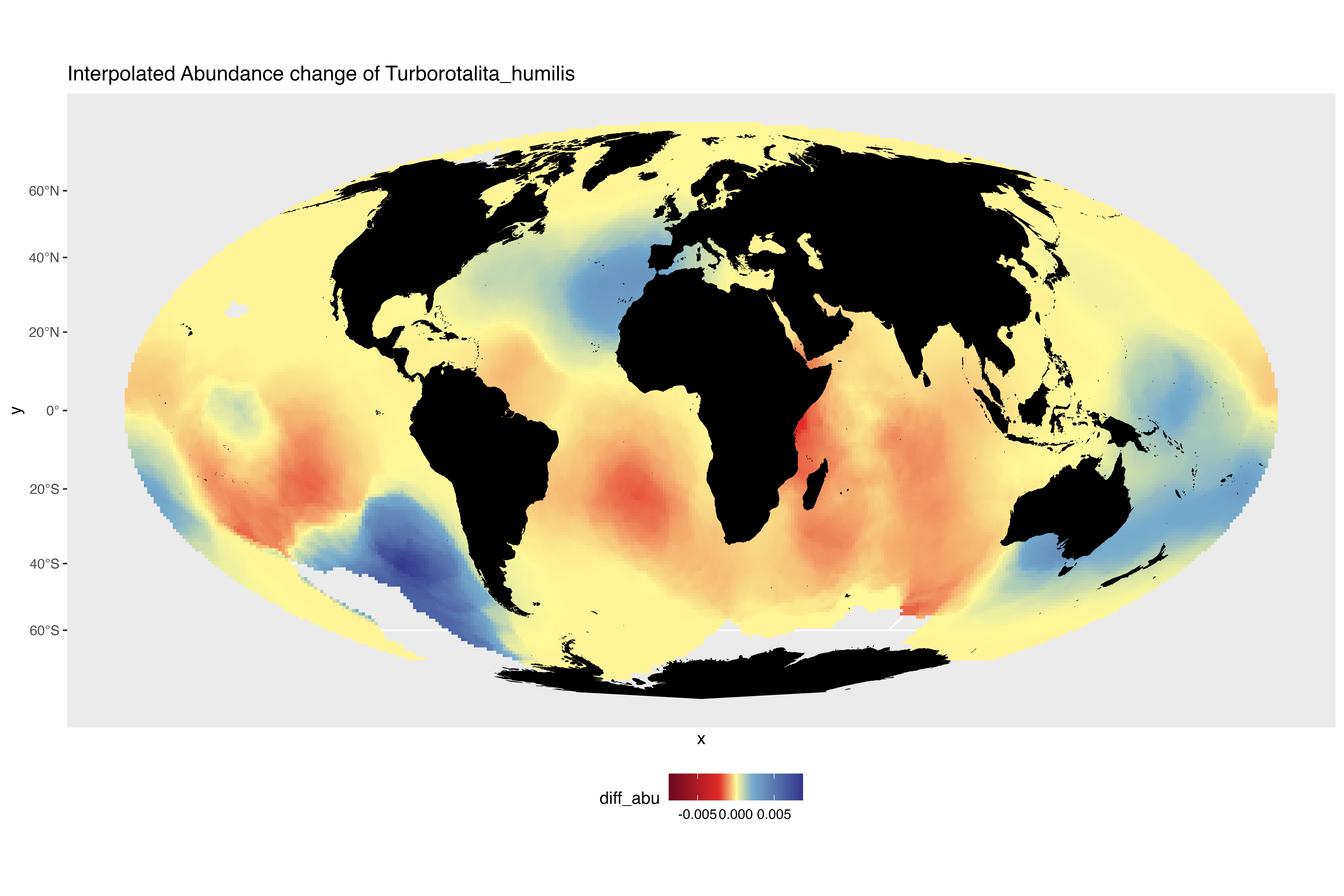

### interpolated_abundance_change_Turborotalita_quinqueloba.png

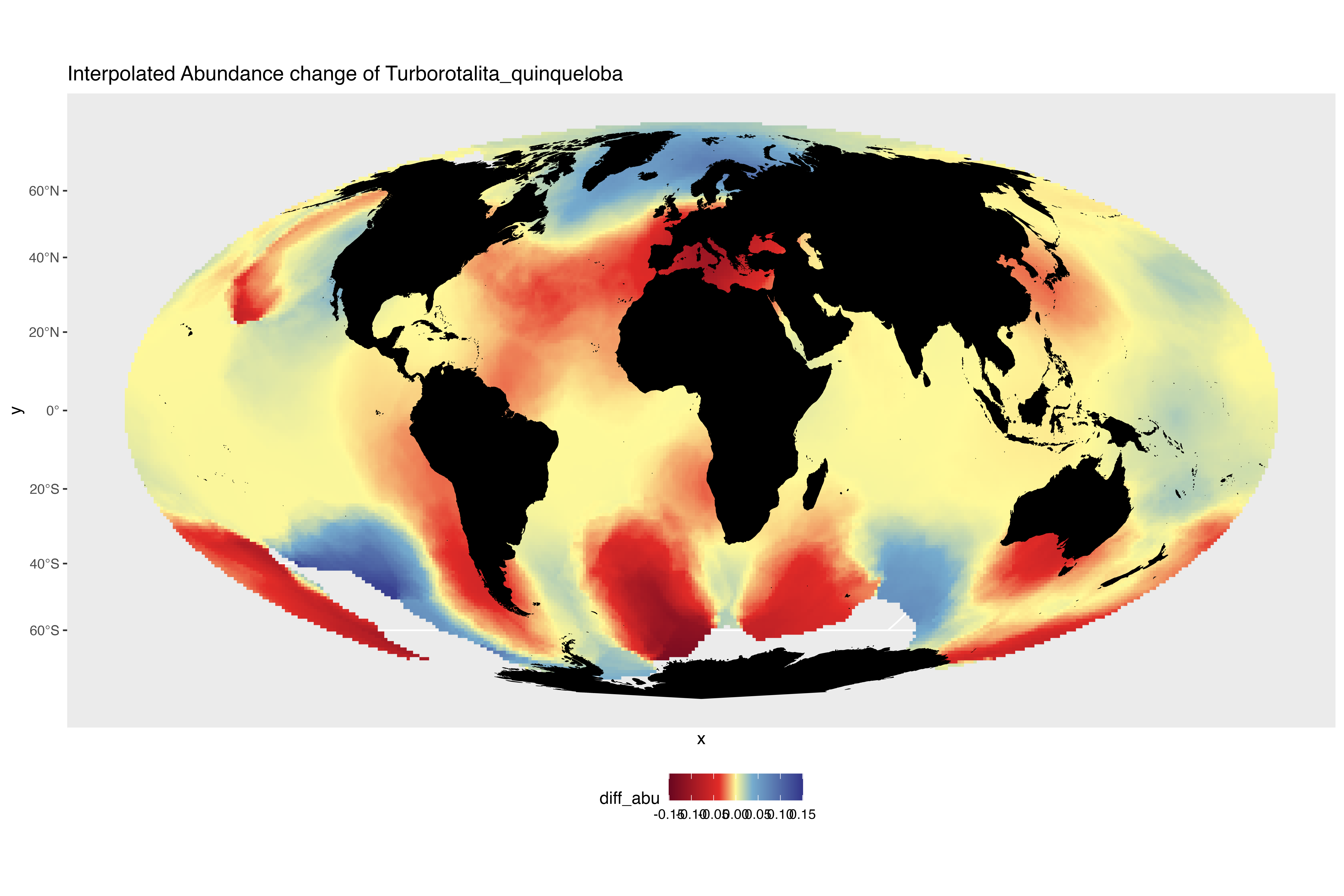
